# Engineered mRNA nanostructures expand the design space of mRNA therapeutics through programmable protein expression and immune stimulation

**DOI:** 10.64898/2026.09.14.751315

**Authors:** Mark D. Knappenberger, Skylar J.W. Henry, Archana Kikla, Matthew Sample, Olivia Roy, Erik Poppleton, Nicholas Stephanopoulos, Michael P. Gustafson, Karen S. Anderson, Petr Šulc

**Author notes:** these authors contributed equally (Mark D. Knappenberger, Skylar J.W. Henry, Archana Kikla).

## Abstract

Messenger RNA (mRNA) therapeutics have transformed vaccination and protein replacement strategies, yet efforts to improve their performance have focused largely on sequence engineering, nucleotide modification, and delivery vehicles. Here we show that mRNA function can be controlled by rational design of higher-order RNA architectures. We develop self-assembling mRNA origami (mRNA-OG), a class of unimolecular RNA nanostructures that encode protein-coding sequences within higher-order, programmable, and compact nucleic acid architectures. Using computational design and experimental validation, we demonstrate that mRNA-OG folds into well-defined nanostructures while remaining translationally competent in mammalian cells. Although folded mRNA-OG recruits ribosomes comparably to unfolded constructs, it produces lower protein output, indicating that RNA architecture can directly influence translational efficiency. The compact geometry of mRNA-OG also enhances encapsulation by cationic lipid delivery systems, suggesting a structural route to improved cargo packaging. Beyond its effects on translation, mRNA architecture modulates innate immune recognition. In primary human dendritic cells, folded and unfolded mRNA-OG elicit distinct cytokine programs and differential activation of stress-response pathways, including a modest induction of the integrated stress response that is not fully explained by canonical PKR signaling. Our results establish programmable structure as a new design parameter for mRNA therapeutics that can affect its functionality, delivery properties, and immune sensing.

## Introduction

Messenger RNA (mRNA) has emerged as a promising class of therapeutic agent, with applications in vaccination, immunotherapy, and gene/protein therapy. Advances in mRNA engineering such as chemical modification, sequence optimization, and scalable purification methods, have accelerated its clinical translation^1,2^. Nevertheless, like other nucleic acid-based therapeutics, mRNA requires a carrier to facilitate efficient cellular delivery and uptake. Lipid nanoparticles (LNPs) represent the most widely adopted carrier platform to date, underpinning the FDA-approved Moderna and Pfizer-BioNTech COVID-19 mRNA vaccines. Despite their clinical success, LNPs are associated with several limitations: low encapsulation efficiency (averaging approximately two mRNA molecules per particle^3^), requirement for cold-chain storage, suboptimal endosomal escape and cytoplasmic delivery, limited colloidal stability and polydispersity, and a formulation complexity that necessitates re-optimization for each target mRNA. Collectively, these constraints highlight the need for improved mRNA stabilization and delivery strategies, as well as the need to seek potential alternatives to LNPs.

Another roadblock to the delivery of exogenous mRNA to cells is immune stimulation due to the molecule’s recognition by endosomal TLR3, TLR7, and TLR8, as well as cytosolic MDA5 and RIG-I (see full list of abbreviations and definitions in Supplementary Tables S1, S2). Recognition by these receptors results in the release of type I interferons (IFNs) that activate the JAK-STAT pathway and trigger the expression of IFN-related genes. These genes include protein kinase R (PKR), the activation of which has been shown to suppress translation of exogenous mRNA through phosphorylation of eIF2α, though the magnitude of this effect is highly context- and structure-dependent^4^. Thus, chemical modifications of the mRNA molecule have been developed to dampen the immunogenicity and improve its stability and translation efficiency. One such modification, implemented in the COVID-19 mRNA vaccines, is the incorporation of N1-methylpseudouridine (m^1^Ψ), which decreases PKR activation and thus enhances translation of the mRNA. However, incorporation of m^1^Ψ has been shown to cause ribosomal slippage at the translation start site, creating frameshifted proteins that elicit unintended immune responses^5^.

Nucleic acid nanotechnology offers a complementary approach to these challenges, one that leverages programmed three-dimensional architecture instead of, or in addition to, chemical modification to achieve stability, controlled immune engagement, and efficient delivery. Over the past four decades, the field of nucleic acid nanotechnology has established design rules for 2D and 3D self-assembled programmable nanostructures built out of DNA or RNA. The most frequently used design paradigm, DNA origami^6^, consists of a long DNA scaffold strand and a large number (∼200) of short staple strands which significantly increase synthesis cost and annealing complexity. Each staple strand requires chemical synthesis, and 100% incorporation of staples onto the DNA scaffold is challenging to accomplish, in turn making it difficult to achieve reliability and scalability with multi-strand nanostructures. Several DNA-origami inspired approaches have also been developed for hybrid DNA-RNA structures, where mRNA was used as a scaffold with DNA staples designed to create a 3D shape^7^, and recently a DNA:RNA hybrid origami with encoded GFP was shown to express in cells as well^8^. In parallel, several recent efforts in mRNA therapeutics have focused on redesigning mRNA synonymous codons in order to increase its number of base pairs in secondary structure^9,10^. These “superfolder mRNAs” were shown to reduce hydrolytic degradation rates by half, by minimizing the fraction of unpaired nucleotides available for participation in in-line hydrolysis^11^. However, choosing different codons can affect translation, and the constraint to keep the amino acid sequences the same still restricts the amount of intramolecular association that can be achieved, as it is impossible to always achieve base pairing between different coding segments.

Here, we leverage a new nucleic nanotechnology design strategy to create compact mRNA folded structures with maximized base pairing. Our approach is based on single-stranded RNA origami (ssRNA-OG) nanostructures. These designs consist of a single long RNA strand that folds into a compact 2D or 3D shape. Two main approaches for ssRNA-OG nanostructures have been proposed. The first relies on kissing loop motifs to position helices into a given target shape^12,13^, which enables cotranscriptionally folded structures. Applications of this design include artificial cytoskeletons in synthetic cell applications^14^, or for nucleic acid sensors^15^. The second ssRNA-OG design approach, which we chose to adapt here, employs alternating duplex regions and paranemic (PX) cross-over motifs^16,17^ to position adjacent helices, resulting in compact, topologically-unknotted structures (Fig. 1a). These ssRNA-OG have demonstrated remarkable stability in serum for up to 16 hours in the absence of a carrier or chemical modifications, a property hypothesized to arise from the compact architecture limiting RNase accessibility^18,19^. The same compact architecture may have beneficial effects in promoting encapsulation efficiency in cationic polymer-based delivery methods, as well as scaffolding cationic peptide-based coatings such as oligolysine^20^. Previous studies have shown ssRNA-OG nanostructures to be potent immunomodulatory agents: stimulation of TLR3 by ssRNA-OG, without induction of high levels of type I IFNs, enabled its use as an immunotherapeutic agent against cancer, inducing significant tumor retardation or regression through activation of NK cell– and CD8⁺ T cell–dependent antitumor immunity and antagonism of the peritoneal immunosuppressive environment^18,21^. ssRNA-OG constructs covalently linked to antigenic peptides have similarly demonstrated efficacy in engaging dendritic cells (DCs) for maturation and peptide cross-presentation to CD8⁺ T cells^21,22^. Notably, nucleotide modification of ssRNA-OG can modulate its immune reactogenicity in a manner analogous to conventional mRNA modifications without impairing its ability to fold into predefined architectures, establishing that the structural and immunological properties of ssRNA-OG can be tuned independently. Previous studies with compact DNA nanostructures showed that they can be efficiently coated with endosome escape peptides for cell delivery^23–25^, and similar techniques for coating compacted RNA nanostructures can thus offer an alternative to LNP delivery for mRNA therapeutics as well.

**Fig. 1.**
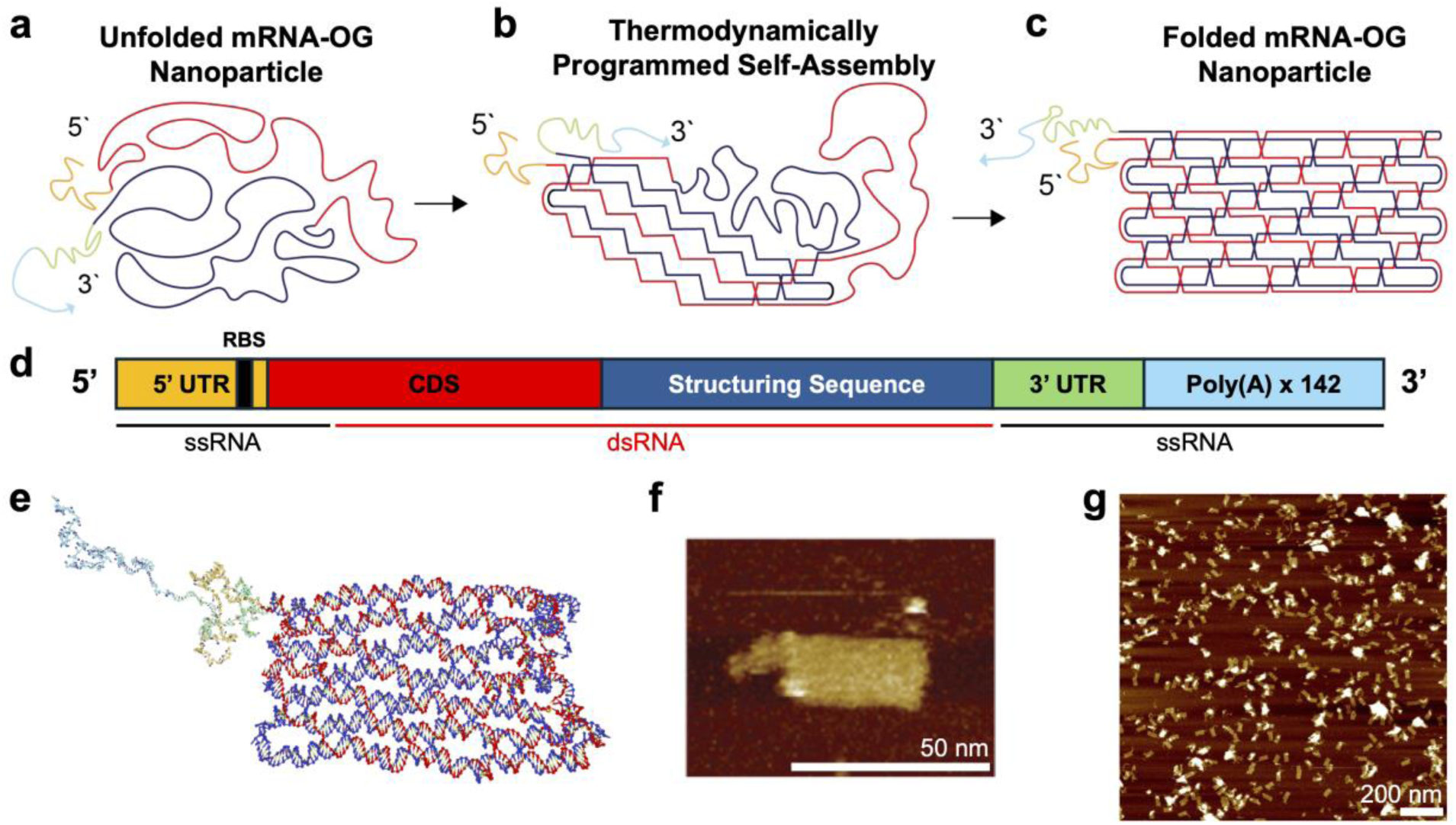
Single-stranded mRNA origami (mRNA-OG) nanoparticle design and structural characterization. **a-c,** Schematic representation of the thermodynamically programmed self-assembly of the mRNA-OG into a predefined structure. **d,** Linear sequence schematic of the mRNA-OG. **e,** Predicted oxRNA structure of the mRNA-OG. **f,** Close-up atomic force microscopy (AFM) image of a mRNA-OG nanoparticle. **g,** Wide-field AFM image of the mRNA-OG.

Given these promising properties of ssRNA origami nanostructures, we explored adapting the paranemic-crossover based designs as an mRNA delivery platform. The incorporation of a translatable open reading frame directly into a ssRNA origami architecture without staple oligonucleotides, chemical crosslinks, or cotranscriptional assembly partners unites structural programmability, cargo encoding, and immunomodulatory properties in a single construct, while enabling thermodynamic self-assembly from sequence alone. Moreover, the single-stranded nature of the particle enables high-yield production with *in vitro* transcription and straightforward purification, simplifying manufacturing and scalability relative to multi-component systems.

First, we develop the mRNA-Design computational pipeline that for any given coding sequence designs a structuring region, which is appended after the coding region to ensure folding into a compact shape. This design paradigm requires no engineering of the mRNA sequence and thus overcomes any codon-optimization limitations. As such, the mRNA coding sequence can be codon-optimized independent of the structure. This is possible because majority of the base-pairs in the PX ssRNA-OG designs are between the first half and second half of the origami sequence and therefore can achieve around 90% designed base-pairs for arbitrary mRNA sequences with an origami length twice the length of the mRNA sequence.

We use the pipeline for a proof-of-principle single-stranded mRNA-OG design encoding EGFP, where 90.3% of the structure is designed to be fully base-paired. Structural characterization by atomic force microscopy and electrophoretic mobility analysis confirmed efficient folding and formation of monodisperse nanoparticles. Importantly, we verified that the resulting nanostructures remained capable of protein expression in HEK293T cells and primary human dendritic cells, demonstrating that complex RNA architectures can support translation despite extensive intramolecular base pairing. We further validated mRNA-OG expression using polysome profiling, where we observe multiple ribosomes are binding to the sequence, confirming the expression of the highly structured mRNA.

A viable delivery platform must occupy a therapeutic window where innate immune sensing is strong enough to be self-adjuvating yet below the threshold at which it arrests translation of its own payload or kills the expressing cell. Exogenous mRNA is surveilled by innate pattern-recognition receptors (PRRs) that distinguish non-self nucleic acids. Endosomal Toll-like receptors (TLR3, TLR7/8) sample internalized RNA, while the cytosolic sensors RIG-I, MDA5, and protein kinase R (PKR) detect RNA reaching the cytoplasm. These sensors converge on two outputs that matter for a structured-mRNA platform. The first is type I interferon induction, which can act as a built-in adjuvant but becomes cytotoxic at high or sustained levels. The second is the integrated stress response (ISR), in which eIF2α phosphorylation attenuates cap-dependent translation and, if severe, triggers CHOP-dependent apoptosis. Therefore, a middle-ground must be achieved where there is enough immune activation to elicit an immune response, but not so high that it is cytotoxic. mRNA-OG is by definition comprised of extensive intramolecular duplex formation, but also carries a Cap 1 (2′-O-methylated) 5′ end on an unstructured UTR (a configuration that limits the 5′-end discrimination used by RIG-I), though its PX-crossover-interrupted helices may not satisfy MDA5’s requirement for long continuous duplex. As such, we hypothesized that PKR, which responds to short internal duplexes independent of 5′-cap chemistry, would be the dominant cytosolic sensor of the folded nanoparticle. We compare the folded mRNA-OG with its unannealed, misfolded variant. We observed that the two variants generated distinct cytokine signatures and patterns of innate immune activation, accompanied by modest eIF2α phosphorylation and stress-response signaling that remained below levels associated with cytotoxic integrated stress response activation.

Finally, we focused on properties that are relevant to potential therapeutic applications. Folded nanostructures displayed markedly improved encapsulation efficiency with cationic lipid delivery reagents compared with linear mRNA and retained structural integrity and expression competence during prolonged refrigerated storage. In primary human dendritic cells, mRNA-OG constructs promoted maturation-associated phenotypes while preserving co-stimulatory signaling in neighboring bystander cells, suggesting that programmable RNA architecture may influence both direct and indirect immune responses.

Together, these results establish single-stranded mRNA-OG as a new class of programmable RNA therapeutic scaffold that combines protein-coding capacity with nanoscale structural control and expands the design space of mRNA therapeutics beyond sequence and chemical composition alone. Our work introduces designed RNA structure as an additional programmable parameter for future mRNA therapeutic applications such as vaccine delivery, immunotherapeutics, antibody-nanoparticle conjugates, or targeted gene delivery.

## Results & Discussion

### Design, Production, and Structural Characterization of mRNA-OG

To utilize ssRNA origami as a versatile platform for mRNA delivery, it is essential to seamlessly integrate any gene of interest into the nanostructure with minimal disruption to its programmed architecture. To achieve this, we developed an automated computational design program named mRNA-Design that embeds arbitrary mRNA coding sequences into ssRNA origami topologies. Building upon the paranemic cohesion ssRNA origami design paradigm established by Han et al.^16^, the mRNA-Design algorithm takes a predesigned paranemic ssRNA-OG nanostructure architecture and redesigns the nucleotide sequence to contain a functional mRNA coding sequence while strictly preserving its defining crossover network and global architecture. Here we use a quadrilateral origami shape shown in Fig. 1c as the input structure, as its length is compatible with the EGFP gene that we aim to encode here, but it can take any paranemic ssRNA origami structure as input, including larger designs (Supplementary Fig. S3). The current design can encode an mRNA sequence length of up to one-half the length of the total ssRNA origami. Larger mRNA sequences can be accommodated with larger ssRNA origami templates or an unstructured 5’ end. The mRNA-Design algorithm partitions the designed ssRNA-OG strand routing into two functional domains: an open reading frame and a structural scaffold (Fig. 1a). From 5’ to 3’, the first half of the sequence contains the coding sequence while the second half is the structuring domain. This physical separation allows the coding sequence to be fixed by the target protein sequence (including any codon optimization desired), while the structuring sequence can be freely assigned to drive nanostructure formation. Given a coding sequence, the algorithm automatically sets the coding-scaffolding domains in the RNA-OG routing, generating the necessary thermodynamic driving force to fold the structure (Fig. 1b-c). Scaffold-scaffold binding domains are also preserved from the original RNA-OG or can be arbitrarily set to minimize sequence conflict with the coding strand. Crucially however, the algorithm cannot structure regions where the RNA-OG strand routing dictates that two segments of the open reading frame domain will base-pair, termed self-binding domains (Supplementary Fig. S1d). As the algorithm redesigns the sequence and not the routing of the ssRNA-OG, local thermodynamic penalties (i.e., mismatches) in these self-binding segments will remain within the final design, while relying on the surrounding complementary scaffold to maintain the global 3D fold. Once the mRNA-Design algorithm has completed the integration of the core gene and assigning the structuring domain, the 5’ and 3’ UTRs as well as a polyadenylate tail are appended onto the structure outside the structured region (Supplementary Fig. S1f-g).

Following the completion of the sequence design algorithm, the mRNA-Design program performs automated thermodynamic evaluation, exports a molecular dynamics ready 3D model of the structure, and exports an annotated sequence file. (Supplementary Fig. S1h-j). The thermodynamic evaluation compares every continuous duplex domain and calculates their nearest-neighbor free energies (ΔG) at 37°C using NUPACK^26^. By this calculation of the free-energy shifts of the redesigned structural domains relative to the original topology, the tool validates the thermodynamic viability of the newly designed mRNA-OG, with large deviations indicating high likelihood of misfolding, in which case a different input structure needs to be used for the design. Ultimately, the pipeline exports thermodynamically validated 3D models (for visualization in oxView^27,28^) and fully annotated sequence files for downstream laboratory use, providing a rapid, automated bridge between in silico nanostructure design and experimental synthesis (Supplementary Fig. S1i-j).

We designed the mRNA-OG construct to fold the open reading frame of the EGFP reporter gene via predefined hybridization with the structuring domain. All unstructured mRNA regulatory elements (i.e., 5’ and 3’ UTRs, CleanCap 5’ Cap 1 structure, and poly(A) tail) extend outwards from a single corner of the rectangular mRNA-OG to facilitate UTR recognition by eIF complexes. The first 40 nucleotides of the coding sequence also remained unstructured to allow for ribosome scanning and identification of the ribosomal binding site^29^ (Fig. 1d).

The mRNA-OG was self-assembled via thermal annealing from 85°C to 25°C at 1°C/s or 5°C/s in 1X PBS, pH 7.4 supplemented with 0.25 M NaCl. These conditions were selected based on fold state and monodispersity as assessed by atomic force microscopy (AFM) and agarose gel electrophoretic mobility shift assay (EMSA) across a range of buffer and salt conditions (Fig. 1e-g, Supplementary Figs. S4, S5). The resulting folded nanoparticles adopted the expected designed rectangular morphology with high monodispersity, as visualized by AFM (Fig. 1f-g). They also migrated as a discrete, compact band shifted relative to the unannealed transcript by EMSA, consistent with successful folding. We performed a slightly modified 2.5 hour fast annealing protocol^16^ and yielded comparable structural results. To assess the impact of structure on downstream applications, we tested both the annealed mRNA-OG (which we refer to as “Folded OG”) (Fig. 1e-f) and the unannealed construct that is not expected to have a compact morphology (referred to as “Unfolded OG”) (Supplementary Figs. S2b, S7 left). A control mRNA lacking the structuring domain is designated as “Regular mRNA”. Comparing Folded and Unfolded OG isolates the effect of tertiary structure on translation, while comparing Unfolded OG to Regular mRNA controls for the presence of the structuring domain sequence itself.

We quantified fold state via gel EMSA densitometry by individually measuring the intensity of bands corresponding to the monomer, dimer, and degraded species (n=3) and manual single-molecule structural classification from AFM (Extended Data Figs. 1-2). mRNA-OG annealed in 1X PBS supplemented with 0.25 M NaCl yielded nanostructures that were correctly folded 75.6 ± 2.7% of the time (n=606 molecules; 4 independent AFM images) with 59.8 ± 1.1% monodispersity (n=13 gel EMSA samples) (Extended Data Figs. 1-2). Dimers and trimers constituted 22.0 ± 0.6% and 15.7 ± 0.6% of band intensity, respectively, while degraded products accounted for only 2.5 ± 0.5%. Qualitatively, the dominant driver of the aggregation is not misfolding of products, but rather interactions between ssRNA UTRs and ssRNA loops protruding from the nanostructure’s 15 nm edge (Fig. 3b, top) as seen in Supplementary Fig. S6.

### Translation of mRNA-OG

The properties of mRNA-OG structure-dependent translation can be subject to artifactual results from nanostructure fold state heterogeneity, cellular stress-dependent autofluorescence, and transfection mechanism biases. Thus, we employed complementary approaches to monitoring the translation of the Folded and Unfolded versions of mRNA-OG, including EGFP reporter quantitation, rigorous nanoparticle characterization prior to transfection, and polysome profiling (Fig. 2a) of lipofected mRNA-OG in HEK293T cells 24 hours post-transfection.

**Fig. 2.**
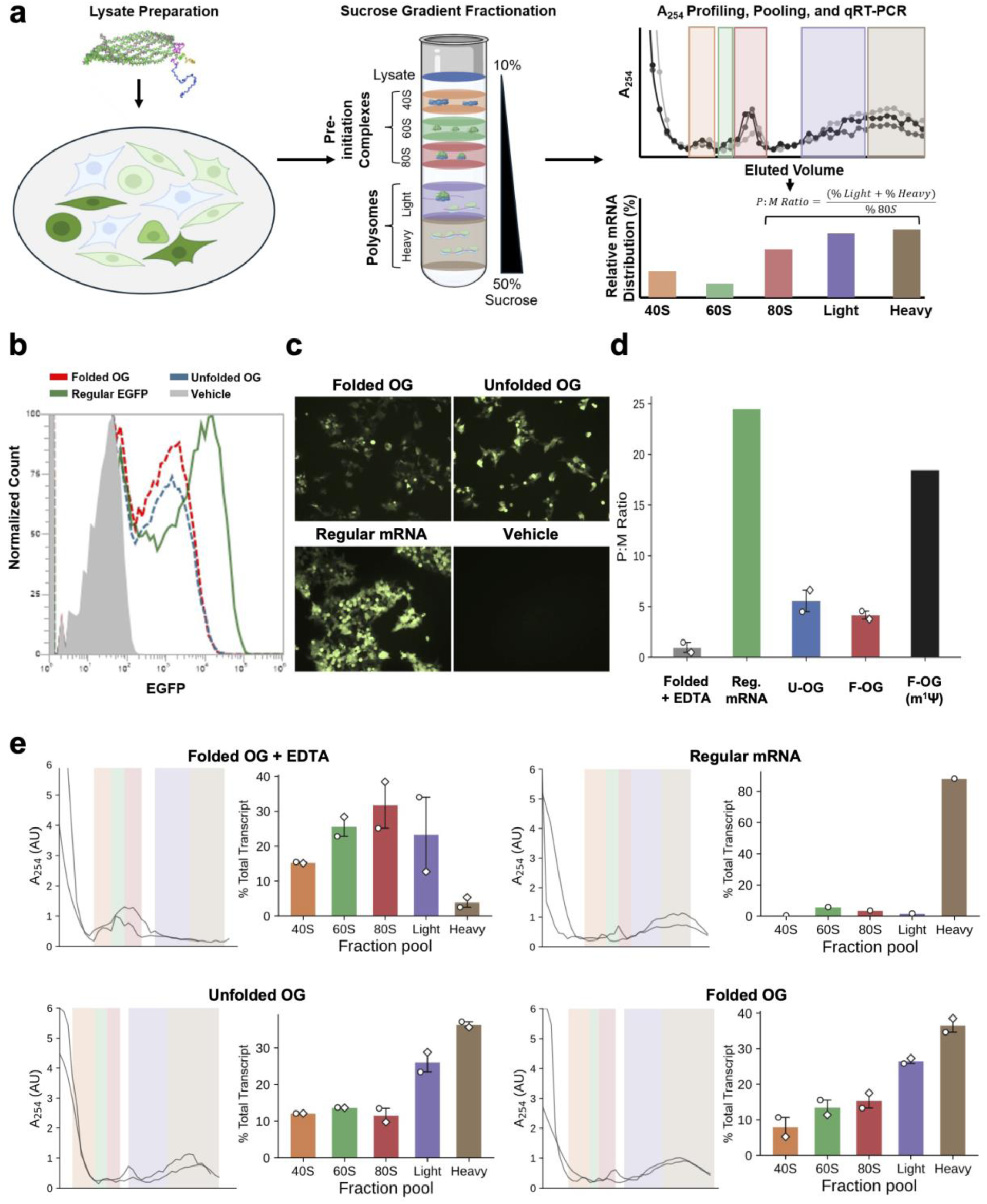
Characterization of EGFP-encoding mRNA origami nanoparticle translation in vitro. **a,** HEK293T cell lysate transfected under identical conditions was fractionated based on sedimentation coefficient through 10-50% sucrose gradients and split into 40 equal-volume fractions. A_254_ traces across all fractions were used to pool fractions corresponding to pre-initiation complexes (40S, 60S), monosome-associated transcripts (80S), or Light and Heavy polysome content (Light, Heavy Polysomes). EGFP-origami or control Regular mRNA copy number distribution was then assessed across the Trizol-LS-purified pools with origami-specific qRT-PCR primers. **b,** HEK293T cells were lipofected with 1 μg of Folded or Unfolded mRNA-OG nanoparticle or sequence-matched Regular EGFP mRNA and evaluated for reporter intensity after 24 hours via flow cytometry (n=1). **c,** FITC-channel fluorescence micrographs at 24-hour timepoint for Folded and Unfolded EGFP origami nanoparticles and positive control Regular EGFP mRNA. **d,** Aggregated polysome to monosome ratio for each sample (n=1 or n=2). F-OG (m^1^Ψ) is Folded mRNA-OG synthesized with 100% substitution of uridine for N1-methylpseudouridine (m^1^Ψ). Replicate 1 shown as circles, replicate 2 shown as diamonds. **e,** A_254_ profiles (left) and mRNA distribution (right) across pools (n=1 or n=2). EDTA was added to disrupt mRNA-OG structure. Replicate 1 shown as circles, replicate 2 shown as diamonds.

Flow cytometry quantification of the EGFP expression in transfected cells was used initially to compare mRNA-OG and Regular mRNA reporter brightness and delivery efficiency. In HEK293T cells subject to equimolar lipofection, we observed reporter expression in 82-91% of OG-transfected cells and 97% of cells transfected with Regular mRNA. The comparable delivery efficiency was not paralleled by per-cell reporter expression, however. Both Folded and Unfolded mRNA-OG consistently yielded approximately 20% of reporter mean fluorescence intensity (MFI) relative to Regular mRNA (Fig. 2b). This transcript potency gap was not rescued by further RP-FPLC purification of the transcripts, ruling out reduced mRNA-OG IVT product quality compared with the Regular mRNA control. Notably, we were not able to utilize the gold standard J2 anti-dsRNA antibody dot blot to quantitate spurious IVT products in our nanoparticle preparations due to a false positive signal from the folded nanoparticle itself (Supplementary Fig. S14). To rule out nanoparticle unfolding as a result of lipofectamine complexation, we electroporated Folded OG, Unfolded OG, and Regular mRNA reporter into primary dendritic cells (DCs) isolated from human PBMCs, achieving 70–85% delivery efficiency in matured DCs and more variable delivery (53–95%) in immature DCs, with both Folded and Unfolded transcripts showing a comparable loss in transcript potency to ∼5–8% of Regular mRNA across both maturation states (Fig. 5a-c). Moreover, to rule out artifactual unfolded nanoparticle biasing, we cross-titrated annealed (Folded) versus unannealed (Unfolded) EGFP-OG in HEK293T cells and observed no dose-dependent Unfolded OG fluorescence increase at 24 hours via flow cytometry (Extended Data Fig. 6). We also titrated increasing Folded mRNA-OG into a constant Unfolded payload and observed no change in MFI or dose-dependent toxicity from the Folded nanostructure.

Polysome profiling was used to validate previous results and demonstrated equivalent recruitment of translation machinery between Folded and Unfolded mRNA-OG at 24 hours post-lipofection. In polysome profiling, actively translated mRNAs co-sediment with heavy ribosomal complexes (polysomes), while untranslated or poorly translated transcripts remain associated with the monosome (80S) fraction or lighter subunits. The polysome-to-monosome (P:M) ratio—the fraction of a transcript found in polysomal fractions relative to the monosome—therefore provides a direct measure of translational engagement, with higher values indicating more efficient ribosome loading and active translation^30–32^. Folded and Unfolded OG demonstrate similar performance, with 63.2% and 62.6% of OG transcripts fractionating with the polysome fractions (Fig. 2e), and Folded and Unfolded P:M ratios of 2.40 ± 0.21 and 3.21 ± 0.46 (n=2) (Fig. 2d). Regular mRNA recruited significantly more polysomes as expected from reporter quantification, with a P:M ratio of 24.0. Interestingly, substitution of uridine with m^1^Ψ raised the Folded OG P:M ratio to 17.3, representing a 7.2× increase in ribosome recruitment; however, as we demonstrate later this increase in P:M ratio does not correspond to a commensurate increase in EGFP reporter expression (Fig. 3e). We suspect that the increased P:M ratio without increased expression is due to ribosomal stalling on stretches of modified nucleotides, as has been previously described^33–35^. As expected, EDTA-chelation of Mg^2+^ ions from a Folded mRNA-OG sample prior to fractionation on a sucrose gradient (serving as a negative control) yielded no heavy polysomal transcripts, demonstrating that the polysomal fractions observed in the non-EDTA-treated samples was not a result of transcript structure affecting sedimentation coefficient.

**Fig. 3.**
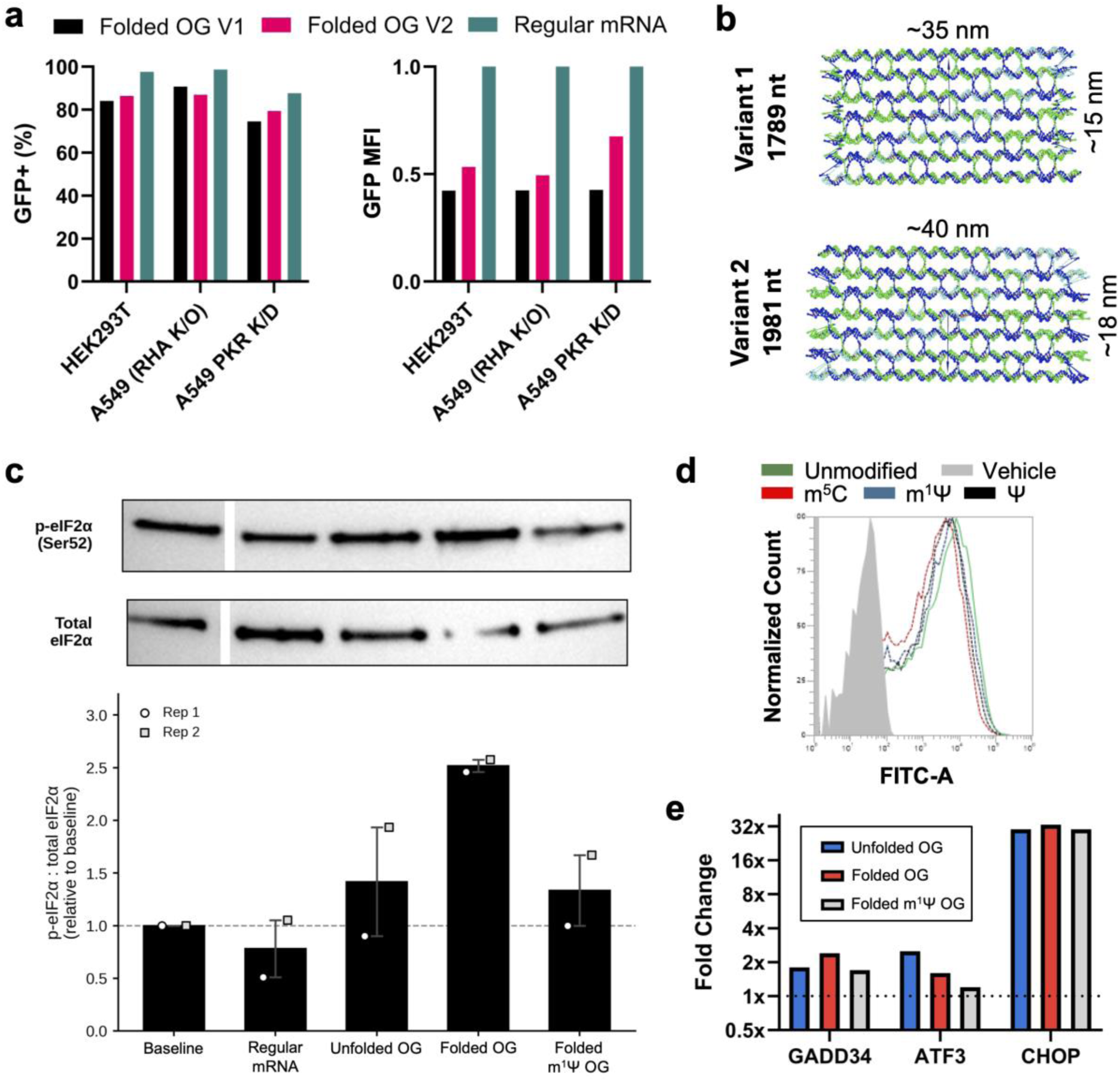
mRNA origami engages subthreshold ISR activation without PKR-dependent translational attenuation. **a,** Transfection efficiency (left) and EGFP mean fluorescence intensity (right) of Folded mRNA-OG (Package Variants 1 and 2) and Regular EGFP mRNA in HEK293T wild-type, A549 RHA knockout, and A549 PKR knockdown cells (n=1). PKR knockdown reduces EGFP expression equivalently across all constructs, indicating the translation deficit of Folded OG is structural rather than ISR-mediated. RHA knockout has no effect, suggesting nanostructure denaturation prior to translation is accomplished by the ribosome itself rather than cytoplasmic helicases. **b,** Structural models of the two mRNA-OG package variants, differing in dsRNA helical edge length (Variant 1: ∼35 nm, 1789 nt; Variant 2: ∼40 nm, 1981 nt). **c,** Western blot quantification of phospho-eIF2α normalized to total eIF2α in HEK293T cells 24 hours post-lipofection (n=2). Folded OG elicits elevated eIF2α phosphorylation relative to Unfolded OG and Regular EGFP mRNA. **d,** Flow cytometry histograms of EGFP expression from Folded mRNA-OG synthesized with unmodified nucleotides or 100% substitution of 5-methylcytidine (m^5^C), N1-methylpseudouridine (m^1^Ψ), or pseudouridine (Ψ) (n=1). Modified nucleotide incorporation does not alter reporter brightness. **e,** qRT-PCR quantification of ISR transcripts GADD34, ATF3, and CHOP in HEK293T cells 48 hours post-electroporation with Unfolded, Folded, or Folded m^1^Ψ-modified mRNA-OG, expressed as fold-change over untransfected control (ΔΔCt, input mass-normalized). GADD34 shows minimal induction across all conditions (1.5–2.4×), consistent with low-grade, compensated ISR signaling. CHOP is uniformly elevated ∼30-fold regardless of mRNA structure or nucleotide modification, consistent with delivery vehicle-induced ER stress rather than dsRNA-specific sensing.

The reduction in EGFP reporter fluorescence and P:M ratio relative to a Regular EGFP mRNA control appears to implicate both reduced ribosome processivity as well as decreased ribosome reinitiation efficiency due to the multi-kilobase non-coding structural sequence between the stop codon and poly(A) tail in the nanoparticle. Comparable P:M ratios between Folded and Unfolded versions of the mRNA-OG, however, rule out transcript secondary structure as the primary factor in reduced reporter expression. Furthermore, the downstream noncoding structural context between the stop codon and poly(A) tail has been demonstrated to reduce translation termination and reinitiation by reducing interactions between the ribosomal complex and poly(A)-binding proteins (PABP)^36–40^. Optimal mammalian stop-PABP (i.e. 3’ UTR) spacing appears to be about 300 nucleotides^41^ of highly structured RNA; however, the mRNA-OG EGFP construct contains 1,133 nucleotides that would be expected to negatively affect translation of Unfolded and Folded transcripts similarly.

### Structure-Dependent Pattern Recognition Receptor Engagement Without ISR-Mediated Translational Arrest

The elongation stalling mechanism underlying the mRNA-OG translation deficit does not require integrated stress response (ISR) pathway activation. To determine whether mRNA-OG nonetheless engages canonical dsRNA-sensing pathways in parallel with ribosome transit, we assessed activation of PKR, MDA5, and RIG-I using orthogonal readouts: 1) phosphorylation of their shared downstream effector eIF2α, 2) expression of ATF4-regulated stress transcripts, and 3) type I IFN secretion. Robust ISR activation would be expected to manifest as sustained eIF2α phosphorylation, induction of CHOP and GADD34, and detectable IFN-β. Such a profile would confirm that the nanostructure’s dsRNA character is immunologically legible even in the absence of a functional translation deficit.

We first assessed IFN-β secretion 24 hours post-lipofection in both HEK293T and THP-1^42^ cells transfected with Folded or Unfolded mRNA-OG, Regular mRNA, or poly(I:C), a dsRNA mimic often used to simulate viral infection that triggers IFN-β secretion (Extended Data Fig. 3). HEK293T cells produced no detectable IFN-β under any condition, consistent with the well-documented STING deficiency^43–45^ of this line and its broadly attenuated innate sensing capacity. In THP-1 monocytes, by contrast, Folded mRNA-OG produced a statistically significant elevation in IFN-β relative to baseline (p=0.040), while Unfolded mRNA-OG showed a directional but non-significant trend, and Regular mRNA remained at baseline. Poly(I:C) produced the highest output, confirming assay sensitivity and establishing that neither mRNA-OG fold variant approaches the response of a canonical dsRNA agonist. The conformation-dependent hierarchy in THP-1 cells (Folded > Unfolded > Regular mRNA) is consistent with the compact PX scaffold in Folded OG engaging cytosolic PKR through proximity-dependent dsRBD bridging following endosomal escape of lipoplex cargo. TLR3 is unlikely to be the dominant sensor despite the endosomal delivery route, as undifferentiated THP-1 cells express low TLR3 and respond weakly to extracellular poly(I:C)^46^, though responsiveness varies with subclone and differentiation^47–50^; the IFN-β signal therefore reflects cytosolic pattern recognition receptor (PRR) engagement downstream of lipoplex endosomal escape.

After establishing that Folded mRNA-OG engages cytosolic PRR signaling in a conformation-dependent manner, we next asked whether PKR activation specifically accounts for the translation deficit observed across mRNA-OG constructs relative to Regular mRNA. We tested this by assessing differential mRNA-OG reporter expression in cell lines with knocked down PKR and RNA helicase A (RHA), which feed directly into the phospho-eIF2α and type I IFN-β pathways respectively (Fig. 3a). To assess the effect of nanostructure size on PKR and RHA activity, we also designed a new mRNA-OG variant with the same mRNA coding sequence by utilizing an alternative, previously published ssRNA-OG shape^21^ with a longer dsRNA helical edge length but similar topology and strand routing (40 nm vs 35 nm, see Fig. 3b) as input to our design algorithm. The RHA K/O condition showed no effect on nanoparticle transfection or expression, suggesting that nanostructure denaturation prior to translation is not achieved by cytoplasmic helicases like RHA, but rather accomplished by the translation complex itself due to the nanostructure’s knotless topology. PKR knockdown reduced EGFP expression equivalently across Folded OG, Unfolded OG, and Regular mRNA conditions, arguing against a selective role for PKR in attenuating nanostructure translation. If PKR activation were responsible for the observed translation deficit in Folded OG, its knockdown would be expected to preferentially rescue reporter expression relative to controls; the absence of selective rescue indicates that PKR-dependent ISR is unlikely to be the primary driver of the translation deficit, suggesting that structural features of the construct play a dominant role.

To assess PKR’s role in mRNA-OG sensing and integrate the orthogonal ISR sensors MDA5, TLR3, and RIG-I activation states, we next directly measured phosphorylation of eIF2α post-lipofection into HEK293T cells. Phospho-eIF2α is a widely used marker of ISR activation and reflects a key step in translational reprogramming toward cap-independent translation of stress transcripts like ATF4. Thus, we quantified phospho-eIF2α relative to total eIF2α from HEK293T lysate 24 hours post-transfection with Folded or Unfolded mRNA-OG or Regular mRNA (Fig. 3c, Supplementary Fig. S15). Only Folded mRNA-OG consistently elevated p-eIF2α expression (normalized to total eIF2α) with respect to baseline (2.52 ± 0.06x), Regular mRNA and Unfolded mRNA-OG were similarly phosphorylated compared with baseline (0.78 ± 0.27× and 1.42 ± 0.52×, respectively). As expected, incorporation of N1-methylpseudouridine into the folded mRNA-OG at 100% substitution rate ablated the nanostructure’s stimulatory potential, resulting in phosphorylation similar to Unfolded mRNA-OG and baseline (1.33 ± 0.33×).

Interestingly, while the increase in ISR signaling is obvious in the Folded sample, it does not appear to be sufficient to fully suppress reporter translation under these conditions because the Unfolded and Folded mRNA-OG still yield the same quantity of reporter product as assessed via flow cytometry. Supporting this apparently insufficient phosphorylation, we also saw no change in mRNA-OG EGFP brightness when synthesized with 100% substitution of uridine or cytidine with pseudouridine, N1-methylpseudouridine, or 5-methylcytidine (Fig. 3d). Differences in reporter expression between unmodified and 100% modified mRNA-OG might be expected if ISR signaling were the dominant determinant of translation efficiency due to the well-documented ability of non-canonical nucleotides to ablate dsRNA detection. As an interesting aside, Ψ and m^1^Ψ incorporation had no effect on thermal annealing of our nanostructures, but m^5^C led to a high misfold rate as assessed via AFM (Supplementary Fig. S6).

As a final orthogonal approach to comprehensive stress state profiling, we utilized qRT-PCR to quantitate relative expression of several transcripts downstream of p-EIF2α/ATF4 post-lipofection of Folded, Folded m^1^Ψ-modified, or Unfolded mRNA-OG, compared with untransfected HEK293T cell lysate. ATF4’s downstream targets CHOP (apoptotic stress indicator) and GADD34 (eIF2α dephosphorylase and negative regulator of ISR) were chosen to differentiate between low-grade constitutive signaling from endogenous transcripts versus apoptotic stressing from exogenously delivered mRNA (Fig. 3e). For the purpose of directly assessing p-EIF2α-mediated translation, GADD34 is the best stress indicator because it is not a convergence node for stress-mediated mechanisms. We observed 2.4-fold, 1.7-fold, and 1.8-fold induction of GADD34 by Folded, Folded Ψ-modified, and Unfolded mRNA-OG over untransfected control, respectively (Fig. 3e). The modest expression of GADD34 in all samples, and the relatively small jump from 1.7 to 2.4-fold when omitting m^1^Ψ in the folded nanostructure, support the idea that the ISR signaling and stress induction of our nanoparticles is low-grade, non-cytotoxic, and compensated for by GADD’s dephosphorylation activity. Of note, a 35-fold induction of CHOP across the board was seen relative to untransfected control, including in the m^1^Ψ-modified sample. In contrast to GADD34, however, CHOP signaling can be activated by several converging inputs (NF-κB, PERK), including those consistent with ionizable lipid-induced ER stress mediated by the delivery mechanism itself, consistent with other reports^51^. This CHOP induction is internally consistent with the phosphorylation of eIF2α across the board observed in our western blotting analysis, as well as with the increased phosphorylation observed in the Folded mRNA-OG sample. ATF3, while not a mechanistic player in mediated ISR cytotoxicity, is an immediate-early reporter of ATF4 cap-independent translation, and its similar kinetics to GADD34 further support a modest stress-induction response.

The conformation-dependent PRR engagement and subthreshold ISR activation observed in HEK293T and THP-1 cells raises a translational question: whether the structural properties of mRNA-OG can be leveraged to tune innate sensing in primary antigen-presenting cells, where controlled PRR activation has well-established immunostimulatory consequences. Low-level type I IFN release downstream of cytosolic RNA sensing has been shown to drive paracrine HLA class I upregulation and co-stimulatory marker expression on surrounding cells^52,53^, an effect that is potentially advantageous in antigen-loading contexts but must remain below the threshold of cytotoxic ISR, as defined by sustained eIF2α phosphorylation, translational arrest, and CHOP-driven apoptosis. The preceding data demonstrate that mRNA-OG occupies this subthreshold regime in transformed cell lines. Whether this property is preserved in primary human monocyte-derived dendritic cells, and whether maturation state modulates the conformation-dependent differences in PRR engagement we have described, will be further addressed below.

### Enhanced Encapsulation, Delivery, and Cold Storage Longevity of mRNA-OG

The observation that Folded versus Unfolded mRNA-OG transfected more cells without an increase in reporter potency led us to next quantify mRNA-OG encapsulation efficiency (EE). Folded and Unfolded mRNA-OG and Regular mRNA transcripts were complexed with Lipofectamine MessengerMAX Transfection Reagent (Thermo Fisher Scientific) under standard conditions. Subsequently, free RNA was quantitated by SYBR Gold intercalation before and after Triton X-100-mediated lipoplex dissociation (Fig. 4a-b). We found that Folded mRNA-OG performed exceptionally well, with 91.1% of input RNA being encapsulated, compared to 13.9% for Unfolded OG. Comparatively, Regular mRNA resulted in only 38.5% EE. We suspect the compact size and structural rigidity of the Folded mRNA-OG promotes efficient encapsulation through its regular patterning of surface charges, allowing polyanionic reagents to interact cooperatively, which may in turn reduce the energetic barrier of dissociating the hydration shell. The gap in EE performance between Unfolded origami and Regular EGFP mRNA is likely due to the origami’s native, incorrectly base-paired structure resembling a knotted string ball without an ordered, evenly charged surface (Supplementary Fig. S7).

**Fig. 4.**
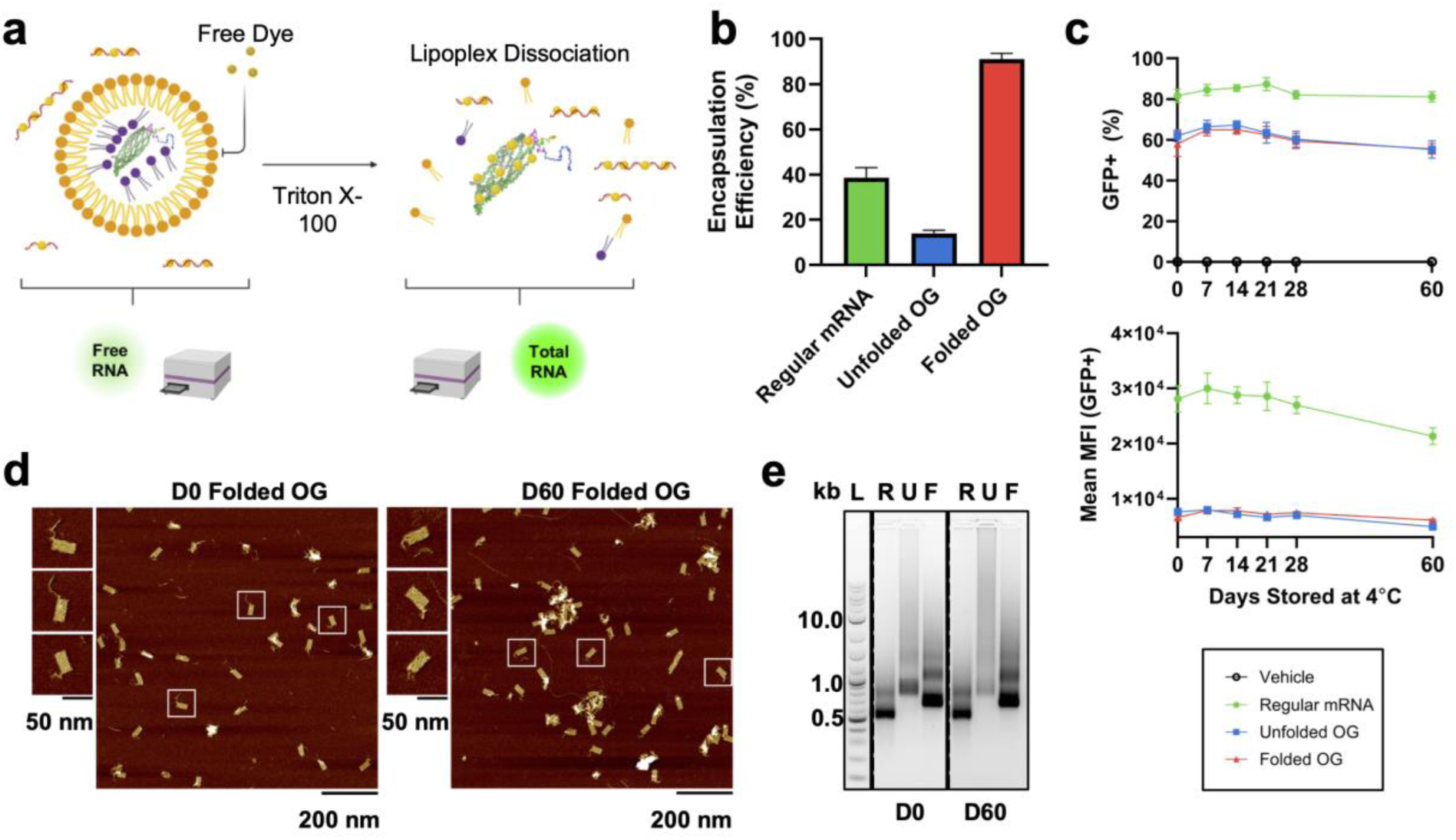
Lipofectamine encapsulation efficiency and mRNA-OG stability over 60 days at 4°C. **a,** Schematic of the SYBR Gold intercalation-based encapsulation efficiency (EE) assay. Free RNA fluorescence is measured prior to lipoplex dissociation; total RNA fluorescence is measured following Triton X-100-mediated lysis. Background-corrected EE is calculated as %EE = 100 × (1 − F_pre,c_ / F_post,c_). **b,** Encapsulation efficiency of Regular mRNA, Unfolded OG, and Folded OG complexed with MessengerMAX lipofectamine. Bars represent mean ± SD; n=4 independent replicates per condition. **c,** GFP+ transfection efficiency (%, top) and GFP+ mean fluorescence intensity (MFI, bottom) of HEK293T cells lipofected with mRNA stored at 4°C in annealing buffer (1X PBS + 0.25 M NaCl) and assessed at 24 hours post-transfection across 60 days of storage. Lines represent mean ± SD; n=3 technical replicates per timepoint. **d,** Representative AFM images of Folded OG at day 0 (D0) and day 60 (D60) with zoomed insets (scale bars: 50 nm inset, 200 nm wide-field). **e,** Agarose gel EMSA of RNA samples at D0 and D60. Lanes: L = ladder, R = Regular mRNA, U = Unfolded OG, F = Folded OG. Minimal degradation is observed across all conditions over 60 days. Collapse of the diffuse Unfolded OG band into a sharper lower-mobility band at D60 is consistent with spontaneous thermal self-annealing observed by AFM (Supplementary Fig. S7).

Interestingly, the divergence in EE of Folded versus Unfolded mRNA-OG with identical sequences suggests transfection reagent biasing toward delivering compact, folded transcripts while omitting partially folded or misfolded ones. Although our gel EMSA and AFM results already demonstrate folded nanoparticle monodispersity and fold efficiencies between 60-70%, in this case this lipofection bias may work as a structure-based purification mechanism, with LNP complexation screening transcripts with non-ideal fold states from interfering with folded transcript translation experiments.

To investigate whether the double-stranded nature of mRNA-OG impacts RNA storage stability, we examined Folded and Unfolded mRNA-OG alongside Regular mRNA for degradation, structure, and expression over 60 days of storage at 4°C in 1X PBS supplemented with 0.25M NaCl. At each timepoint, RNA was characterized by agarose gel EMSA, AFM, and lipofection into HEK293T cells followed by flow cytometry (Fig. 4c). All constructs demonstrated functional expression across the full 60-day storage period in Mg-free buffer, with no evidence of progressive degradation-driven decline in per-cell expression or delivery efficiency through Day 28. A modest decline in MFI was observed at Day 60 for all constructs, most pronounced for Unfolded mRNA-OG, consistent with gradual RNA degradation through decapping or in-line hydrolysis over extended storage rather than acute structural instability. Transfection efficiency similarly showed no progressive decline, with all constructs maintaining delivery into HEK293T cells throughout the storage period. RNA integrity was preserved across all conditions as assessed by agarose gel EMSA, with no evidence of significant degradation over 60 days (Fig. 4e, Extended Data Fig. 4). AFM imaging revealed slight increases in particle aggregation at later timepoints but intact monomer structure throughout (Fig. 4d, Extended Data Fig. 5). Of note, after 21 days of storage at 4°C, some Unfolded mRNA-OG sequences spontaneously adopted a folded conformation detected by AFM (Supplementary Fig. S7), consistent with the thermodynamic favorability of the designed PX architecture even in the absence of thermal annealing^54^.

### Extending mRNA-OG to a Translational Platform in Primary Human Dendritic Cells

The preceding data establish that mRNA-OG is translationally competent in adherent and suspension cell lines, and that its conformation-dependent PRR sensing produces differential ISR activation in those systems. To assess platform viability in a clinically relevant primary cell type, and to evaluate whether the conformation-encoded innate sensing observed in THP-1 cells is preserved in professional antigen-presenting cells, we apply the mRNA-OG platform to primary human monocyte-derived DCs (moDCs). We generated both immature (iDC) and TNFα/PGE_2_-matured (mDC) moDCs from healthy donor CD14+ monocytes (Donors D2–D5; see Methods, Supplementary Table S3) and electroporated each population with molar-equivalent doses of Folded mRNA-OG, Unfolded mRNA-OG, or Regular mRNA (Fig. 5a). Conditioned supernatants and phenotypic readouts were collected 48 hours post-electroporation without secondary stimulation or CD40 ligation, such that all outputs reflect the cells’ endogenous response to the RNA payload and electroporation (EP) procedure alone.

**Fig. 5.**
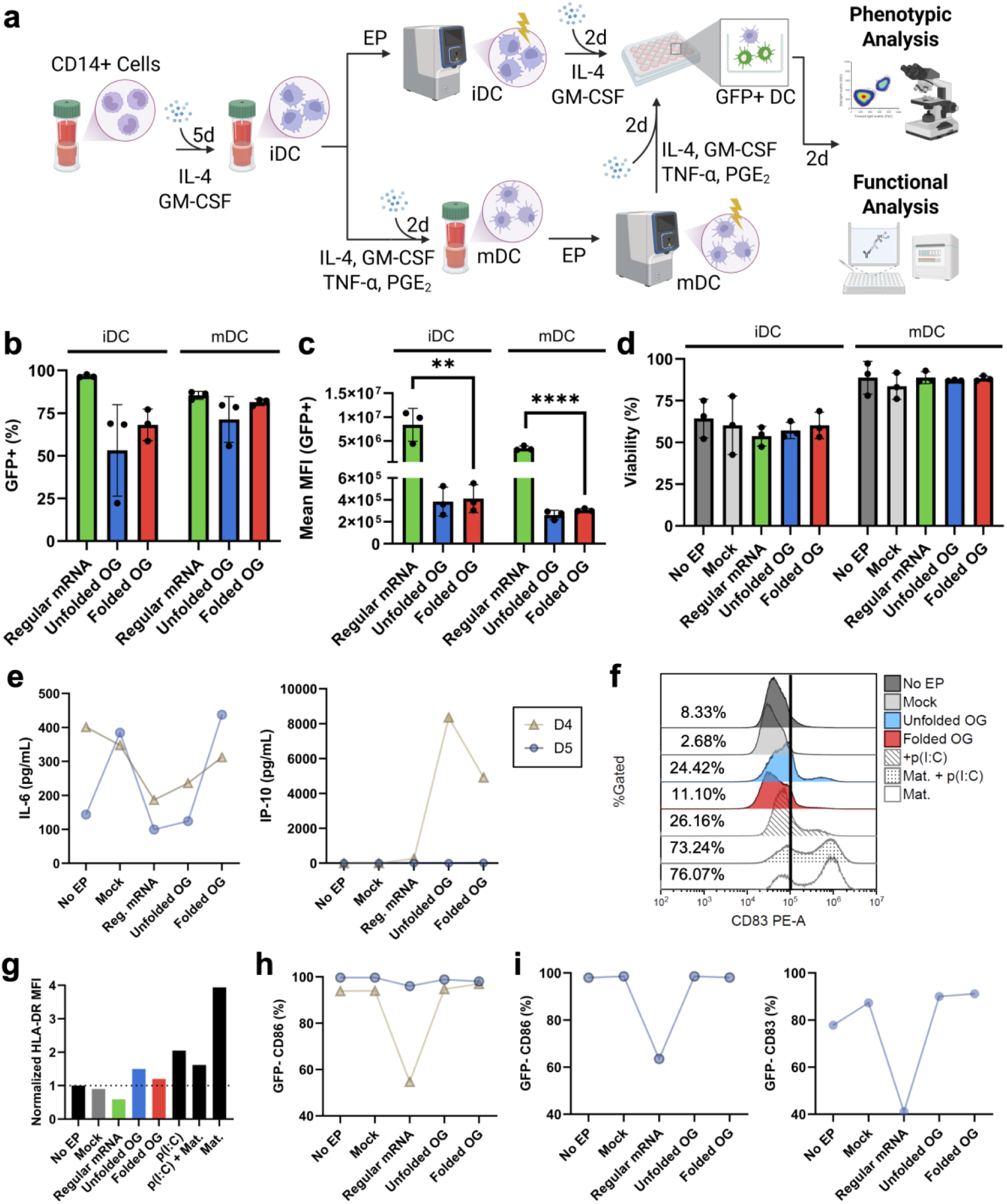
mRNA-OG is translationally competent in primary human monocyte-derived dendritic cells. **a,** Schematic of the experimental workflow. All conditions received molar-equivalent RNA doses (235 pM). Conditioned supernatants and phenotypic readouts were collected at 48 hours post-electroporation. **b,** GFP+ transfection efficiency (%) **c,** GFP+ mean fluorescence intensity (MFI), and **d,** post-electroporation viability (%) assessed by live/dead staining, at 48 hours post-electroporation. Bars represent mean ± SD; individual donor points shown. n=3 biological replicates for iDC; n=3 biological replicates for mDC. No EP = cells not subjected to electroporation. GFP+ cell populations were gated on live (7-AAD-) cells. GFP MFI statistical comparisons by one-way ANOVA with Tukey’s post-hoc test; ** p<0.01, **** p<0.0001 vs Regular mRNA within each maturation state. **e,** Selected results from multiplexed secretome of conditioned supernatants at 48 hours post-electroporation of immature DC. IL-6 (left) and IP-10/CXCL10 (right) are shown as individual donor values; D4 shown as gold triangles, D5 as blue circles. Donor 3 excluded from secretome analysis due to non-standard 24-hour collection timepoint. Data represent technical duplicates within each donor. Full analyte panel (MIP-1β, MCP-1, IL-8, IL-10) provided in Supplementary Tables S4, S5. mDC secretome data provided in Supplementary Fig. S10. **f,** CD83 surface expression in Donor 4 iDCs 48 hours post-electroporation. Left: representative stacked histogram overlay of CD83-PE fluorescence on all live cells. Traces shown for No EP (dark grey), Mock (light grey), Unfolded OG (blue), Folded OG (red), +p(I:C) (hatched), Mat. + p(I:C) (dotted), and Mat. cocktail (outline). Regular mRNA excluded from histogram display due to GFP spectral spillover into the PE channel at high transfection efficiency in this donor (GFP GMean = 9,368,469); this condition is represented quantitatively in Panel G. Vertical line indicates CD83+ gate threshold. %Gated values for each condition are annotated. **g,** Normalized HLA-DR MFI across iDC conditions in Donor 4. HLA-DR (PB450) GMean reported for GFP+ cells in electroporated RNA conditions (Regular mRNA, Unfolded OG, Folded OG); bulk live-cell GMean reported for non-electroporated reference conditions (No EP, Mock) and stimulation benchmarks (p(I:C), p(I:C) + Mat., Mat. cocktail) where no GFP+ population is present. All values normalized to No EP GMean (dashed reference line at 1.0). **h,** GFP-negative bystander CD86 expression (%) in iDC at 48 hours post-electroporation. Individual donor values shown as connected lines. **i,** GFP-negative bystander marker expression in D4 iDCs receiving post-EP maturation cocktail (TNFα/PGE_2_ applied equivalently to all conditions following electroporation). Left: GFP-negative CD86 (%). Right: GFP-negative CD83 (%). Points are connected by donor to visually track how each donor’s response shifts across electroporation conditions.

Electroporation represents a clinically precedented route for mRNA delivery into primary human moDCs, with multiple groups demonstrating efficient transgene delivery and antigen-specific T cell stimulation across maturation states^55–57^. Prior work has established that mDCs electroporated with antigen-encoding mRNA exhibit superior post-EP viability, transgene expression, and antigen-specific T cell stimulatory capacity relative to iDCs loaded under the same conditions^58–60^, positioning mDC electroporation as the preferred arm for future mRNA-OG antigen loading studies.

In the mDC arm, all three constructs produced comparable GFP+ cell percentages across donors, confirming equivalent delivery efficiency in the mature state (Fig. 5b). In the iDC arm, more significant inter-donor variability was seen in the efficiency of transfection. The observed mildly reduced iDC mRNA delivery efficiency relative to that of mDC is attributed to maturation-state-dependent differences in sensitivity to electroporation-induced stress rather than cytotoxicity of the mRNA-OG itself, as viability was maintained across all iDC conditions (Fig. 5d). Whether innate sensing additionally contributes to cell loss in mRNA-OG-transfected iDCs cannot be determined without formal apoptosis profiling stratified by transfection status. Despite these delivery differences, GFP MFI was significantly reduced for both mRNA-OG constructs relative to Regular mRNA in both iDC and mDC when assessed within GFP+ cells (Fig. 5c; p<0.01 iDC, p<0.0001 mDC, Supplementary Fig. S9). Folded and Unfolded mRNA-OG were not statistically distinguishable from each other in either maturation state, consistent with the polysome profiling result. The preservation of this translation deficit across delivery routes and primary cell types confirms that elongation stalling on the mRNA-OG scaffold is a structural property independent of delivery mechanism or cell type.

To characterize the innate immune secretory response to structured mRNA delivery beyond single-analyte readouts, conditioned supernatants were profiled across 22 analytes spanning pro-inflammatory cytokines, chemokines, and effector proteins. Multiplexed cytokine analysis of conditioned supernatants (Bruker IsoLight CodePlex Adaptive Human chip; n=2 iDC: D4, D5; n=4 mDC: D2, D3, D4, D5, with D3 collected at 24 h) revealed analyte-selective rather than broadly inflammatory secretion across both maturation states. IL-10 was below the assay limit of detection in all conditions and both cell types, providing no evidence of tolerogenic polarization. MIP-1β (a chemokine produced downstream of NF-κB activation that recruits monocytes and T cells), MCP-1 (a monocyte chemoattractant), and IL-8 (a general inflammatory stress marker) were present at comparable levels across RNA conditions and across non-electroporated and mock controls in both maturation states, identifying them as constitutive DC products rather than RNA-or EP-induced and arguing against nonspecific NF-κB activation by structured RNA delivery. Full analyte reporting is provided in Supplementary Tables S4 and S5. IL-6, an NF-κB-dependent, broadly pro-inflammatory cytokine produced by activated DCs, showed a cross-donor consistent ordering in iDC: Regular mRNA produced the lowest IL-6 among RNA conditions in both donors, while Folded mRNA-OG produced the highest, with Unfolded mRNA-OG intermediate (Fig. 5e, left). However, all RNA conditions remained within ∼0.5 log10 of the non-electroporated and mock baselines, so this ordering is small in magnitude and directional only. In mDC, Folded mRNA-OG produced the lowest IL-6 of the three RNA conditions in every donor with detectable IL-6 (3 of 4; the remaining donor, D5, had no detectable IL-6 in any condition). The iDC secretomes showed an inverse pattern, in which Folded was highest. The ranking of Regular versus Unfolded mRNA-OG was not consistent across donors (Regular > Unfolded in D2 and D3, Unfolded > Regular in D4), so only the Folded-lowest pattern is reproducible (Supplementary Fig. S11, Supplementary Table S4). These data cannot be tested statistically given the donor numbers and the technical-duplicate chip architecture, and the construct differences sit near the secretory baseline; we therefore present the Folded-lowest mDC pattern and the inverse iDC ordering as directional observations rather than as evidence for a specific signaling mechanism.

IP-10 (CXCL10), whose production requires active type I IFN signaling through IFNAR, making it a functional reporter of the complete dsRNA → IFN-β axis rather than PRR activation alone^61,62^, showed a striking RNA-dependent signal in Donor D4 iDC, where both mRNA-OG conformations elevated IP-10 far above Regular mRNA and mock-electroporated controls (Unfolded ∼8,400 pg/mL; Folded ∼4,900 pg/mL). IP-10 was the only analyte in the panel that was at the assay floor in mock and non-electroporated controls, yet strongly induced by RNA, identifying it as the single unambiguously RNA-dependent secretome readout. This induction was confined to one donor: in Donor D5 iDC, IP-10 was near zero across all RNA conditions (Fig. 5e, right). The mRNA-OG-above-Regular and Unfolded-above-Folded relationships therefore rest on a single biological replicate. IP-10 was at or below the assay floor across all OG and mock conditions in mDC across four donors at two timepoints (D2, D3, D4, D5; 24h and 48h), suggesting that the mDC IP-10 null reflects suppression of the IRF3/ISRE axis in the mature state rather than a temporal artifact due to the collection window (Supplementary Fig. S10, Supplementary Table S5). The reproducible distinction in these data is between mRNA-OG and Regular mRNA rather than between fold states. Where a fold-state comparison could be made, its direction was inconsistent across assays — Unfolded exceeded Folded for IP-10 in D4 iDC, whereas Folded exceeded Unfolded for IFN-β in THP-1 cells and for eIF2α phosphorylation in HEK293T cells — and each rests on one or two replicates. In the single donor where the type I IFN/ISRE axis was active, both mRNA-OG conformations engaged it more strongly than Regular mRNA.

The within-donor IP-10 suppression in D4 mDC relative to D4 iDC is consistent with PGE_2_-mediated attenuation of the IRF3-dependent arm of cytosolic RNA sensing in the mDC state, as PGE_2_ has been shown to inhibit type I IFN production in human myeloid DCs responding to poly(I:C)^63^ and to suppress IRF3-driven antiviral responses via EP2/EP4 receptor signaling^64^. The iDC IL-6 ordering (Folded > Unfolded > Regular) was consistent in direction across both donors, but the differences are small and sit near the secretory baseline; we therefore describe it as the most reproducible construct ordering in the secretome. The apparent IL-6 inversion between maturation states is likewise a directional observation; the donor numbers, the proximity to baseline, and the opposing fold-state directions seen across our other assays preclude assigning it to a defined mechanism.

Though the 48-hour collection timepoint for Donors 2, 4, and 5 was chosen to correspond to peak transgene expression, Donor 3 datapoints collected at 24 hours showed similarly null IP-10 signals across all OG and mock mDC conditions. This result, coupled with the IP-10 null confirmation across four donors, provides no evidence that earlier collection would reveal higher OG-specific cytokine outputs in the mature state. The null IP-10 mDC secretome is therefore more likely a consequence of PGE_2_-mediated innate sensing suppression than a temporal artifact. An important residual caveat is that the TNFα/PGE_2_ maturation cocktail used here suppresses type I IFN secretory capacity in mDC, limiting detection of fold-state-dependent innate sensing differences that would be apparent in IFN-I-competent mDC formats. Alternative maturation approaches preserving IFN-I competency have demonstrated clinical proof-of-concept for mRNA-electroporated DC vaccines encoding patient-specific neoantigens where EP-loaded mDCs induced durable T cell responses detectable up to 19 months post-vaccination^65^. IFN-competent maturation of mRNA-OG-loaded DCs represents an ideal next step for evaluating the full innate sensing profile of mRNA-OG in a clinically translatable context.

Beyond secretory output, mRNA-OG electroporation induced maturation-related phenotypic changes in iDCs from Donor 4. Live, GFP+ iDC CD83+ surface expression revealed that Unfolded (24.4%) and Folded OG (11.1%) both exceeded the No EP (8.3%) baseline and mock (2.7%) transfected samples, while not quite meeting the poly(I:C) benchmark (26.2%) (Fig. 5f, Supplementary Figs. S12, S13). Quantitative analysis of HLA-DR, a marker that is upregulated during DC maturation, mirrored this CD83 trend. Though HLA-DR expression was >98% in all conditions, Unfolded and Folded OG GFP+ cells showed HLA-DR above the No EP baseline (at 1.50× and 1.20×, respectively), while Regular mRNA-transfected GFP+ cells fell below the baseline at 0.59× (Fig. 5f). Though both mRNA-OG conformations induced partial maturation above the No EP and Regular mRNA baselines across CD83 and HLA-DR readouts in this donor, and while directional differences between conformations were observed, the two constructs are considered roughly equivalent at n=1 donor, where biological replication is insufficient to distinguish a true conformation-dependent difference from donor-specific variation. However, these partial maturation signatures may indicate that a subthreshold activation of innate sensing is occurring in these cells: sufficient PRR engagement to initiate co-stimulatory upregulation above what Regular mRNA achieves, but insufficient to trigger translational arrest or apoptotic ISR.

The second immunological advantage of mRNA-OG over Regular mRNA emerges in the bystander population. In both donors, GFP-negative iDCs in both Unfolded and Folded mRNA-OG conditions maintained the CD86 expression required for productive antigen-specific T cell priming and maintenance of an inflammatory phenotype (∼95– 99%), whereas bystander cells in Regular mRNA EP conditions showed marked CD86 suppression (∼55–80% of baseline) (Fig. 5g, right). This paracrine CD86 preservation was equivalent between Folded and Unfolded conformations, indicating that the bystander suppression observed with Regular mRNA is not a function of fold state. Its underlying cause is not resolved by our secretome data, which did not reveal a Regular mRNA-specific increase in any secreted cytokine. This finding was replicated across two donors and in a third independent experimental context in which a post-EP maturation cocktail was applied to all conditions (Fig. 5i, left). In that experiment, bystander GFP-negative DCs in OG-transfected batches reached CD86 levels of 98.6% and 98.1% for

Unfolded and Folded OG respectively, while Regular mRNA bystander cells were suppressed to 63.5% despite receipt of identical maturation stimulus. Strikingly, bystander CD83 was similarly impaired in Regular mRNA bystanders (41.1%) while OG conditions preserved its expression, suggesting that Regular mRNA EP may generate a paracrine suppressive environment that actively antagonizes DC maturation signaling in neighboring untransfected cells, a property absent from both OG conformations (Fig. 5i, right). The per-cell maturation advantage along with the bystander preservation effects define a coherent immunological profile for mRNA-OG. In transfected cells, the structured RNA scaffold appears to provide stronger self-adjuvancy than Regular mRNA, whereas in untransfected neighboring cells, it leaves costimulatory function and maturation responsiveness intact. In a mixed-efficiency EP preparation where transfection of every DC is not achievable, this combination means that an mRNA-OG-loaded batch maximizes both the activation of antigen-carrying cells and the functional availability of bystander DCs for T cell co-stimulation. These data provide preliminary support for immunological advantages compared to traditional unmodified mRNA-loaded DCs that operate independently of per-cell transgene expression level and may be tunable at the level of fold state without altering the encoded antigen.

## Conclusions

We have established a unimolecular mRNA origami platform that integrates an arbitrary protein-coding sequence into a programmable single-stranded RNA architecture without staple oligonucleotides, chemical crosslinks, or co-transcriptional assembly partners. The accompanying sequence design algorithm, paired with the oxView/oxDNA validation workflow, automates the conversion of user-defined mRNA payloads into thermodynamically validated 3D models, providing a rapid in silico-to-bench bridge for downstream design. The resulting EGFP-encoding mRNA-OG construct adopted the predicted rectangular morphology with high monodispersity by AFM and EMSA, and folded reproducibly under defined thermal annealing and buffer conditions.

mRNA-OG is translation-competent and highly deliverable in adherent cell lines, suspension monocytic cells, and primary human moDCs, but results in approximately 20% of the per-cell reporter expression of a typical EGFP-encoded mRNA control across all delivery routes and cell types tested^66–68^. Our reporter transfection efficiency is in-line with independent reports of gene-encoded RNA-OG delivery methods from multiple labs. The mRNA nano-lantern reported by the Hu lab previously demonstrated the best uptake efficiency (∼98% positive by flow cytometry after 2 hours), but uses a non-quantitative readout for transgene expression and a minimal two-staple origami^67^. To our knowledge, every publication encoding EGFP in a real origami architecture either stops at structural characterization^7^, or only reports vague relative fluorescence from plate readers in HeLa cells^67,69,70^. Our work represents the first quantitative comparison of mRNA-OG transgene expression alongside a traditional mRNA format as a rigorous control.

Polysome profiling demonstrated equivalent ribosome loading for Folded and Unfolded mRNA-OG. Preservation of this expression deficit persists across lipofection and electroporation, and across HEK293T, THP-1, iDC, and mDC contexts. Regarding the translation gap of mRNA-OG relative to Regular mRNA, we suspect that our per-cell yield is hampered primarily by the insertion of the scaffolding strand between the stop codon and poly(A) tail, which is likely not optimal for ribosomal termination, cycling, and reinitiation through PABP-dependent mechanisms. We plan to address this hypothesis in our future work through redistribution of structuring responsibilities to other system components.

Orthogonal readouts of ISR activation including IFNβ, eIF2α phosphorylation, ATF4-axis transcript induction, and PKR/RHA knockdown demonstrate that mRNA-OG engages canonical dsRNA-sensing pathways more strongly than Regular mRNA without crossing the threshold into cytotoxic ISR activation. Fold-state differences were observed in individual assays—Folded OG gave the higher IFN-β in THP-1 monocytes and the higher eIF2α phosphorylation in HEK293T cells, whereas Unfolded OG gave the higher IP-10 in iDCs—but these directions were inconsistent across assays and rested on one or two replicates, so no specific sensor is assigned to either conformation. In all cases, eIF2α phosphorylation and GADD34 induction remained subthreshold, allowing translation to proceed and excluding PKR-mediated translational arrest as the operative cause of the OG translation deficit. This subthreshold profile establishes mRNA-OG as a self-adjuvanting payload that is more immunostimulatory than Regular mRNA, without altering the encoded antigen or chemical composition of the construct.

The compact, rigid architecture of the folded nanostructure confers two additional advantages relevant to delivery and manufacturing. First, Folded mRNA-OG complexes with cationic lipid carriers at near-quantitative encapsulation efficiency (∼91%), exceeding both Unfolded mRNA-OG and Regular mRNA, and providing a structure-based purification effect that biases lipoplex formation toward correctly folded transcripts. Second, Folded mRNA-OG retained expression competence and structural integrity over 60 days of refrigerated storage in physiological buffer, suggesting that the programmed dsRNA architecture is at least as stable as conventional mRNA prepared under identical conditions. Our mRNA-OG design platform can express mRNA coding sequences of around 900 nucleotides, and able to accommodate even longer sequences by leaving part of the coding sequence unstructured or using a larger origami template, making the mRNA-OG platform versatile for many applications. Together, these properties address two of the principal logistical constraints of current mRNA therapeutic formats.

Characterization of mRNA-OG in primary human moDCs identified a possible OG-specific innate sensing axis with potential relevance to downstream ex vivo vaccine development. In iDCs, both OG conformations trended toward stronger per-cell CD83 upregulation than Regular mRNA against the electroporation-stress background, while appearing to preserve CD86 expression in untransfected bystander DCs that would otherwise be suppressed by linear mRNA delivery. If confirmed at larger donor numbers, the combination of stronger self-adjuvancy in transfected cells and preserved co-stimulatory function in bystanders would constitute a cellular product-level immunological advantage that operates independently of per-cell transgene expression reflecting the structured scaffold rather than fold state, with direct implications for clinical DC vaccine formats in which uniform transfection across the loaded population cannot be assumed.

Looking forward, the unimolecular architecture of mRNA-OG opens several directions not accessible to linear mRNA or multi-component nanostructure formats.. For example, multimerized mRNA-OG monomers (e.g. by stacking multiple particles using kissing loops integrated into the structuring sequence of mRNA-OG) carrying distinct payloads would enable stoichiometric co-delivery of multiple antigens or signaling components from a single particle, a capability of particular relevance to neoantigen vaccine formats requiring controlled co-loading of multiple patient-specific epitopes^68^. Finally, the compact morphology and regularly patterned surface charge of Folded mRNA-OG suggests that direct coating with cationic peptides, polymers, or other biomolecule-based delivery agents^20,71,23–25^ may be feasible without lipid nanoparticle complexation, offering a route to cold-chain-independent, formulation-simplified delivery that circumvents the limitations of current LNP platforms. Collectively, these directions position mRNA-OG not as a replacement for conventional mRNA but as an orthogonal scaffold whose structural programmability extends the accessible design space of nucleic acid therapeutics.

Situated against the current trajectory of the field, this work reframes the higher-order architecture of an mRNA as a design parameter on equal footing with the sequence-, nucleotide-, and vehicle-level engineering that has defined mRNA therapeutics to date. Where superfolder and structure-stabilized mRNAs remain bound to the coding sequence through synonymous-codon constraints, and where staple-directed and multi-component nanostructures introduce assembly complexity and additional oligonucleotide components, the unimolecular mRNA-OG format achieves a compact, defined fold from a single transcript while leaving the encoded protein untouched. By pairing an automated design pipeline with a quantitative benchmark against a conventional mRNA control, we provide both a reproducible route from arbitrary payload to validated nanostructure and a rigorous baseline against which future structured-mRNA formats can be measured. More broadly, our results argue that translational efficiency, delivery, stability, and innate immune engagement need not be treated as independent problems addressed by separate modifications, but can be co-tuned through a single structural design choice, positioning programmed RNA architecture as a unifying handle for the design of mRNA therapeutics.

## Methods

### Automated mRNA-OG design program

To facilitate the design of single-stranded mRNA origami nanostructures capable of carrying arbitrary coding sequences, we developed an automated Python-based design program (Supplementary Fig. S1a). Building upon the foundational ssRNA origami architectures initially developed by Han et. al.^16^, this algorithm accepts an existing ssRNA origami nanostructure design in oxDNA format and systematically redesigns its nucleotide sequence to incorporate a desired gene. Crucially, the pipeline accomplishes this while preserving the original crossover topology and global structural integrity.

The pipeline requires several key inputs: an initial oxRNA structure file and corresponding topology file, an oxRNA trajectory file (optional but recommended for robust base-pair averaging), the desired mRNA coding sequence, and the sequence definitions for the 5’ and 3’ untranslated regions (UTRs) or tails. The algorithm executes the redesign through the following sequence of operations:

1. **Hydrogen Bond Network Extraction:** First, the algorithm evaluates the provided oxRNA trajectory using the *output_bonds* utility from the oxDNA Analysis Tools (OAT)^27^. By averaging the binding interactions across the trajectory, the script extracts the ideal hydrogen-bonding network, identifying the exact indices of all paired nucleotides that dictate the origami’s predefined shape.
2. **Structural Partitioning and Capacity Validation:** The single-stranded origami structure is divided into two halves: one designated to host the coding sequence and the other to serve as the complementary structural scaffold. The pipeline then verifies that the length of the requested coding sequence does not exceed the nucleotide capacity of the coding half.
3. **Index Mapping and Identification of Self-Binding Regions:** The algorithm isolates the structural indices assigned to the coding sequence and maps them to their paired complementary indices on the scaffold half. Because the target mRNA sequence must remain entirely conserved to yield the correct protein, the algorithm specifically identifies "self-binding" domains within the structure— instances where the origami topology dictates that a segment of the coding sequence must base-pair with another segment of the coding sequence itself.
4. **Sequence Assignment and Strand Complementarity:** The base nanostructure is computationally edited to embed the target mRNA sequence into the designated coding indices. To ensure the origami folds into the correct configuration, the algorithm then modifies the opposing non-coding structural scaffold to be perfectly complementary to the newly embedded mRNA. For the previously identified self-binding coding regions, the sequence is dictated entirely by the desired mRNA sequence, even if this results in local structural mismatches in the final folded state.
5. **Terminal Modifications:** Following the core redesign, the designated 5’ and 3’ UTR sequences are appended and prepended to the appropriate structural termini. The algorithm uses the positional coordinates and backbone vectors of the terminal nucleotides to ensure the newly added tails extend naturally from the origami core.
6. **Validation and Export:** Finally, the algorithm performs comprehensive validation checks to ensure proper complementarity across the structure and evaluates the relative free-energy shifts (calculated based on nearest-neighbor model for RNA^72,73^ for a given base paired segment) of the redesigned structural domains. It outputs new oxRNA topology and configuration files, an interactive annotated oxView file detailing the paired/unpaired states and destabilized domains, and a .dna file with sequence annotations for downstream laboratory visualization in Benchling. By automating these structural calculations, the pipeline ensures that virtually any therapeutic mRNA sequence with a length less than one-half the ssRNA-OG can be rapidly converted into a compact, structurally defined mRNA-OG nanoparticle optimized for physical stability and delivery (see final sequence in Supplementary Table S6).

### In vitro transcription and mRNA origami annealing

mRNA-OG in vitro transcription was performed using TriLink CleanScript kit without synthetic nucleotides, substituting an equimolar quantity of CleanCap AG rather than the included 3-MeO cap derivative. For origami constructs only, IVT was performed at 30.5°C to promote polymerase processivity in difficult template regions. Unstructured templates were incubated at 37°C. All incubations were carried out for 5 hours followed by supplementation of the mixture to 2 mM CaCl_2_ and 2 µL of TurboDNase incubated at 37°C for 30 min. Reactions were then purified using Qiagen RNeasy spin columns.

To thermally anneal mRNA-OG into the folded state, the mRNA was diluted to 100 ng/µL (between 50-500 ng/µL was seen to be an acceptable range as shown in Supplementary Figs. S4b, S5) in 1X PBS supplemented with an additional 0.25 M NaCl. Annealing reactions were heated to 85°C and cooled at −1°C/min down to 60°C, followed by cooling at −1°C/5 min between 65-45°C to allow for optimal annealing efficiency, followed by −1°C/min from 45-25°C and then immediately cooled to and stored at 4°C for up to 1 week prior to transfection.

Alternative annealing conditions over a wide range of biological buffers and salt conditions (0-1 M NaCl) (Supplementary Fig. S4) were tested and the optimized conditions were selected based on fold state and nanoparticle monodispersity evaluated via AFM.

### Atomic Force Microscopy

The RNA sample was diluted to 10 ng/µL with 1X TAE + 6 mM Mg^2+^ buffer to 15 µL volume total and deposited onto a freshly cleaved mica surface. RNA solution was incubated on mica for 3 min. 35 µL of 1X TAE + 6 mM NiCl_2_ buffer was then added to the mica. Measurements were performed using ScanAsyst Fluid mode with ScanAsyst-Fluid+ probes (Bruker) using MultiMode 8. AFM images were processed with the Nanoscope Analysis application.

### Agarose Gel Electrophoretic Mobility Shift Assay

1 µg of RNA was loaded onto a 1% (w/v) agarose gel (1X TAE buffer) pre-stained with GelRed at 90 V for 90 min and visualized with a Bio-Rad Gel Doc XR+ imager.

### Lipofection into Suspended Cell Lines

HEK293T cells were transfected at 6 million cells/dish into a 15-cm dish with 0.75 µL Lipofectamine MessengerMAX Transfection Reagent (Thermo Fisher Scientific) and 20 µg mRNA each, prepared according to manufacturer’s instructions.

### Polysome Profiling of Transfected mRNA-OG

1. **Buffers.** Polysome resuspension (RES) buffer contained 20 mM Tris-HCl (pH 7.5), 10 mM MgCl_2_, 150 mM KCl, 100 µg mL^-1^ cycloheximide (CHX), 100 µM DTT, and 0.2 U µL^-1^ SUPERase-In RNase Inhibitor (Invitrogen). Lysis (LYS) buffer was supplemented with 5 mM CaCl_2_, 1% (v/v) Triton X-100, and 10 U mL^-1^ TURBO DNase (Invitrogen). Sucrose buffers (10%, 20%, 30%, 40%, and 50% w/v) were prepared by dissolving RNase-free sucrose (Invitrogen) in RES buffer.
2. **Sucrose gradient preparation.** Linear 10–50% (w/v) sucrose gradients were prepared in open-top polyallomer ultracentrifuge tubes (Beckman, SW40 Ti compatible) by the freeze-thaw layering method. Briefly, 2 mL of each sucrose concentration was sequentially layered from highest to lowest density, with each layer frozen at −80°C for 10 min before the next was added. Completed gradients were equilibrated at 4°C overnight prior to loading.
3. **Lysate preparation.** At 16 h post-transfection, the medium was replaced with pre-warmed complete medium containing 100 µg mL^-1^ CHX, and cells were incubated for 3 min at 37°C to stall translating ribosomes. Plates were rapidly washed with ice-cold 1X PBS containing 100 µg mL^-1^ CHX, and all subsequent steps were performed on ice. Cells were lysed in situ with 1.5 mL of ice-cold LYS buffer per dish for 3 min, scraped, and transferred to pre-chilled microcentrifuge tubes. Lysates were clarified at 12,000 × g for 10 min at 4°C, and the supernatant was retained. For EDTA-mediated polysome collapse controls, 0.5 M EDTA (pH 8.0) was added to clarified lysate to a final concentration of 20 mM, followed by incubation on ice for 10 min. A_260_ of each clarified lysate was measured by NanoDrop.
4. **Ultracentrifugation and fractionation.** Five to ten A_260_ units of clarified lysate (or the maximum loadable volume, up to ∼800 µL) were gently layered onto pre-chilled gradients. Tubes were balanced using LYS buffer and centrifuged in a Beckman SW40 Ti rotor at 210,000 × g for 2 h at 4°C with maximum acceleration and unbraked deceleration. Gradients were fractionated by bottom puncture using a fine-gauge needle, collecting 48 sequential fractions of ∼250 µL into pre-chilled tubes; fractions were subsequently renumbered from the top (fraction 1) to the bottom (fraction 48) of the gradient. A_254_ was measured for each fraction by NanoDrop against a RES buffer blank to generate the polysome absorbance profile with baseline subtraction at 340 nm.
5. **RNA extraction from fractions.** Fractions were pooled according to the A_254_ profile, typically grouping ∼3-7 fractions per pool spanning the free RNA, 40S/60S, monosome, light polysome, and heavy polysome regions, with equal µL contributed from each constituent fraction to reach a total pool volume of 250 µL. Pools were mixed with TRIzol-LS (Invitrogen) at a 1:4 sample-to-reagent volume ratio, inverted 15 times, incubated for 5 min at room temperature, and snap-frozen in liquid nitrogen. After thawing on subsequent day, phase separation was performed by addition of chloroform (0.2 volumes per volume of TRIzol-LS), inversion, 3 min incubation at room temperature, and centrifugation at 12,000 × g for 15 min at 4°C. The aqueous phase was precipitated with isopropanol (0.67 volumes per volume of TRIzol-LS) for 10 min at room temperature, pelleted at 12,000 × g for 10 min at 4°C, washed with an equivalent volume of isopropanol, centrifuged at 7,500 × g for 5 min at 4°C, washed with 70% EtOH twice to remove residual phenol, and air-dried for 5–10 min. RNA pellets were resuspended in 30 µL of nuclease-free water at 60°C for 10–15 min and quantified using NanoDrop or Qubit RNA HS Assay (Invitrogen).
6. **Transcript quantification across fractions.** Transcript-specific abundance in each pool was measured by one-step RT-qPCR using the Luna Universal One-Step RT-qPCR Kit (NEB) according to the manufacturer’s instructions, with primers specific to the mRNA-OG structuring sequence. Standard curves were generated from serial dilutions of in vitro transcribed template. Polysome-to-monosome (P:M) ratios were calculated from the integrated transcript abundance in polysome pools relative to the monosome pool.

### Encapsulation Efficiency Assay

Encapsulation efficiency of EGFP reporter mRNAs formulated with Lipofectamine MessengerMAX was quantified using a SYBR Gold dye-exclusion assay. The assay was performed to determine whether transcript structuring altered the fraction of RNA protected within lipid-based nanoparticles prior to transfection into HEK293T cells.

Briefly, mRNA–MessengerMAX particles were prepared using a fixed formulation ratio of 1.5 µL MessengerMAX per 1 µg RNA. Formulations were diluted 1:50 prior to analysis. The assayed constructs included Regular EGFP mRNA, mRNA-OG Unfolded RNA, and mRNA-OG Folded RNA. The mRNA-OG constructs were 2221 nt in length, whereas the regular EGFP mRNA was 996 nt.

RNA accessibility was measured in flat-bottom 96-well plates using SYBR Gold fluorescence. For each RNA identity, standards containing 0, 1, 2, 5, 10, or 20 ng RNA per well were included in duplicate. LNP samples were measured in quadruplicate. Fluorescence was first measured before detergent treatment to quantify dye-accessible RNA. Triton X-100 was then added to each well to a final concentration of 0.1% to disrupt the nanoparticles and expose total RNA, after which fluorescence was measured again from the same wells. Pre- and post-Triton values were background-corrected using the corresponding dye-only and dye-plus-Triton blank wells, respectively.

Encapsulation efficiency was calculated for each well as:

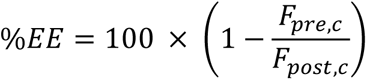

where F_pre,c_ is the background-corrected fluorescence before Triton treatment and F_post,c_ is the background-corrected fluorescence after Triton treatment. Thus, the assay reports the fraction of total RNA that was inaccessible to SYBR Gold prior to detergent lysis. Values were summarized as mean ± SD across four technical replicate wells.

Post-Triton standard curves were generated independently for each RNA identity to evaluate RNA-specific fluorescence response and assay linearity. Because encapsulation efficiency is calculated as a same-well ratio of dye-accessible to total RNA for each RNA species, conversion from RNA mass to RNA moles does not alter the percentage encapsulation value. For absolute molar estimates, RNA mass was converted using an average molecular weight of 340 g mol^-1^ nt^-1^ and the corresponding transcript length. Thus, %EE was interpreted as the fraction of RNA molecules protected from dye prior to lysis, rather than the fraction of lipid particles containing RNA or the number of RNA molecules per particle.

### PKR-Dependent Translation in A549 Cell Lines

RHA and PKR-deficient A549 cell lines were a generous donation from Junior Ayuk-Enow^74,75^. Lipofection and flow cytometry was carried out as described for all other HEK293T-based assays.

### IFN-ꞵ and IL-6 ELISA from Cell Culture Supernatant

1 mL of unconditioned immature DC culture media was cryopreserved on day 0, and conditioned media supernatant was collected and cryopreserved on days 5 (prior to processing cells for electroporation), and 7 (during harvest, 48 hours post-electroporation). Upon thawing and brief centrifugation at 10,000 × g for 2 min, 50 μL of supernatant was used as the input to an IFN-β Quantikine QuickKit ELISA (R&D Systems, #QK410) directly according to the manufacturer’s protocol.

### EIF2α Phosphorylation Western Blot

Cell lysates were prepared from HEK293T cells 24 h after transfection. Cells were centrifuged (300 RCF, 5 min) and washed twice with ice-cold PBS. Cell pellets were lysed in M-PER Mammalian Protein Extraction Reagent (Thermo Fisher Scientific, #78501) with HALT Protease Inhibitor Cocktail (Thermo Fisher Scientific, #78430), and incubated on ice for 20 min. After centrifugation (16,000 × g, 15 min, 4°C), a part of the supernatant was taken for protein concentration assay using the Qubit Protein and Protein Broad Range (BR) Assay (Thermo Fisher Scientific, #A50668). Samples were then mixed with Laemmli’s loading buffer and heated at 95°C for 5 min. Equal amount of protein extracts normalized against cell count was applied to sodium dodecyl sulphate-polyacrylamide gel electrophoresis and transferred into a PVDF membrane using Trans-Blot SD Semi-Dry Transfer Cell (Bio-Rad) according to the manufacturer’s protocol. Following transfer, membranes were incubated with specific primary antibodies. Anti-EIF2s1 total (Thermo Fisher Scientific, #82936-1-RR) and Anti-phospho EIF2α Ser52 (Thermo Fisher Scientific, #MA5-15133) were used at 8,000-fold dilution and 1,000-fold dilution, respectively. Beta-Actin antibody (Proteintech, #20536-1-AP) was used at 10,000-fold dilution. Then, the blot was incubated with secondary antibodies. Goat Anti-Rabbit IgG (H+L)-HRP conjugate (Proteintech, #SA00001-2) were used at 10,000-fold dilution. The blots were detected with SuperSignal West Pico PLUS Chemiluminescent Substrate (Thermo Scientific, #34580) using iBright FL1500 Imaging System (Invitrogen). Protein expression level was calculated from bands intensities with ImageJ (NIH).

### Storage Stability of mRNA-OG

RNA origami samples were prepared as described above. “Regular mRNA” sample was prepared using the same EGFP full-length mRNA sequence as TriLink Biotechnologies. All mRNA samples were prepared at 100 ng/µL in 1X Phosphate-Buffered Saline (Corning) supplemented with 0.25 M NaCl (NaCl, nuclease-free H_2_O). Samples were briefly vortexed, spun down, aliquotted into predetermined tubes for each timepoint, and left to incubate at 4°C, protected from light. At each timepoint, the RNA stored at 4°C from the allotted tube was characterized by AFM, agarose gel EMSA, and lipofection followed by flow cytometry on a Thermo Fisher Scientific Attune NxT using a singlets gating strategy (Supplementary Fig. S8).

### Dendritic Cell Culture & mRNA Electroporation

Dendritic cells were cultured as described previously^76^. Specifically, peripheral blood mononuclear cells (PBMCs) were isolated from healthy human donor leukopheresis products by Ficoll-Paque gradient centrifugation. Human immature dendritic cells were generated via immunomagnetic selection of CD14+ monocytes (CD14 MicroBeads, human, #130-097-052, Miltenyi Biotec) using an AutoMACS Pro Separator (Miltenyi Biotec) and cultured for 5 days in CellGenix GMP DC Media supplemented with 1000 IU/mL rhGM-CSF and 1000 IU/mL rhIL-4. Cultures were maintained at 37°C in a humidified atmosphere containing 5% CO₂. All cytokines were purchased from R&D Systems.

On day 5, immature DCs were washed and resuspended in Electroporation Buffer (MaxCyte) with 800 U/mL RNaseOUT (Invitrogen) at 5×10^7^ cells/mL. Immediately prior to electroporation, 235 pM mRNA was added to each 25 µL sample. Mock transfected samples were alternatively loaded with a volume of electroporation buffer equivalent to the volume of RNA loaded into other samples. Each sample was then loaded into individual wells of OC-25×3 processing assemblies (with the exception of the “No EP” negative controls) and electroporated with a MaxCyte ATx electroporator using the pre-loaded THP-1 program. Alternatively, iDC were supplemented on day 5 with 1000 IU/mL rhTNFα and 0.35 µg/mL Prostaglandin E_2_ (PGE_2_, Sigma-Aldrich) and matured for 48 hours. On day 7, mature DC were electroporated as above using the pre-loaded DC-2 program.

After electroporation, cells were transferred to a 24-well plate each and matured in 1 mL CellGenix GMP DC Media supplemented with 1000 IU/mL rhGM-CSF, 1000 IU/mL rhIL-4, 1000 IU/mL rhTNFα, and 0.35 µg/mL PGE_2_ (Invitrogen) for 48 hours. GFP fluorescence was monitored at 24 and 48 hours using fluorescence microscopy and flow cytometry (at 48 h only). Dendritic cells were phenotypically characterized at days 5 and 7 via flow cytometry on a Beckman Coulter CytoFlex.

### Innate Immune Secretome Analysis

Conditioned media supernatants were collected 48 hours post-electroporation from all experimental conditions. For the iDC arm, this corresponded to day 7 of culture; for the mDC arm, day 9. Non-electroporated negative controls (No EP) were maintained in parallel under identical media conditions and collected at the corresponding timepoints. A single unconditioned iDC differentiation media sample was collected on day 0 as a background reference. All supernatants were immediately aliquoted and cryopreserved at −80°C until analysis.

Supernatant cytokine and chemokine concentrations were quantified using a Bruker IsoPlexis IsoLight system with a CodePlex Adaptive Immune — Human chip. For each sample, 5.5 µL of thawed supernatant was loaded per well; chips were processed and imaged on the IsoLight instrument and analyzed using IsoSpeak software (Bruker) with concentrations interpolated from on-chip standard curves. Analytes reflecting exogenous media components (TNFα, IL-4, GM-CSF) were excluded from analysis. Analytes below the limit of detection across all conditions, or exhibiting technically anomalous signal patterns were excluded from primary analyses.

## Data Availability

The source data underlying the main-text and Extended Data figures are provided with this paper. The oxRNA structure, topology, and trajectory files, the annotated oxView design files, and the full nucleotide sequences for all mRNA-OG constructs and reporter controls used in this study have been deposited in Zenodo and are publicly available at 10.5281/zenodo.21657417. Uncropped agarose gel and immunoblot images, atomic force microscopy fields, flow cytometry data files (.fcs), polysome profiling traces, RT-qPCR records, and multiplex cytokine/ELISA datasets are provided within the paper, its Supplementary Information, and the associated Zenodo repository deposit at 10.5281/zenodo.21657417. Any additional data supporting the findings of this study are available from the corresponding authors upon reasonable request. Biological materials, including in vitro transcription templates and mRNA-OG constructs, are available from Petr Šulc under a standard material transfer agreement.

## Code Availability

The custom Python mRNA-OG sequence-design pipeline (mRNA-Design), which redesigns a paranemic single-stranded RNA origami to encode an arbitrary coding sequence, together with the example scripts used to generate the results reported here, is available at http://github.com/mlsample/mRNA-Design and archived at 10.5281/zenodo.21657417 (release v0.1.0). The pipeline calls the oxDNA Analysis Tools (OAT) for hydrogen-bond network extraction and NUPACK for nearest-neighbor free-energy evaluation; oxDNA, oxView, OAT, and NUPACK are available from their respective developers under their own licenses. mRNA-Design is released under the GPL-3.0 license.

## Acknowledgments

M.P.G. and K.S.A acknowledge support from Mayo Clinic Breast Cancer SPORE grant P50 CA116201 from the National Institutes of Health. M.P.G. and P. Š. acknowledge support from the 2025 Arizona State University-Mayo Clinic Alliance for Healthcare SEED program. P. Š. additionally acknowledges funding from the National Science Foundation DMR 2239518. Research reported in this publication was supported by The National Institute of General Medical Sciences of the National Institutes of Health under grant number 1R01GM145916-01A1 to P. Š and N.S., and grant number DP2GM132931 to N.S. The content is solely the responsibility of the authors and does not necessarily represent the official views of the National Institutes of Health. N.S. gratefully acknowledges Steve Huffman for a generous gift supporting this research.

## Appendix 1. List of Extended Data Figures

**Extended Data Fig. 1.**
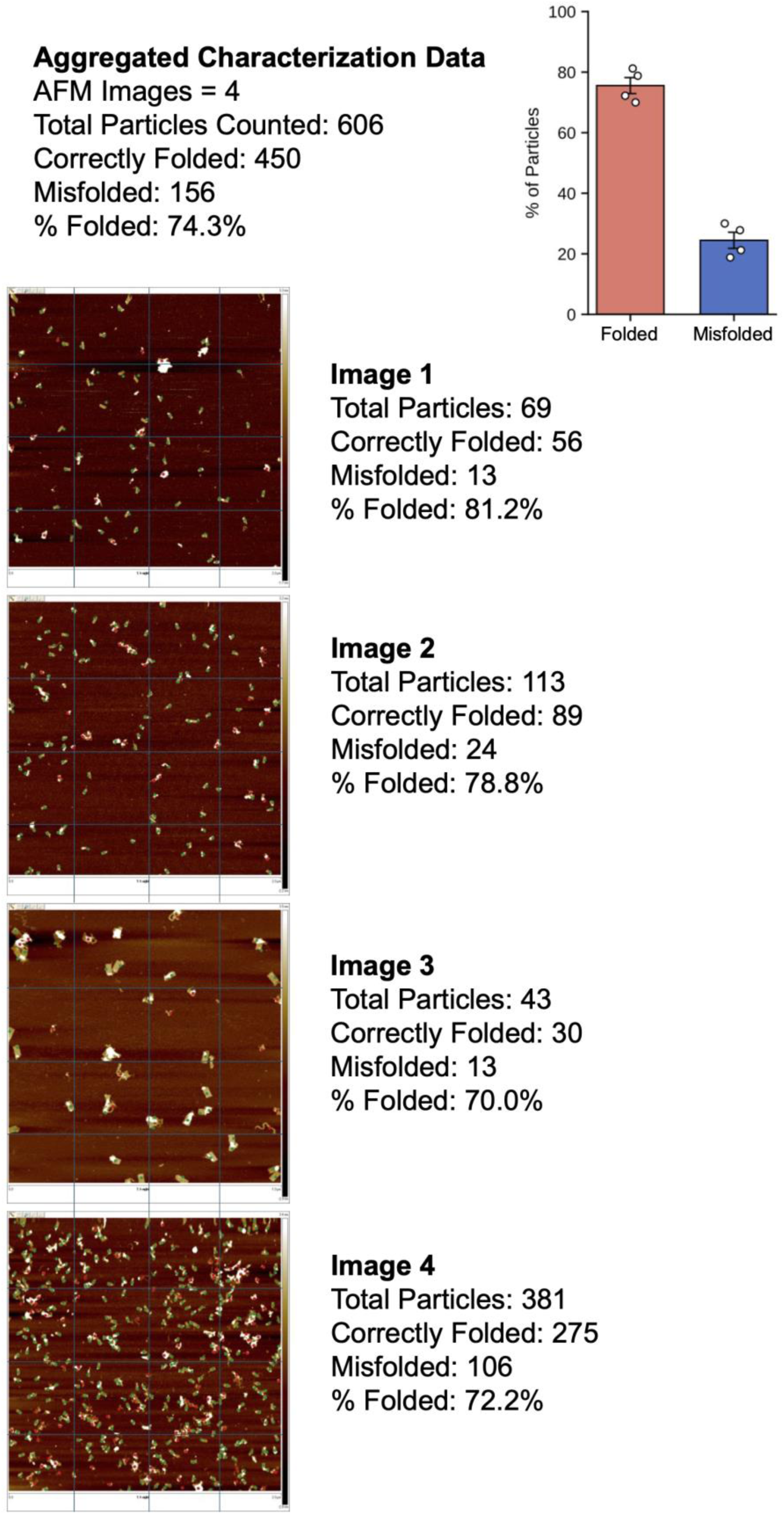
Quantitative nanoparticle fold state characterization via Atomic Force Microscopy. 4 AFM fields spanning either 2.5 μm or 1.0 μm were selected from independent folding reactions on independent days and split in 16 quadrants each. Particles were manually characterized as Folded or Misfolded based on if they met the following criteria: (1) distinguishable dsRNA with at least 2 defined edges with a 90 degree corner, (2) absence of ssRNA with an apparent length greater than 2× the nanoparticles longest visible edge. For aggregated clusters, the visible nanoparticles also had to: (3) be visibly distinguishable.

**Extended Data Fig. 2.**
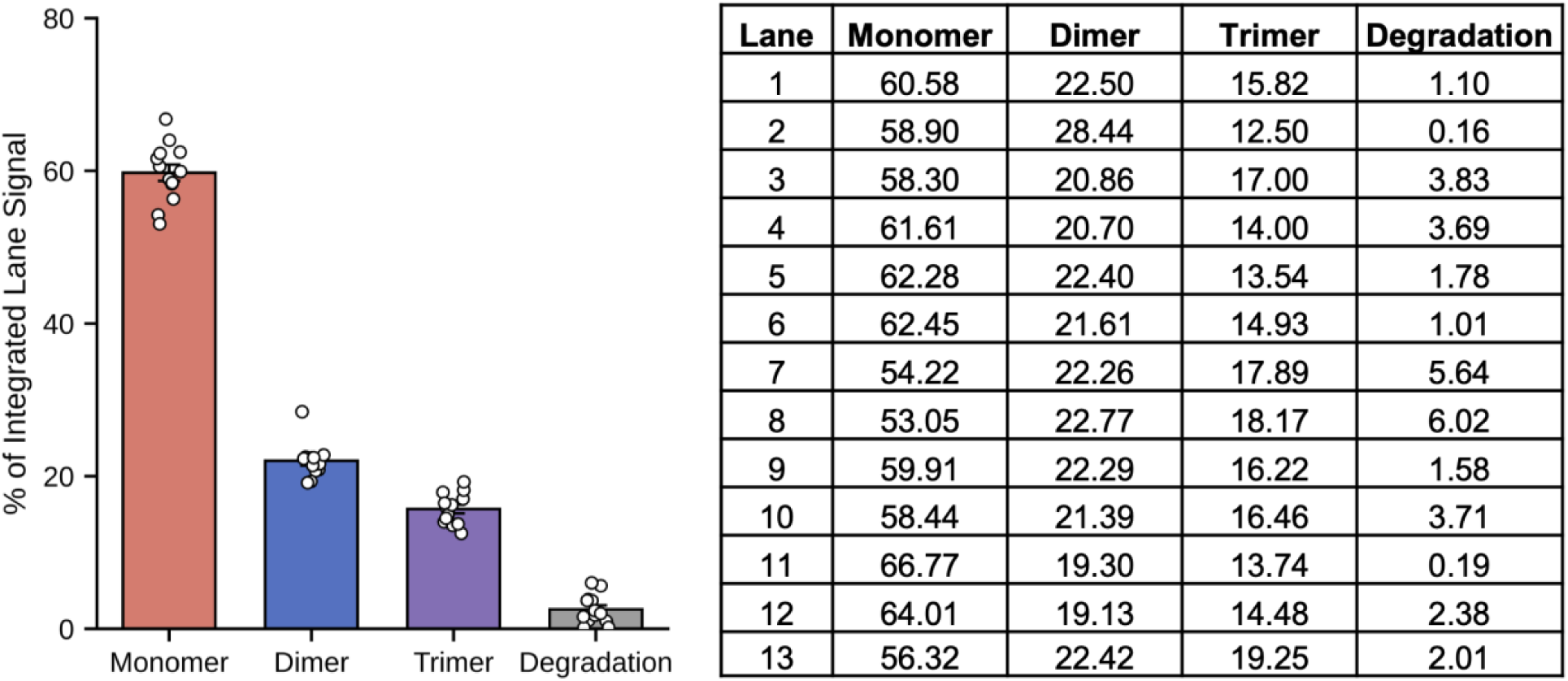
Quantitative nanoparticle monodispersity and orthogonal monodispersity via agarose gel EMSA. 13 independent folding reactions from 3 independent IVT reactions conducted under identical conditions were annealed using the fast-anneal protocol (see Methods) in 1X PBS + 0.25 M NaCl at a concentration of 100 ng/μl. 1 μg (10 μL) of each annealing reaction was then loaded onto a 1% TAE-agarose gel and electrophoresed for 60 minutes at 80 V. Bands were quantified with ImageJ as total area under curve for each lane’s profile with baseline subtraction. Monomers, dimers, and trimers were quantified as individual peaks, while degradation was considered to be all signal above baseline with a higher electrophoretic mobility than the monomer band.

**Extended Data Fig. 3.**
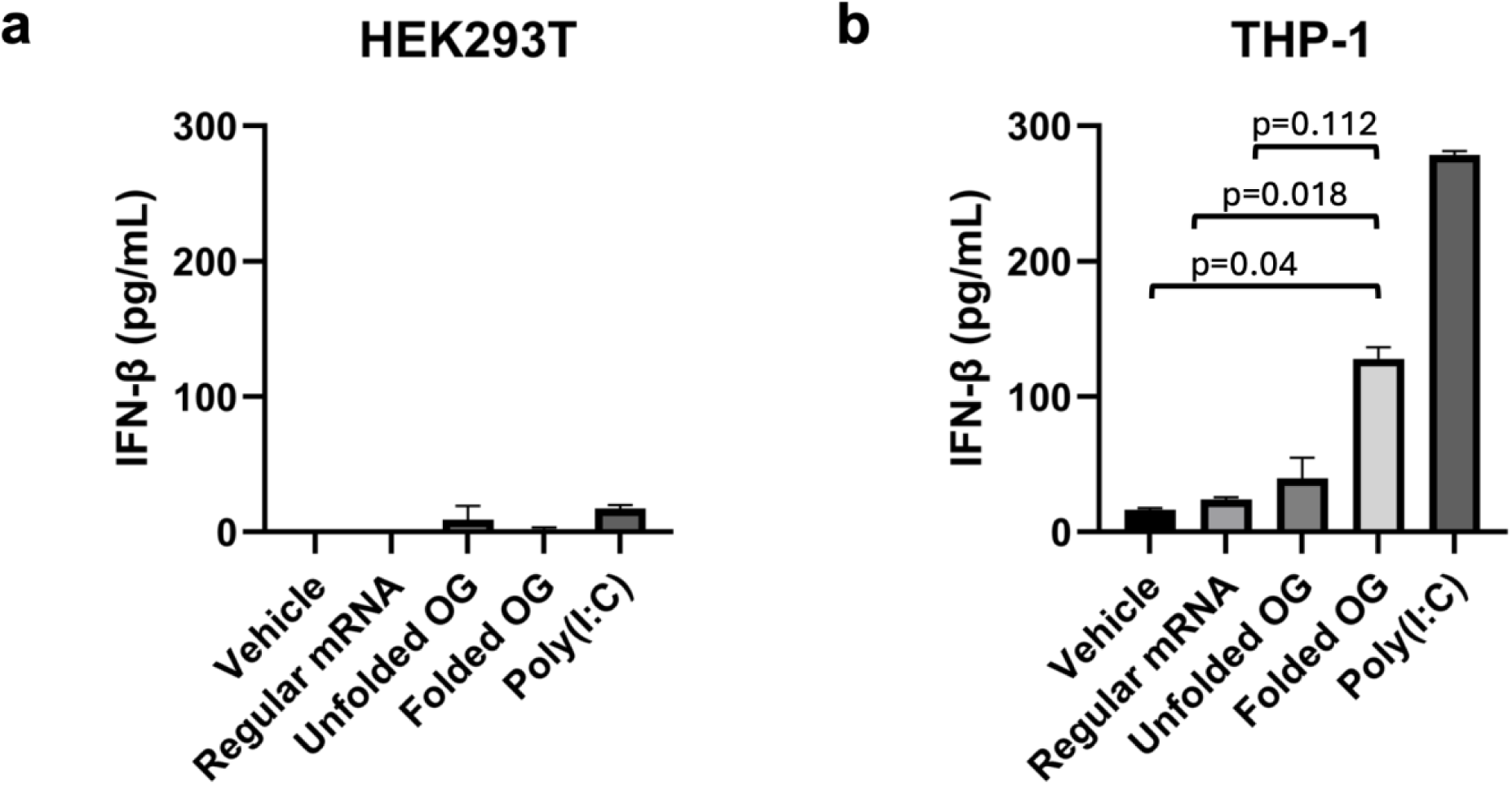
Quantification of IFN-β secreted by HEK293T and THP-1 in response to Vehicle, Regular mRNA, Unfolded mRNA-OG, Folded mRNA-OG, or Poly(I:C). HEK293T cells did not secrete levels above the LLOD (lower limit of detection) for IFN-β in any condition, including the positive control, consistent with reports of deficient RIG-I / MDA5 signaling and STING deficiency. THP-1 cells responded strongly (200-300 pg/mL) to positive control poly(I:C) and Folded mRNA-OG, with a less robust response to Unfolded mRNA-OG and Regular mRNA. No response above the LLOD was observed for lipofectamine vehicle in this study.

**Extended Data Fig. 4.**
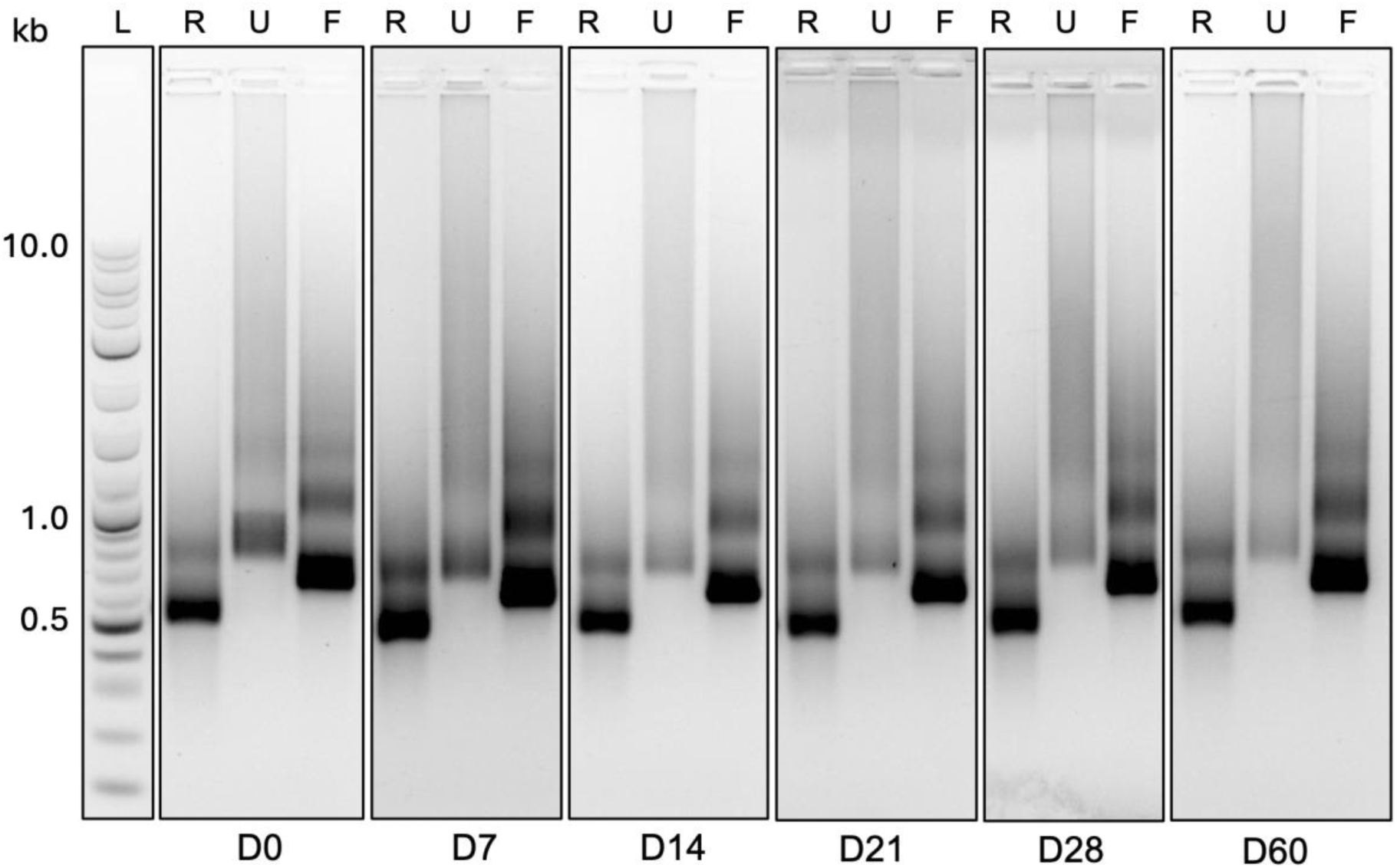
Agarose gel integrity of Regular EGFP mRNA (R), Unfolded mRNA-OG (U), and Folded mRNA-OG (F) Longevity over 60 days of storage at 4°C. 1 μg (10 μL) of each mRNA stored at 100 ng/μL in annealing buffer (1X PBS + 0.25 M NaCl) was electrophoresed on a 1% TAE-agarose gel under identical conditions at 0, 7, 14, 21, 28, and 60 days after IVT and purification without RNase inhibitor present. No bands show apparent degradation in size over the time period studied, however the Unfolded mRNA-OG mRNA collapsed from a broad band into a less intense, faster traveling band, aligning with our AFM data demonstrating collapse of the unfolded to a near-folded state (its energetic minimum) over the study period.

**Extended Data Fig. 5.**
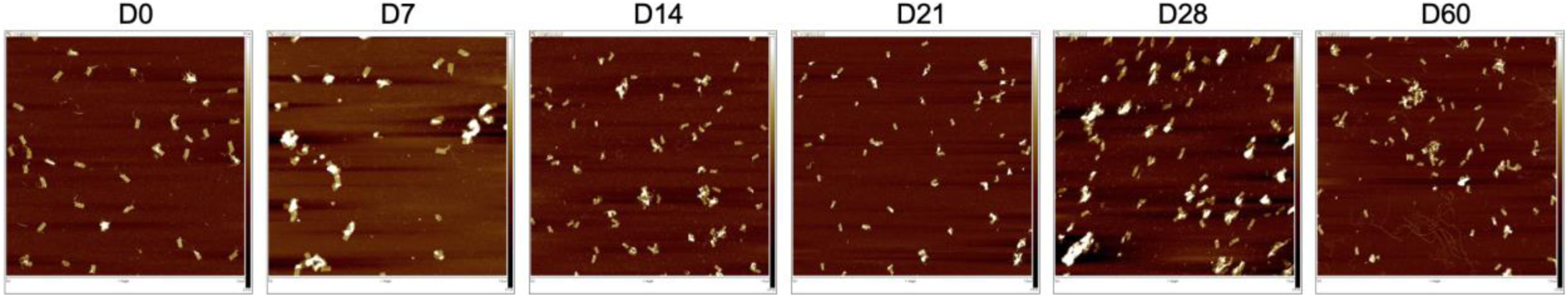
AFM study of Folded mRNA-OG nanoparticle structure longevity over 60 days at 4°C. The same samples from Extended Data Fig. 4 were subjected to AFM at each longevity timepoint to study their structural stability. While the structural integrity of the nanoparticles is consistent with the gel EMSA study, by day 60 nanoparticle aggregation via UTR cross-hybridization is evident without apparent OG unfolding.

**Extended Data Fig. 6.**
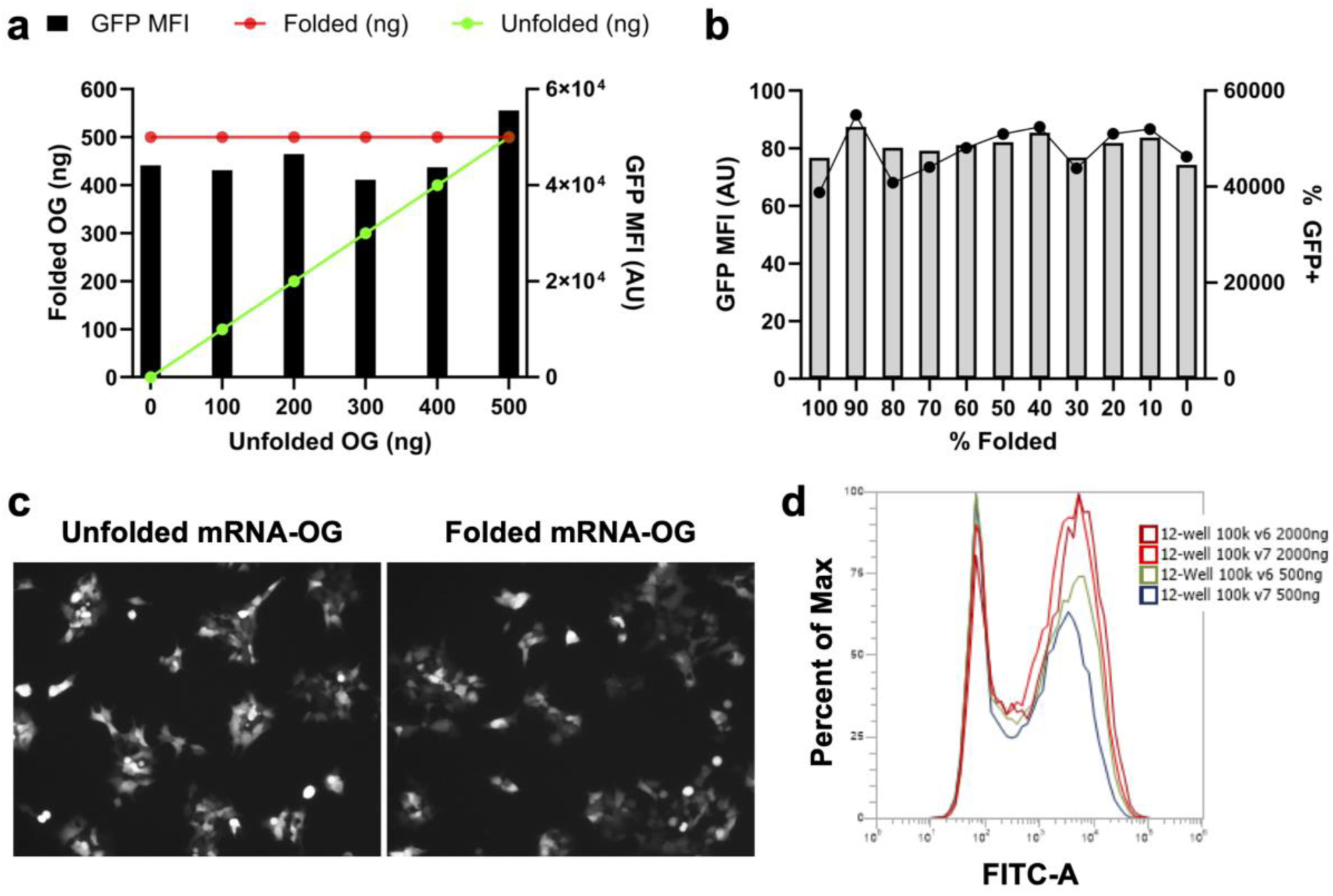
Validation of equivalent Folded/Unfolded mRNA-OG performance in lipofected HEK293T cells. **a,** Cross-titration of thermally annealed (Folded) vs annealing-naive (Unfolded) mRNA-OG from the same IVT preparation were combined in 1X PBS supplemented with 0.25 M NaCl and lipofected at 1 μg of RNA / 100k HEK293T cells without appreciable differences in transfection efficiency or EGFP MFI at 24 hours. **b,** We next added increasing amounts of Unfolded mRNA-OG to a constant 500 ng of Folded and noted that the transfection was probably saturated due to a lack of change in MFI at 24 hours. **c,** Fluorescence micrographs of HEK293T cells from (a) demonstrating roughly equivalent EGFP expression between Unfolded and Folded samples. **d,** Cells were lipofected with either 2 μg or 500 ng of mRNA-OG with either synthetic (v6) or murine β-globin (v7) UTRs and EGFP reporter expression was measured at 24 hours post-transfection via flow cytometry (n=1). Increased input mass yields higher transfection efficiency but not higher MFI, consistent with expression being capped intracellularly by ribosomal processivity or innate immune signaling instead of copy number delivered.

## Notes

### Competing Interest Statement

The authors have declared no competing interest.

## References

1. Sahin, U., Karikó, K. & Türeci, Ö. mRNA-based therapeutics — developing a new class of drugs. Nat Rev Drug Discov 13, 759–780 (2014).

2. Kulkarni, J. A. et al. The current landscape of nucleic acid therapeutics. Nat. Nanotechnol. 16, 630–643 (2021).

3. Li, S. et al. Payload distribution and capacity of mRNA lipid nanoparticles. Nat Commun 13, 5561 (2022).

4. Dzananovic, E., McKenna, S. A. & Patel, T. R. Viral proteins targeting host protein kinase R to evade an innate immune response: a mini review. Biotechnol Genet Eng Rev 34, 33–59 (2018).

5. Mulroney, T. E. et al. N1-methylpseudouridylation of mRNA causes +1 ribosomal frameshifting. Nature 625, 189–194 (2024).

6. Rothemund, P. W. K. Folding DNA to create nanoscale shapes and patterns. Nature 440, 297–302 (2006).

7. Parsons, M. F. et al. 3D RNA-scaffolded wireframe origami. Nat Commun 14, 382 (2023).

8. Seitz, I. et al. Folding of mRNA-DNA Origami for Controlled Translation and Viral Vector Packaging. Advanced Materials 37, 2417642 (2025).

9. Wayment-Steele, H. K. et al. Theoretical basis for stabilizing messenger RNA through secondary structure design. Nucleic Acids Res 49, 10604–10617 (2021).

10. Zhang, H. et al. Algorithm for optimized mRNA design improves stability and immunogenicity. Nature 621, 396–403 (2023).

11. Leppek, K. et al. Combinatorial optimization of mRNA structure, stability, and translation for RNA-based therapeutics. Nat Commun 13, 1536 (2022).

12. Geary, C., Rothemund, P. W. K. & Andersen, E. S. A single-stranded architecture for cotranscriptional folding of RNA nanostructures. Science 345, 799–804 (2014).

13. Geary, C., Grossi, G., McRae, E. K. S., Rothemund, P. W. K. & Andersen, E. S. RNA origami design tools enable cotranscriptional folding of kilobase-sized nanoscaffolds. Nat. Chem. 13, 549–558 (2021).

14. Tran, M. P. et al. Genetic encoding and expression of RNA origami cytoskeletons in synthetic cells. Nat. Nanotechnol. 20, 664–671 (2025).

15. Vallina, N. S., McRae, E. K. S., Geary, C. & Andersen, E. S. An RNA origami robot that traps and releases a fluorescent aptamer. Science Advances 10, eadk1250 (2024).

16. Han, D. et al. Single-stranded DNA and RNA origami. Science 358, eaao2648 (2017).

17. Qi, X. et al. Programming molecular topologies from single-stranded nucleic acids. Nat Commun 9, 4579 (2018).

18. Qi, X. et al. RNA Origami Nanostructures for Potent and Safe Anticancer Immunotherapy. ACS Nano 14, 4727–4740 (2020).

19. Dai, K. et al. Edge Length-Programmed Single-Stranded RNA Origami for Predictive Innate Immune Activation and Therapy. J. Am. Chem. Soc. 145, 17112– 17124 (2023).

20. Ponnuswamy, N. et al. Oligolysine-based coating protects DNA nanostructures from low-salt denaturation and nuclease degradation. Nat Commun 8, 15654 (2017).

21. Yip, T., Qi, X., Yan, H. & Chang, Y. RNA Origami Functions as a Self-Adjuvanted Nanovaccine Platform for Cancer Immunotherapy. ACS Nano 18, 4056–4067 (2024).

22. Yip, T., Tu, X., Qi, X., Yan, H. & Chang, Y. Adjuvanted RNA Origami—A Tunable Peptide Assembly Platform for Constructing Cancer Nanovaccines. Vaccines (Basel) 13, 560 (2025).

23. Smolková, B. et al. Protein Corona Inhibits Endosomal Escape of Functionalized DNA Nanostructures in Living Cells. ACS Appl. Mater. Interfaces 13, 46375–46390 (2021).

24. Elblová, P. et al. Peptide-coated DNA nanostructures as a platform for control of lysosomal function in cells. Chem Eng J 498, 155633 (2024).

25. Elblová, P. et al. Geometrically constrained cytoskeletal reorganisation modulates DNA nanostructures uptake. J. Mater. Chem. B 13, 2335–2351 (2025).

26. Fornace, M. E. et al. NUPACK: Computational Nucleic Acid Analysis and Design. ACS Synth. Biol. 15, 1426–1441 (2026).

27. Poppleton, E. et al. Design, optimization and analysis of large DNA and RNA nanostructures through interactive visualization, editing and molecular simulation. Nucleic Acids Research 48, e72 (2020).

28. Bohlin, J. et al. Design and simulation of DNA, RNA and hybrid protein–nucleic acid nanostructures with oxView. Nat Protoc 17, 1762–1788 (2022).

29. Ma, D. et al. Multi-arm RNA junctions encoding molecular logic unconstrained by input sequence for versatile cell-free diagnostics. Nat. Biomed. Eng 6, 298–309 (2022).

30. Chassé, H., Boulben, S., Costache, V., Cormier, P. & Morales, J. Analysis of translation using polysome profiling. Nucleic Acids Res 45, e15 (2017).

31. Panda, A. C., Martindale, J. L. & Gorospe, M. Polysome Fractionation to Analyze mRNA Distribution Profiles. Bio Protoc 7, e2126 (2017).

32. Pringle, E. S., McCormick, C. & Cheng, Z. Polysome Profiling Analysis of mRNA and Associated Proteins Engaged in Translation. Current Protocols in Molecular Biology 125, e79 (2019).

33. Svitkin, Y. V. et al. N1-methyl-pseudouridine in mRNA enhances translation through eIF2α-dependent and independent mechanisms by increasing ribosome density. Nucleic Acids Res 45, 6023–6036 (2017).

34. Svitkin, Y. V., Gingras, A.-C. & Sonenberg, N. Membrane-dependent relief of translation elongation arrest on pseudouridine- and N1-methyl-pseudouridine-modified mRNAs. Nucleic Acids Res 50, 7202–7215 (2022).

35. Rozman, B. et al. N1-Methylpseudouridine directly modulates translation dynamics. Nature 651, 533–541 (2026).

36. Ivanov, A. et al. PABP enhances release factor recruitment and stop codon recognition during translation termination. Nucleic Acids Res 44, 7766–7776 (2016).

37. Mangkalaphiban, K. et al. Extended stop codon context predicts nonsense codon readthrough efficiency in human cells. Nat Commun 15, 2486 (2024).

38. Alekhina, O. M., Terenin, I. M., Dmitriev, S. E. & Vassilenko, K. S. Functional Cyclization of Eukaryotic mRNAs. International Journal of Molecular Sciences 21, 1677 (2020).

39. Lai, W.-J. C. et al. Intrinsically Unstructured Sequences in the mRNA 3ʹ UTR Reduce the Ability of Poly(A) Tail to Enhance Translation. Journal of Molecular Biology 434, 167877 (2022).

40. Afonina, Z. A. & Vassilenko, K. S. Circularization and Ribosome Recycling: From Polysome Topology to Translational Control. International Journal of Molecular Sciences 27, 1251 (2026).

41. Salman, A. et al. Dual role of 3’ UTR length in modulating translation termination efficiency. RNA 32, 1129–1146 (2026).

42. Lama, L., Morozov, P., Garzia, A. & Tuschl, T. Dissection of innate-immune-ligand- and interferon-protein-mediated transcriptional responses in human THP1 cell states. Commun Biol 9, 239 (2026).

43. Sun, L., Wu, J., Du, F., Chen, X. & Chen, Z. J. Cyclic GMP-AMP Synthase Is a Cytosolic DNA Sensor That Activates the Type I Interferon Pathway. Science 339, 786–791 (2013).

44. Ishikawa, H. & Barber, G. N. STING is an endoplasmic reticulum adaptor that facilitates innate immune signalling. Nature 455, 674–678 (2008).

45. Reus, J. B., Trivino-Soto, G. S., Wu, L. I., Kokott, K. & Lim, E. S. SV40 Large T Antigen Is Not Responsible for the Loss of STING in 293T Cells but Can Inhibit cGAS-STING Interferon Induction. Viruses 12, 137 (2020).

46. Remer, K. A., Brcic, M., Sauter, K.-S. & Jungi, T. W. Human monocytoid cells as a model to study Toll-like receptor-mediated activation. Journal of Immunological Methods 313, 1–10 (2006).

47. Alvarez-Carbonell, D. et al. Toll-like receptor 3 activation selectively reverses HIV latency in microglial cells. Retrovirology 14, 9 (2017).

48. Fenizia, C. et al. Human T-Cell Leukemia/Lymphoma Virus Type 1 p30, but Not p12/p8, Counteracts Toll-Like Receptor 3 (TLR3) and TLR4 Signaling in Human Monocytes and Dendritic Cells. Journal of Virology 88, 393–402 (2014).

49. Murshid, A., Borges, T. J., Lang, B. J. & Calderwood, S. K. The Scavenger Receptor SREC-I Cooperates with Toll-Like Receptors to Trigger Inflammatory Innate Immune Responses. Front. Immunol. 7, (2016).

50. Song, J., Guan, M., Zhao, Z. & Zhang, J. Type I Interferons Function as Autocrine and Paracrine Factors to Induce Autotaxin in Response to TLR Activation. PLOS ONE 10, e0136629 (2015).

51. Li, Z. et al. Lipofectamine 2000/siRNA complexes cause endoplasmic reticulum unfolded protein response in human endothelial cells. J Cell Physiol 234, 21166– 21181 (2019).

52. Simmons, D. P. et al. Type I IFN Drives a Distinctive Dendritic Cell Maturation Phenotype That Allows Continued Class II MHC Synthesis and Antigen Processing. J Immunol 188, 3116–3126 (2012).

53. Montoya, M. et al. Type I interferons produced by dendritic cells promote their phenotypic and functional activation. Blood 99, 3263–3271 (2002).

54. Rossi-Gendron, C. et al. Isothermal self-assembly of multicomponent and evolutive DNA nanostructures. Nat. Nanotechnol. 18, 1311–1318 (2023).

55. Van Tendeloo, V. F. I. et al. Highly efficient gene delivery by mRNA electroporation in human hematopoietic cells: superiority to lipofection and passive pulsing of mRNA and to electroporation of plasmid cDNA for tumor antigen loading of dendritic cells. Blood 98, 49–56 (2001).

56. Schaft, N. et al. Generation of an Optimized Polyvalent Monocyte-Derived Dendritic Cell Vaccine by Transfecting Defined RNAs after Rather Than before Maturation. J Immunol 174, 3087–3097 (2005).

57. Michiels, A. et al. Electroporation of immature and mature dendritic cells: implications for dendritic cell-based vaccines. Gene Ther 12, 772–782 (2005).

58. Met, Ö., Eriksen, J. & Svane, I. M. Studies on mRNA Electroporation of Immature and Mature Dendritic Cells: Effects on their Immunogenic Potential. Mol Biotechnol 40, 151–160 (2008).

59. Wilgenhof, S. et al. Therapeutic vaccination with an autologous mRNA electroporated dendritic cell vaccine in patients with advanced melanoma. J Immunother 34, 448–456 (2011).

60. Keersmaecker, B. D. et al. TriMix and tumor antigen mRNA electroporated dendritic cell vaccination plus ipilimumab: link between T-cell activation and clinical responses in advanced melanoma. J Immunother Cancer 8, (2020).

61. Donlin, L. T., Jayatilleke, A., Giannopoulou, E. G., Kalliolias, G. D. & Ivashkiv, L. B. Modulation of TNF-Induced Macrophage Polarization by Synovial Fibroblasts. J Immunol 193, 2373–2383 (2014).

62. Tassiulas, I. et al. Apoptotic Cells Inhibit LPS-Induced Cytokine and Chemokine Production and IFN Responses in Macrophages. Human Immunology 68, 156–164 (2007).

63. Son, Y. et al. Prostaglandin E2 is a negative regulator on human plasmacytoid dendritic cells. Immunology 119, 36–42 (2006).

64. Coulombe, F. et al. Targeted Prostaglandin E2 Inhibition Enhances Antiviral Immunity through Induction of Type I Interferon and Apoptosis in Macrophages. Immunity 40, 554–568 (2014).

65. Ingels, J., et al. Neoantigen-targeted dendritic cell vaccination in lung cancer patients induces long-lived T cells exhibiting the full differentiation spectrum. CR Med 5, (2024).

66. Lin-Shiao, E. et al. CRISPR–Cas9-mediated nuclear transport and genomic integration of nanostructured genes in human primary cells. Nucleic Acids Research 50, 1256–1268 (2022).

67. Hu, M. et al. Lantern-shaped flexible RNA origami for Smad4 mRNA delivery and growth suppression of colorectal cancer. Nat Commun 14, 1307 (2023).

68. Kretzmann, J. A. et al. Gene-encoding DNA origami for mammalian cell expression. Nat Commun 14, 1017 (2023).

69. Seitz, I. et al. Folding of mRNA-DNA Origami for Controlled Translation and Viral Vector Packaging. Advanced Materials 37, 2417642 (2025).

70. Wu, X. et al. An RNA/DNA hybrid origami-based nanoplatform for efficient gene therapy. Nanoscale 13, 12848–12853 (2021).

71. Anastassacos, F. M., Zhao, Z., Zeng, Y. & Shih, W. M. Glutaraldehyde Cross-Linking of Oligolysines Coating DNA Origami Greatly Reduces Susceptibility to Nuclease Degradation. J. Am. Chem. Soc. 142, 3311–3315 (2020).

72. Xia, T. et al. Thermodynamic Parameters for an Expanded Nearest-Neighbor Model for Formation of RNA Duplexes with Watson−Crick Base Pairs. Biochemistry 37, 14719–14735 (1998).

73. Mathews, D. H., Sabina, J., Zuker, M. & Turner, D. H. Expanded sequence dependence of thermodynamic parameters improves prediction of RNA secondary structure1. Journal of Molecular Biology 288, 911–940 (1999).

74. Li, Y. et al. Activation of RNase L is dependent on OAS3 expression during infection with diverse human viruses. Proceedings of the National Academy of Sciences 113, 2241–2246 (2016).

75. Li, Y. et al. Ribonuclease L mediates the cell-lethal phenotype of double-stranded RNA editing enzyme ADAR1 deficiency in a human cell line. eLife 6, e25687 (2017).

76. Parney, I. F. et al. Novel strategy for manufacturing autologous dendritic cell/allogeneic tumor lysate vaccines for glioblastoma. Neurooncol Adv 2, vdaa105 (2020).

