## Supplementary Materials for "Engineered mRNA nanostructures expand the design space of mRNA therapeutics through programmable protein expression and immune stimulation"

#### Supplementary Table S1. List of abbreviations.

| Abbreviation | Definition |
|---|---|
| **7-AAD** | 7-aminoactinomycin D (viability dye) |
| **A549** | human lung carcinoma cell line |
| **AFM** | atomic force microscopy |
| **ANOVA** | analysis of variance |
| **ATF3** | activating transcription factor 3 |
| **ATF4** | activating transcription factor 4 |
| **CCL2** | C-C motif chemokine ligand 2 (= MCP-1) |
| **CCL4** | C-C motif chemokine ligand 4 (= MIP-1β) |
| **CD8 / CD14 / CD40 / CD83 / CD86** | cluster of differentiation (surface markers 8, 14, 40, 83, 86) |
| **CDS** | coding sequence |
| **CHOP** | C/EBP-homologous protein (DDIT3) |
| **CXCL8** | C-X-C motif chemokine ligand 8 (= IL-8) |
| **CXCL10** | C-X-C motif chemokine ligand 10 (= IP-10) |
| **dsRBD** | double-stranded RNA-binding domain |
| **EDTA** | Ethylen ediaminetetraacetic acid |
| **EE** | encapsulation efficiency |
| **EGFP** | enhanced green fluorescent protein |
| **eIF2α** | eukaryotic translation initiation factor 2α |
| **ELISA** | enzyme-linked immunosorbent assay |
| **EMSA** | electrophoretic mobility shift assay |
| **EP** | electroporation |
| **EP2 / EP4** | prostaglandin E2 receptor 2 / receptor 4 |
| **ER** | endoplasmic reticulum |
| **FITC** | fluorescein isothiocyanate |
| **GADD34** | growth arrest and DNA-damage-inducible protein 34 (PPP1R15A) |
| **GFP** | green fluorescent protein |
| **GM-CSF** | granulocyte-macrophage colony-stimulating factor |
| **GMean** | geometric mean |
| **HEK293T** | human embryonic kidney 293T cell line |
| **HLA-DR** | human leukocyte antigen – DR isotype (MHC class II) |
| **HRP** | horseradish peroxidase |
| **IFN** | interferon |
| **IFN-β** | interferon beta |

| Abbreviation | Definition |
|---|---|
| **IFN-γ** | interferon gamma |
| **IFN-I** | type I interferon |
| **IFNAR** | type I interferon (α/β) receptor |
| **IgG** | immunoglobulin G |
| **IP-10** | interferon-γ–inducible protein 10 (= CXCL10) |
| **IRF3** | interferon regulatory factor 3 |
| **ISR** | integrated stress response |
| **ISRE** | interferon-stimulated response element |
| **IVT** | in vitro transcription |
| **JAK-STAT** | Janus kinase / signal transducer and activator of transcription |
| **K/O, K/D** | knockout, knockdown |
| **LNP** | lipid nanoparticle |
| **m5C** | 5-methylcytosine |
| **MCP-1** | monocyte chemoattractant protein-1 (= CCL2) |
| **MDA5** | melanoma differentiation-associated protein 5 (IFIH1) |
| **MFI** | mean fluorescence intensity |
| **MIP-1α** | macrophage inflammatory protein-1α (= CCL3) |
| **MIP-1β** | macrophage inflammatory protein-1β (= CCL4) |
| **moDC (iDC / mDC)** | monocyte-derived dendritic cell (immature / mature) |
| **mRNA-OG (OG)** | mRNA origami; OG-F = Folded, OG-U = Unfolded |
| **NF-κB** | nuclear factor κB |
| **OAT** | oxDNA Analysis Tools |
| **oxDNA / oxRNA** | coarse-grained DNA / RNA simulation models |
| **P:M** | polysome-to-monosome (ratio) |
| **PABP** | poly(A)-binding protein |
| **PB450** | Pacific Blue 450 (fluorophore) |
| **PBMC** | peripheral blood mononuclear cell |
| **PE** | phycoerythrin |
| **PERK** | PKR-like endoplasmic reticulum kinase |
| **$PGE_2$** | prostaglandin E2 |
| **PKR** | protein kinase R (EIF2AK2) |
| **poly(I:C)** | polyinosinic:polycytidylic acid (synthetic dsRNA) |
| **PRR** | pattern-recognition receptor |
| **PVDF** | polyvinylidene difluoride |
| **PX** | paranemic crossover |
| **RHA** | RNA helicase A (DHX9) |
| **RIG-I** | retinoic acid-inducible gene I (DDX58) |
| **SDS-PAGE** | sodium dodecyl sulfate–polyacrylamide gel electrophoresis |
| **STING** | stimulator of interferon genes |
| **TLR** | Toll-like receptor (TLR3, TLR7, TLR8) |

| Abbreviation | Definition |
| --- | --- |
| **TNF-α** | tumor necrosis factor alpha |
| **TNF-β** | tumor necrosis factor beta (lymphotoxin-α) |
| **UTR** | untranslated region |
| **Ψ** | pseudouridine |
| **ΔΔCt** | comparative (delta-delta) cycle threshold |

### Supplementary Table S2. Glossary of immunological readouts, receptors, and terms referenced in this study.

| Term / Readout | Definition / Assay (if measured directly) |
|---|---|
| **Directly measured readouts in this study** | |
| **phospho-eIF2α (p-eIF2α)** | ISR-activation marker (eIF2α phosphorylated on Ser51). Western blot, HEK293T. |
| **GADD34** | IF2α phosphatase (PPP1R15A) that terminates the ISR; the most specific eIF2α-axis reporter. qRT-PCR, HEK293T. |
| **CHOP** | Pro-apoptotic ATF4-axis factor (DDIT3) with convergent inputs (PERK, NF-κB), not eIF2α alone. qRT-PCR, HEK293T. |
| **ATF3** | Stress-inducible transcription factor downstream of the ATF4/ISR axis. qRT-PCR, HEK293T. |
| **IFN-β** | Type I interferon; secreted product of PRR→IRF3 signaling. ELISA, THP-1 and HEK293T. |
| **IP-10 (CXCL10)** | Interferon-stimulated chemokine; functional reporter of the complete sensing→IFN-β→IFNAR→ISRE axis. Secretome (CodePlex). |
| **IL-6** | NF-κB-dependent, broadly pro-inflammatory cytokine. Secretome (CodePlex). |
| **MIP-1β (CCL4) · MCP-1 (CCL2) · IL-8 (CXCL8) · MIP-1α (CCL3)** | Chemokines recruiting monocytes, granulocytes, and T cells. Secretome (CodePlex). |
| **IL-10** | Anti-inflammatory / tolerogenic cytokine. Secretome (CodePlex). |
| **T/NK effector analytes (IFN-γ, granzyme B, perforin, IL-2, sCD137, TNF-β)** | Adaptive/effector outputs on the Adaptive-Immune panel. Secretome (CodePlex). |
| **CD83** | Dendritic-cell maturation marker. Flow cytometry. |
| **CD86 (with CD40)** | Co-stimulatory molecule required for productive T-cell priming. Flow cytometry. |
| **HLA-DR** | MHC class II antigen-presentation molecule, upregulated on maturation. Flow cytometry. |
| **Nucleic-acid–sensing receptors (pattern-recognition receptors, PRRs)** | |
| **PKR (EIF2AK2)** | Cytosolic dsRNA-activated kinase; on binding dsRNA it dimerizes and phosphorylates eIF2α, triggering the integrated stress response. |
| **RIG-I (DDX58)** | Cytosolic sensor of short dsRNA bearing an uncapped 5′-triphosphate on a base-paired blunt end; signals via MAVS→IRF3 to type I IFN. A 5′ Cap1 structure marks RNA as "self" and evades it. |
| **MDA5 (IFIH1)** | Cytosolic sensor of long, extended dsRNA; signals via MAVS→IRF3 to type I IFN. |
| **TLR3** | Endosomal dsRNA receptor; signals via TRIF to IRF3 and NF-κB. |
| **TLR7 / TLR8** | Endosomal single-stranded-RNA receptors (via MyD88); not dsRNA sensors. |
| **RHA / DHX9** | Cytosolic RNA helicase that unwinds structured RNA and can feed type I IFN signaling. |
| **PRR (general)** | Germline-encoded receptor that detects conserved foreign/danger motifs, here exogenous RNA. |
| **Signaling adaptors, transcription factors & stress pathway** | |

| Term / Readout | Definition / Assay (if measured directly) |
|---|---|
| **MAVS** | Mitochondrial adaptor relaying RIG-I and MDA5 signals onward to IRF3 and NF-κB. |
| **TRIF / MyD88** | Adaptors downstream of TLR3 (TRIF) and TLR7/8 (MyD88). |
| **IRF3 / IRF7** | Transcription factors that drive type I IFN genes once phosphorylated by TBK1/IKKε. |
| **NF-κB (p65 / RelA)** | Transcription factor for inflammatory genes, including IL-6. |
| **eIF2α** | Eukaryotic translation-initiation factor; its phosphorylation halts cap-dependent initiation (measured as p-eIF2α, see readouts). |
| **ISR (integrated stress response)** | eIF2α-phosphorylation program that globally lowers translation while favoring stress transcripts (ATF4→CHOP, GADD34). |
| **Type I interferon axis (signaling nodes)** | |
| **Type I IFN (IFN-α/β)** | Antiviral cytokines induced by PRR→IRF3 signaling; act back on cells through IFNAR (IFN-β measured by ELISA, see readouts). |
| **IFNAR** | Receptor for type I IFN; engaging it activates JAK-STAT and induces interferon-stimulated genes. |
| **JAK-STAT** | Cascade downstream of IFNAR that transcribes ISGs via ISRE promoter elements. |
| **ISRE / ISG** | Interferon-stimulated response element / interferon-stimulated gene. |
| **Dendritic-cell biology & maturation** | |
| **moDC / iDC / mDC** | Monocyte-derived dendritic cell; immature versus TNFα/PGE2-matured. |
| **CD14** | Monocyte surface marker used to isolate the DC precursors. |
| **PGE2 (prostaglandin E2)** | Lipid mediator in the maturation cocktail; suppresses IRF3-driven type I IFN via EP2/EP4 receptors. |
| **EP2 / EP4** | PGE2 receptors that raise cAMP and dampen IRF3-driven antiviral signaling. |
| **Nucleic-acid & control terms** | |
| **dsRNA / ssRNA** | Double- versus single-stranded RNA. |
| **Cap1 / 5′-triphosphate** | 5′ structures distinguishing “self” (Cap1: $m^7G$ + 2′-O-methyl on N1) from RIG-I-activating uncapped 5′-ppp RNA. |
| **poly(I:C)** | Synthetic double-stranded RNA; canonical PRR agonist. |
| **Thapsigargin** | ER-stress inducer. |
| **Electroporation (EP)** | Delivery introducing RNA directly to the cytosol, bypassing endosomes. |

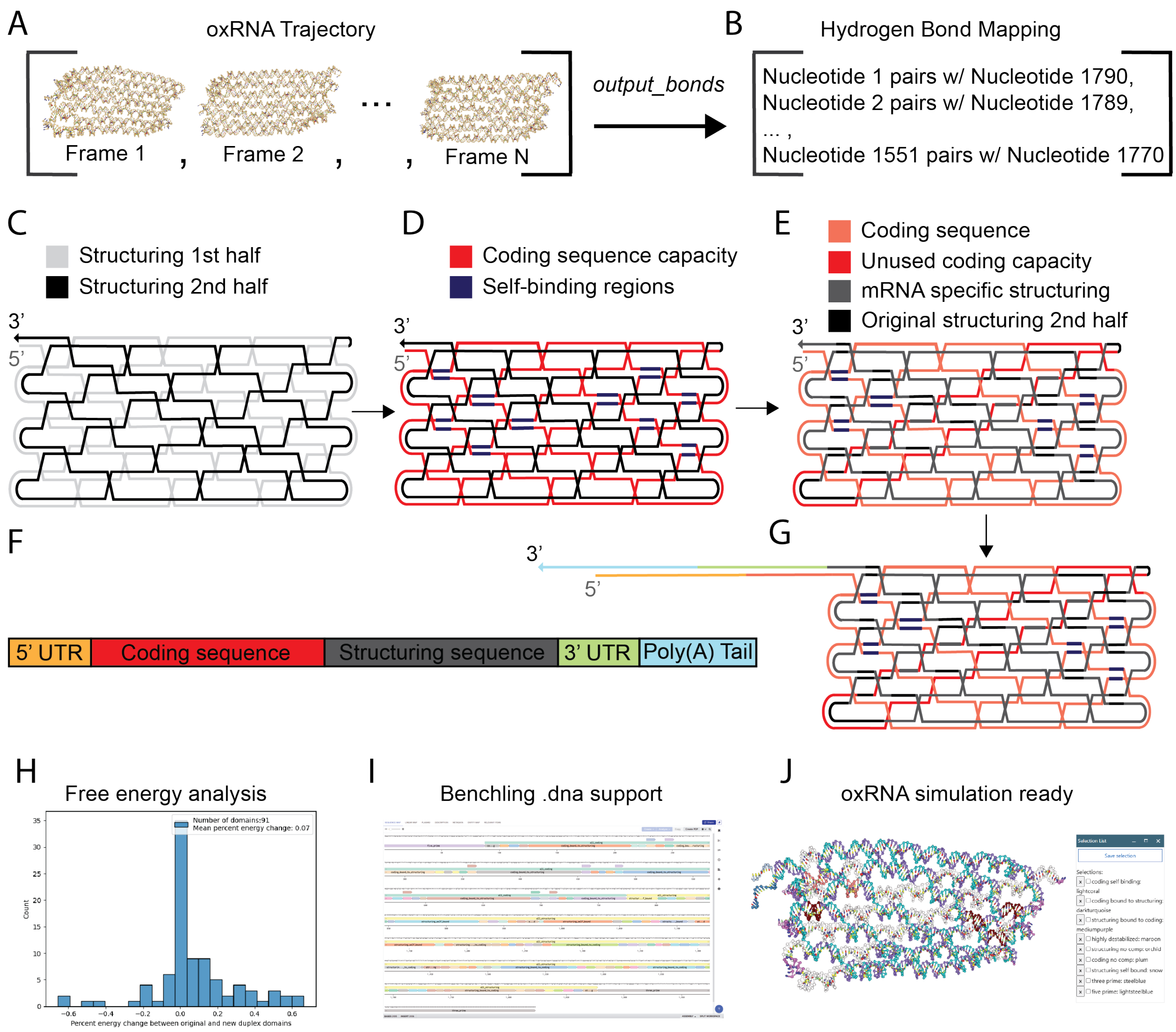


**Supplementary Figure S1. Messenger RNA origami (mRNA-OG) design algorithm, free energy validation, .dna file support, and simulation-ready export. (A-B)** The algorithm maps all hydrogen bond pairs from an optimally designed single-stranded RNA origami (ssRNA OG) using an oxRNA trajectory. **(C)** Strand routing schematic for a ssRNA OG, partitioned into two halves from 5' to 3'. **(D)** The first half defines the coding sequence capacity. Larger sequences require larger templates or an unstructured 5' tail. Structural "self-binding regions" are left unstructured. **(E)** The coding sequence is embedded into the capacity region (5' to 3'). Nucleotides in the second half designed to bind this sequence are mutated to complementary bases; unused capacity remains unchanged. **(F)** Linear sequence schematic correlating regions to specific colors. **(G)** 5' and 3' untranslated regions (UTRs) are added to the structure. **(H)** NUPACK calculates the free binding energy of all continuous duplexes before and after sequence embedding. The graph displays the percent free energy change, validating thermodynamic stability. **(I)** The mRNA-OG sequence is exported as an annotated .dna file for Benchling. **(J)** The embedded structure is exported as a simulation-ready, color-annotated oxView file.

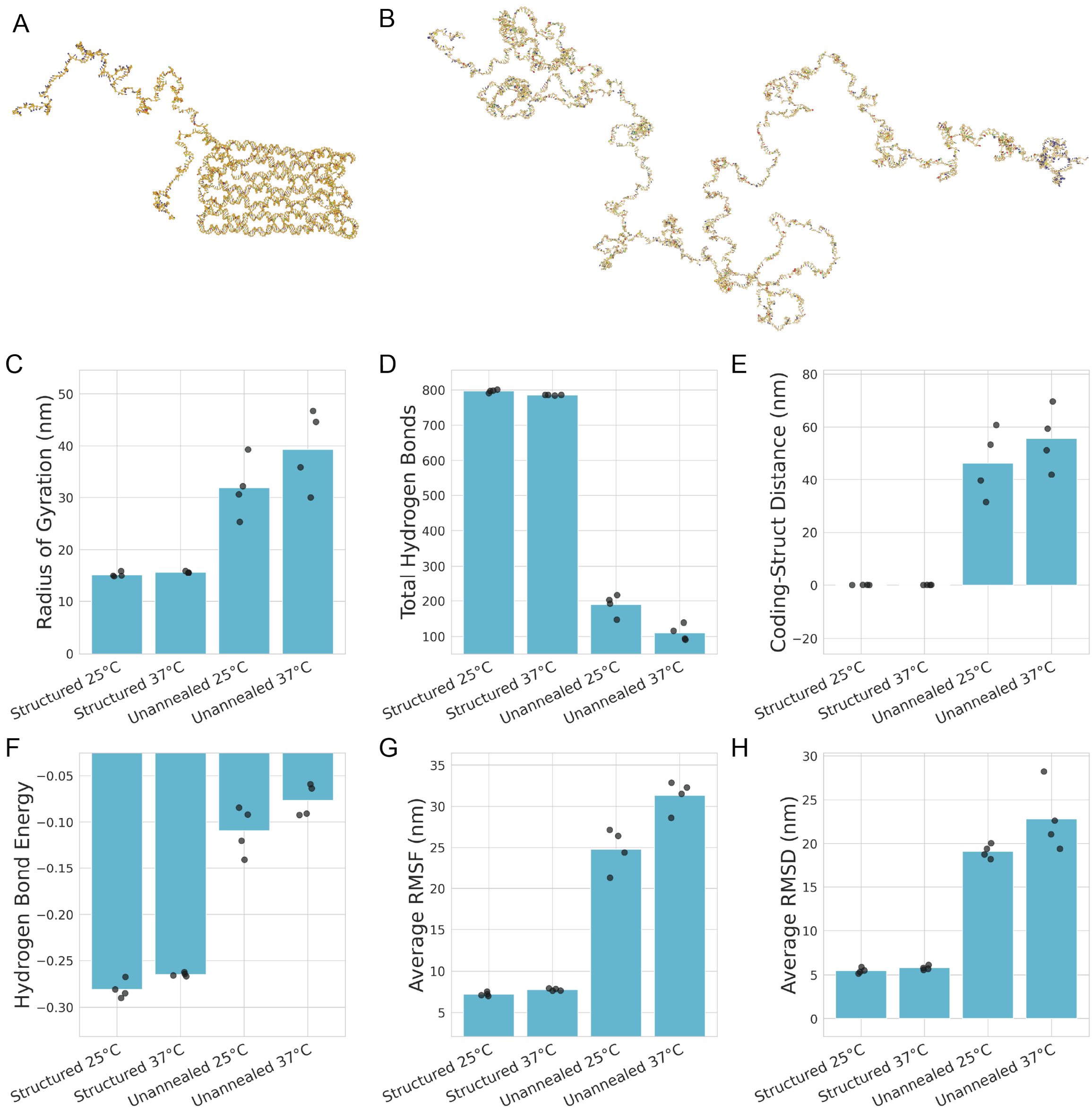


**Supplementary Figure S2. oxRNA simulations and measurements. (A)** compact, correctly folded structure of the mRNA-OG versus the **(B)** unannealed (Unfolded) mRNA-OG. We additionally measured **(C)** radius of gyration, **(D)** H-bond count, **(E)** mean distance between coding and structuring base pairs, **(F)** H-bond energy, **(G)** mean structure RMSF, and **(H)** mean structure RMSD over five independent trajectories.

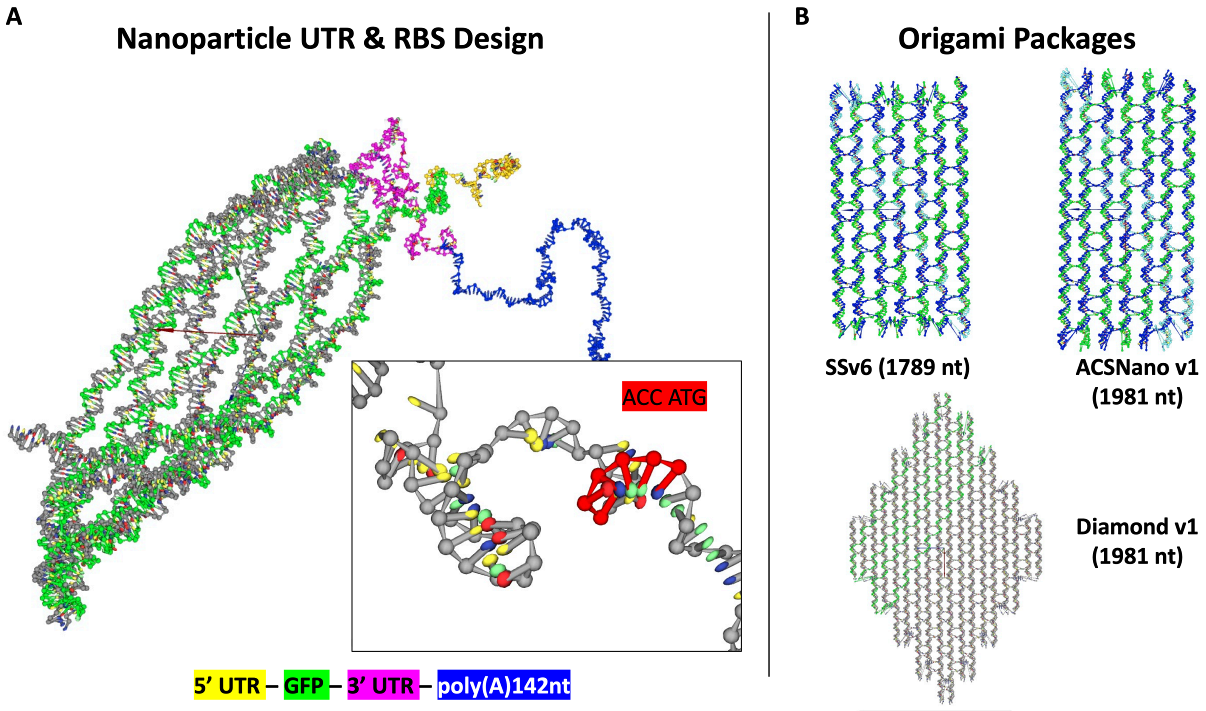


**Supplementary Figure S3. oxRNA render of relaxed nanoparticle and package options**. oxRNA render of mRNA origami packages including **(A)** single-stranded overhang regions with RBS (red) depicted and **(B)** alternative packaging strands (blue/grey strands) for identical EGFP coding sequences (green strand) demonstrating how nanoparticle CDS capacity is limited to about 50% of its total length.

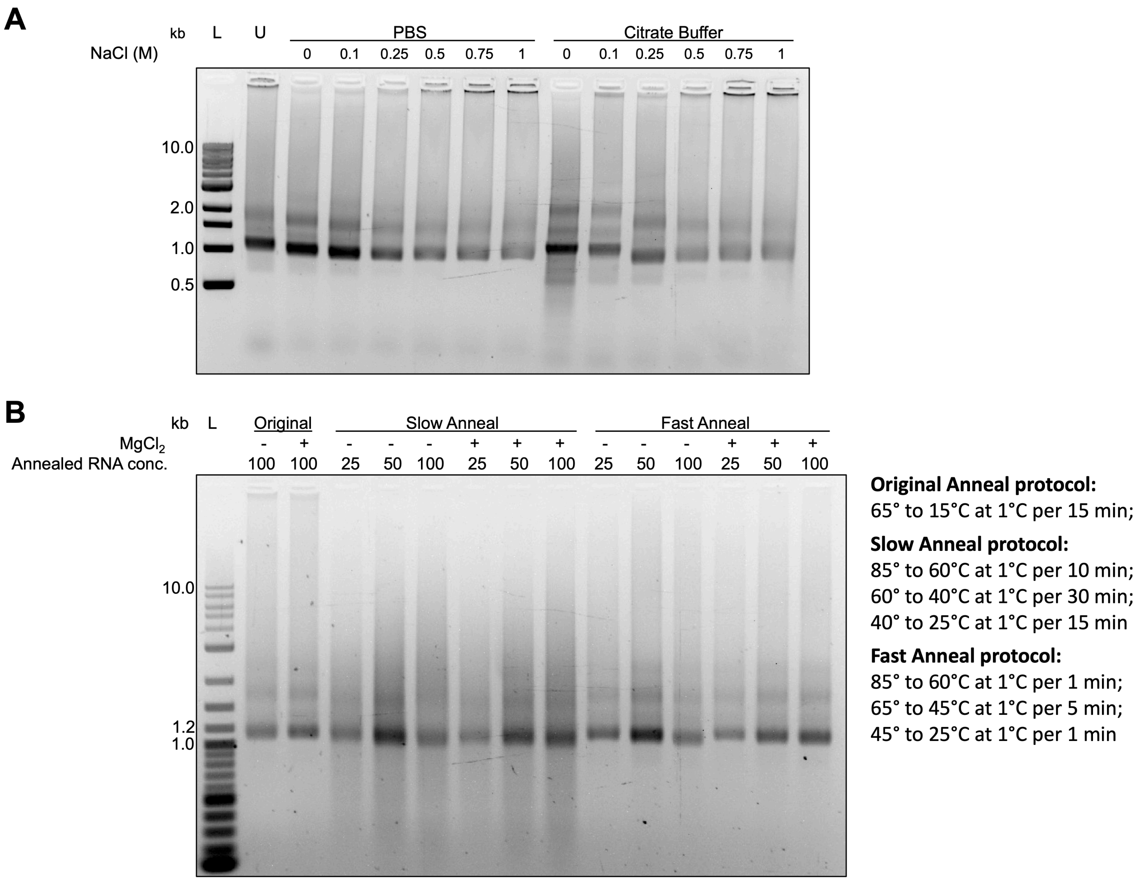


**Supplementary Figure S4. EGFP-OG RNA annealing characterization via agarose gel EMSA. (A)** EGFP-OG RNA titrated with various NaCl concentrations in 1x PBS and Citrate Buffer to find optimal annealing buffer determined by presence of agarose gel electrophoresis discrete band. Samples were annealed using Original annealing protocol (see B) with 100 ng/µL RNA. U is Unfolded mRNA-OG as control band. **(B)** EGFP-OG RNA thermally annealed across multiple RNA concentrations with three different protocols (original, slow, fast). Correctly folded product migrates as a discrete, compact band.

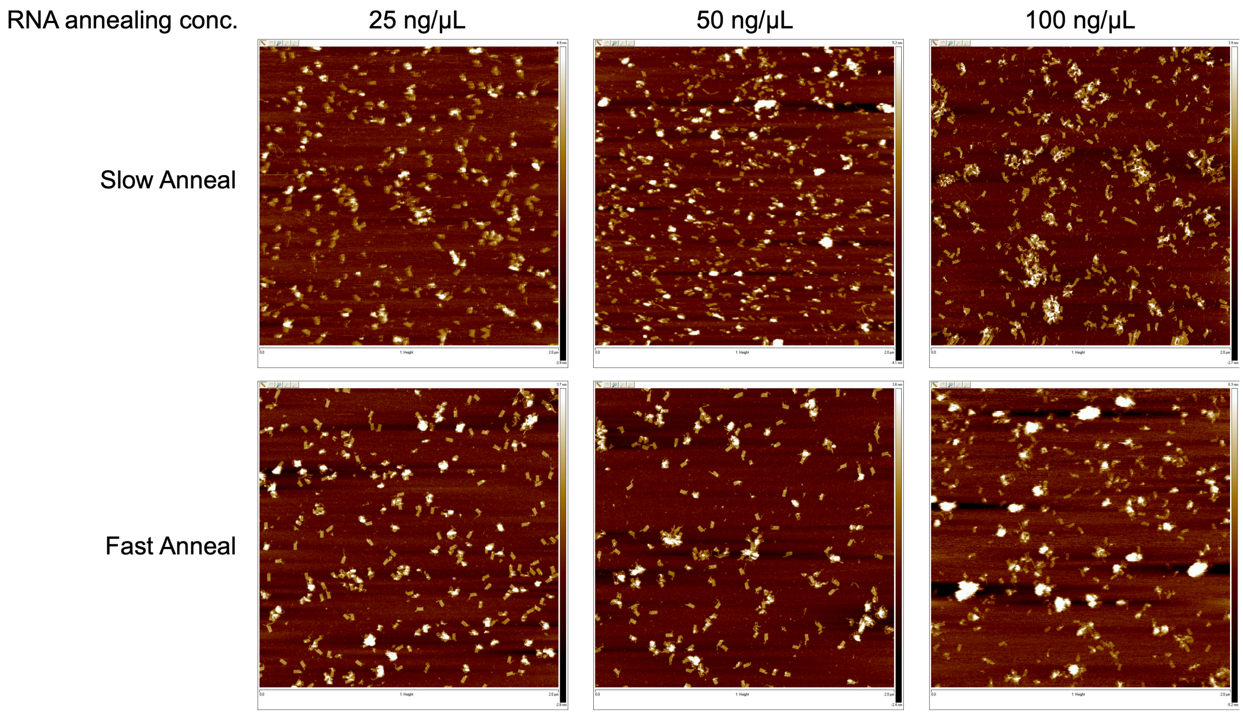


**Supplementary Figure S5. EGFP-OG RNA annealing concentration and thermal annealing characterization via atomic force microscopy (AFM).** EGFP-OG RNA thermally annealed across multiple concentrations with two protocols (slow, fast). Under optimal folding conditions, Folded OG adopts expected designed rectangular morphology with high monodispersity.

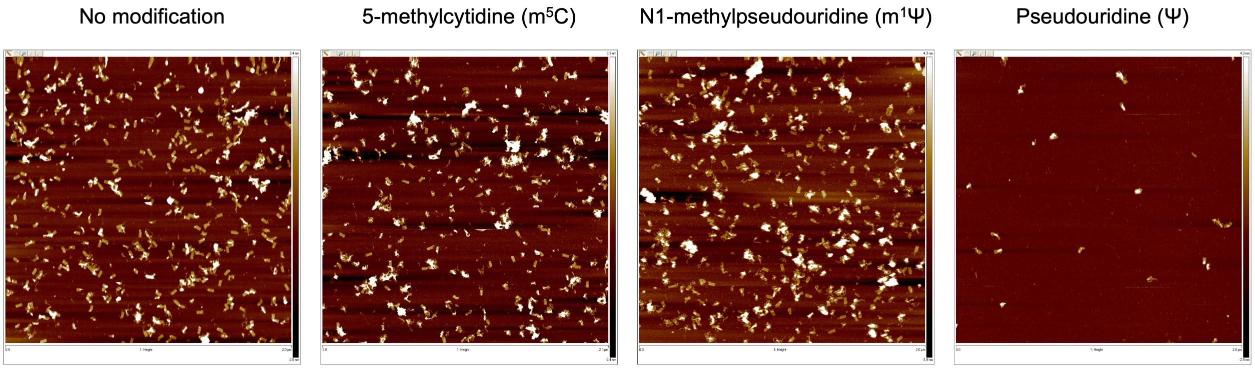


**Supplementary Figure S6. EGFP-OG modified nucleotide characterization.** AFM characterization of EGFP-OG RNA synthesized with 100% substitution of uridine or cytidine with 5-methylcytidine, N1-methylpseudouridine, or pseudouridine. AFM images also depict aggregation of nanoparticles due to the interaction between ssRNA UTRs of one particle and ssRNA loops present on 15 nm edge of another particle.

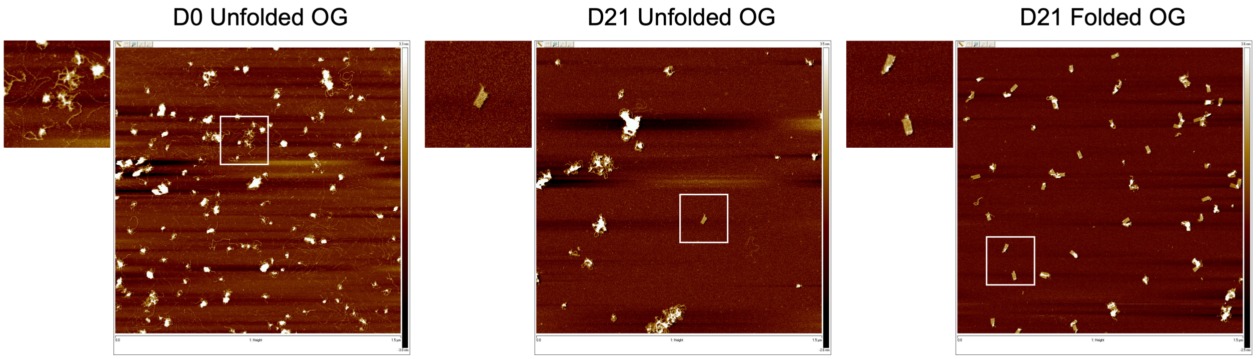


**Supplementary Figure S7. Spontaneous folding of Unfolded OG.** AFM characterization of EGFP-OG RNA at day 0 and day 21 of cold storage longevity study. At day 0, Unfolded mRNA-OG resembles a knotted string ball. By day 21 of 4°C storage, some Unfolded mRNA-OG sequences spontaneously adopt a folded conformation even in the absence of thermal annealing.

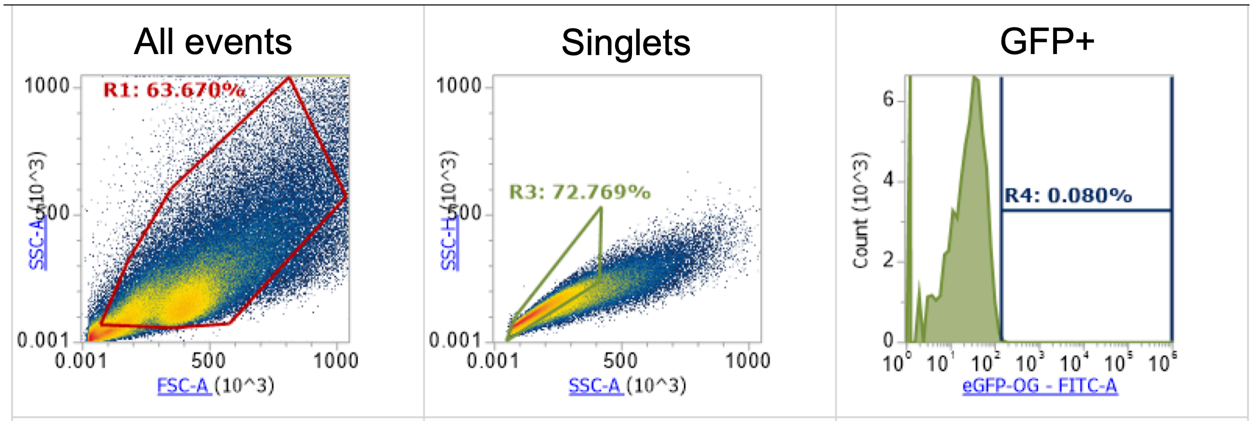


**Supplementary Figure S8. Flow cytometry gating strategy for cold storage longevity study in HEK293T cells of negative control at day 0.** Singlets gating strategy of HEK293T cells 24 hours post-transfection of negative control containing no EGFP mRNA. All samples in longevity study gated with same method.

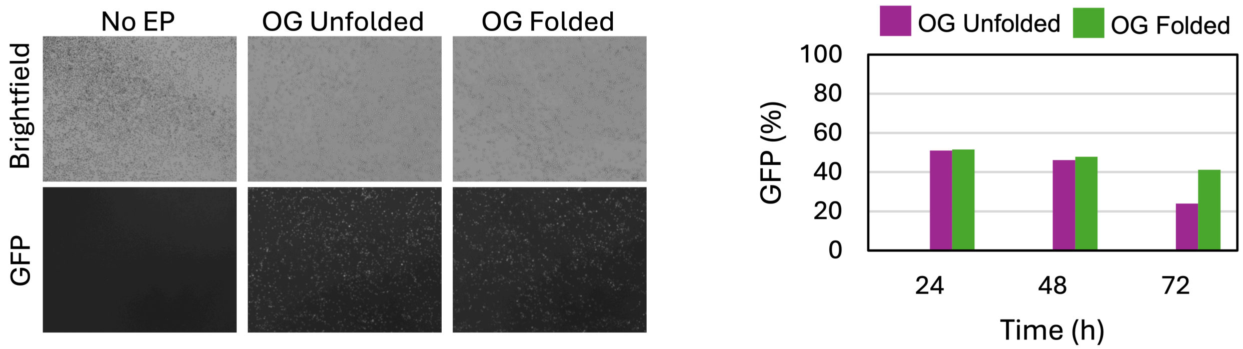


**Supplementary Figure S9. EGFP-OG expresses in primary DCs.** Brightfield and GFP micrographs depicting GFP expression in mature primary human dendritic cells (n=1).

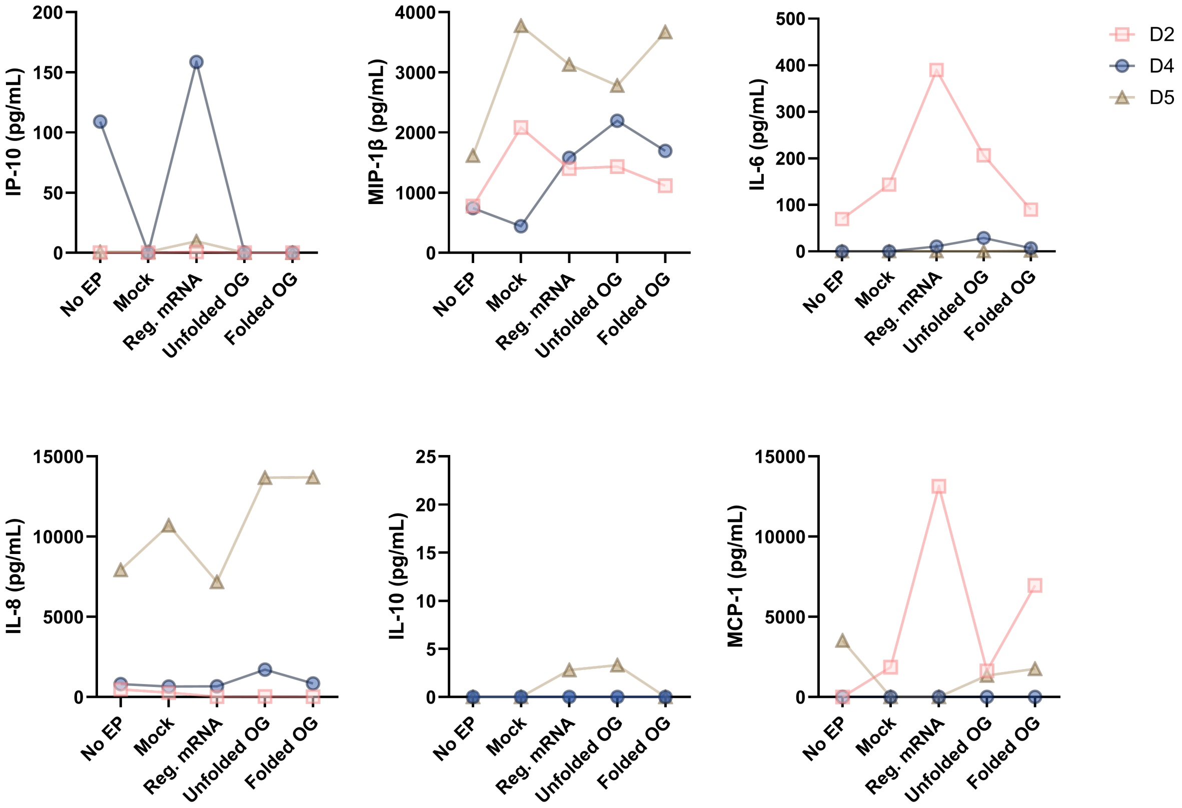


**Supplementary Figure S10. Selected analytes from multiplexed secretome results of mDC conditioned supernatants.** Concentrations of IP-10/CXCL10, MIP-1β/CCL4, IL-6, IL-8/CXCL8, IL-10, and MCP-1/CCL2 in conditioned supernatants collected 48 hours post-electroporation from TNFα/$PGE_2$-matured monocyte-derived dendritic cells (mDC) from donors D2 (pink squares), D4 (blue circles), and D5 (gold triangles). Donor D3 is excluded from this panel due to a non-standard 24-hour collection timepoint. IL-8 and MCP-1 were dominated by donor-level variation (D5 and D2, respectively) and showed no consistent RNA-condition-specific pattern across donors. MIP-1β and IL-10 were similarly unremarkable across RNA conditions. All values in pg/mL. Individual donor values shown as connected lines. Data represent technical duplicates within each donor (n = 3 biological donors at 48h). No statistical comparisons are made. See Supplementary Table 5 for complete numeric values.

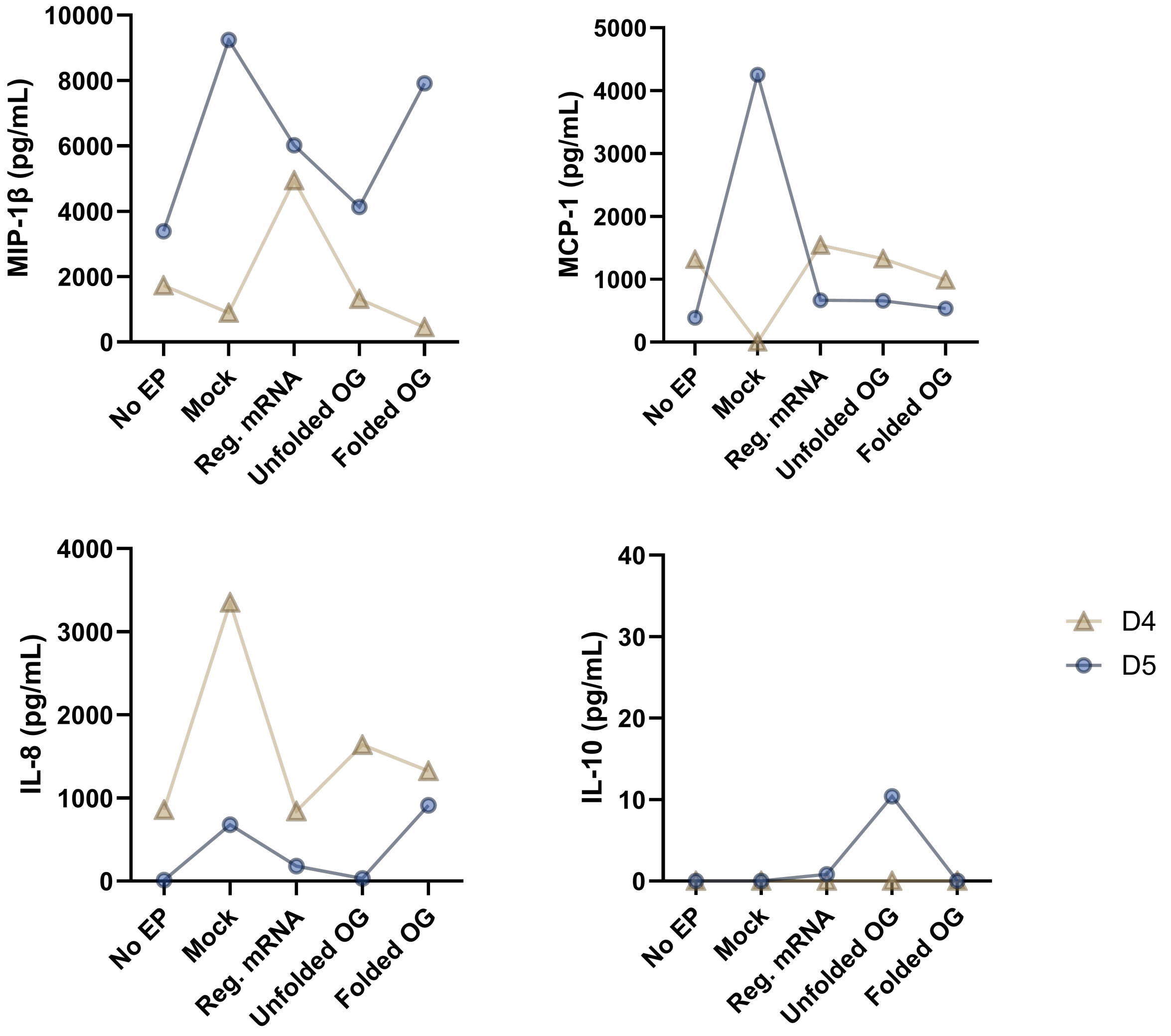


**Supplementary Figure S11. Selected analytes from multiplexed secretome results of iDC conditioned supernatants.** Concentrations of MIP-1β/CCL4, MCP-1/CCL2, IL-8/CXCL8, and IL-10 in conditioned supernatants collected 48 hours post-electroporation from immature monocyte-derived dendritic cells (iDC) from donors D4 (gold triangles) and D5 (blue circles). MIP-1β, MCP-1, and IL-8 showed no consistent RNA-condition-specific elevation above No EP or Mock controls across either donor. IL-10 was below the limit of detection in 9 of 10 samples; a single low-level signal (~10 pg/mL) was detected in the D5 Unfolded OG condition and is not interpreted as evidence of tolerogenic polarization given the absence of a cross-donor or cross-condition pattern. All values in pg/mL. Individual donor values shown as connected lines. Data represent technical duplicates within each donor (n = 2 biological donors). No statistical comparisons are made. See Supplementary Table 4 for complete numeric values.

| ID | Age (years) | Biological Sex (M/F/Other) | Race / Ethnicity | BMI ($kg/m^2$) | Smoking Status | Monocyte % (of PBMC) |
|---|---|---|---|---|---|---|
| D1 | 19 | M | Asian | 20.8 | Nonsmoker | 24.2 |
| D2 | 41 | F | White | 22.7 | Nonsmoker | 31.2 |
| D3 | 26 | M | White | 23.1 | Nonsmoker | 21.1 |
| D4 | 42 | M | White | 32.0 | Nonsmoker | 25.5 |
| D5 | N/A | F | N/A | 30.0 | Nonsmoker | 20.7 |

**Supplementary Table S3. Healthy donor demographics.** All donors provided by external vendors or Mayo Clinic Arizona (under IRB approval) via leukapheresis.

| Condition | Granzyme B | IFN-γ | IL-2 | IL-5 | IL-6 | IL-7 | IL-8 | IL-9 | IL-10 | IL-13 | IL-15 | IL-17A | IP-10 / CXCL10 | MCP-1 / CCL2 | MIP-1α / CCL3 | MIP-1β / CCL4 | Perforin | sCD137 | TNF-β |
|---|---|---|---|---|---|---|---|---|---|---|---|---|---|---|---|---|---|---|---|
| Donor 04 | | | | | | | | | | | | | | | | | | | |
| **No EP** | < LOD | < LOD | < LOD | < LOD | **398** | < LOD | 851 | < LOD | < LOD | < LOD | < LOD | < LOD | **1.17** | 1318 | < LOD | 1738 | < LOD | < LOD | < LOD |
| **Mock** | < LOD | < LOD | < LOD | < LOD | **347** | < LOD | 3388 | < LOD | < LOD | < LOD | < LOD | < LOD | < LOD | < LOD | 1.48 | 891 | < LOD | < LOD | < LOD |
| **Reg. mRNA** | 155 | < LOD | < LOD | < LOD | **186** | 204 | 832 | 2.51 | < LOD | 83.2 | < LOD | < LOD | **257** | 1549 | 3.89 | 4898 | < LOD | < LOD | < LOD |
| **Unfolded OG** | < LOD | < LOD | < LOD | < LOD | **234** | < LOD | 1622 | < LOD | < LOD | < LOD | < LOD | < LOD | **8318** | 1318 | < LOD | 1318 | < LOD | < LOD | < LOD |
| **Folded OG** | < LOD | < LOD | < LOD | < LOD | **309** | < LOD | 1318 | < LOD | < LOD | < LOD | < LOD | < LOD | **4898** | 1000 | < LOD | 457 | < LOD | < LOD | < LOD |
| Donor 05 | | | | | | | | | | | | | | | | | | | |
| **No EP** | < LOD | 214 | < LOD | < LOD | **145** | 245 | 11.5 | < LOD | < LOD | 23.4 | < LOD | < LOD | < LOD | 389 | 110 | 3388 | < LOD | < LOD | < LOD |
| **Mock** | < LOD | 72.4 | < LOD | < LOD | **389** | < LOD | 676 | < LOD | < LOD | 22.9 | < LOD | < LOD | < LOD | 4266 | 19.1 | 9333 | < LOD | 18.6 | 22.9 |
| **Reg. mRNA** | < LOD | < LOD | < LOD | < LOD | **100** | < LOD | 182 | < LOD | < LOD | 1.23 | < LOD | < LOD | **13.2** | 661 | 20.0 | 6026 | 67.6 | 110 | < LOD |
| **Unfolded OG** | < LOD | 457 | 11.5 | < LOD | **123** | 145 | 31.6 | 72.4 | 10.5 | < LOD | < LOD | < LOD | **6.31** | 661 | 135 | 4169 | 316 | 132 | < LOD |
| **Folded OG** | < LOD | < LOD | < LOD | < LOD | **437** | < LOD | 912 | < LOD | < LOD | < LOD | < LOD | < LOD | **23.4** | 537 | 6.03 | 7943 | < LOD | < LOD | < LOD |

**Supplementary Table S4. Complete multiplexed secretome of conditioned supernatants from immature monocyte-derived dendritic cells (iDC) electroporated with mRNA-OG constructs or controls.** Concentrations (pg/mL) of 19 analytes quantified from conditioned supernatants collected 48 hours post-electroporation from two healthy donors (D4 and D5). Supernatants were analyzed using a Bruker IsoLight system with a CodePlex Adaptive Immune — Human chip; concentrations were interpolated from on-chip standard curves and are reported as pg/mL back-calculated from $\log_{10}$ reporter fluorescence units (reported value = $10^{x}$). Values reported as '< LOD' were at or below the assay limit of detection (raw value = 0.00). Three analytes reflecting exogenous media components—GM-CSF (1,000 IU/mL), IL-4 (1,000 IU/mL), and TNF-α—are excluded from this table per the analysis described in the Methods. Three analytes—IL-5, IL-15, and IL-17A—were below the limit of detection in all 10 iDC samples across both donors and all conditions and are included for completeness. Data represent technical duplicates (duplicate CodePlex wells per condition per donor); iDC secretome comprises n = 2 biological donors.

| Condition | Granzyme B | IFN-γ | IL-2 | IL-5 | IL-6 | IL-7 | IL-8 | IL-9 | IL-10 | IL-13 | IL-15 | IL-17A | IP-10 / CXCL10 | MCP-1 / CCL2 | MIP-1α / CCL3 | MIP-1β / CCL4 | Perforin | sCD137 | TNF-β |
|---|---|---|---|---|---|---|---|---|---|---|---|---|---|---|---|---|---|---|---|
| **Donor D2** | | | | | | | | | | | | | | | | | | | |
| **No EP** | < LOD | < LOD | < LOD | < LOD | **69.2** | < LOD | 479 | < LOD | < LOD | 67.6 | < LOD | < LOD | < LOD | < LOD | 3.09 | 776 | < LOD | < LOD | < LOD |
| **Mock** | < LOD | < LOD | < LOD | < LOD | **145** | < LOD | 269 | < LOD | < LOD | < LOD | < LOD | 7.24 | < LOD | 1862 | 3.72 | 2089 | 30.2 | 2.95 | < LOD |
| **Reg. mRNA** | < LOD | < LOD | < LOD | 7.94 | **389** | < LOD | 17.4 | 13.2 | < LOD | 12.9 | < LOD | 1.78 | < LOD | 13183 | 63.1 | 1413 | 4.57 | < LOD | < LOD |
| **Unfolded OG** | < LOD | < LOD | < LOD | < LOD | **209** | < LOD | 33.9 | < LOD | < LOD | < LOD | < LOD | < LOD | < LOD | 1622 | 2.34 | 1445 | < LOD | < LOD | < LOD |
| **Folded OG** | < LOD | < LOD | 3.98 | < LOD | **89.1** | < LOD | 19.5 | < LOD | < LOD | 37.2 | < LOD | < LOD | < LOD | 6918 | 25.1 | 1122 | 29.5 | < LOD | < LOD |
| **Donor D3** | | | | | | | | | | | | | | | | | | | |
| **No EP** | < LOD | 447 | < LOD | < LOD | **275** | < LOD | < LOD | 562 | < LOD | < LOD | < LOD | < LOD | < LOD | 28840 | 15.8 | 871 | 138 | 132 | < LOD |
| **Mock** | < LOD | 525 | < LOD | < LOD | **219** | < LOD | < LOD | 871 | < LOD | < LOD | < LOD | < LOD | **10.0** | 30903 | 37.2 | 105 | 617 | 263 | < LOD |
| **Reg. mRNA** | < LOD | < LOD | < LOD | 49.0 | **64.6** | 14.5 | < LOD | < LOD | < LOD | < LOD | < LOD | < LOD | < LOD | 28184 | < LOD | 95.5 | 832 | < LOD | < LOD |
| **Unfolded OG** | < LOD | 148 | < LOD | < LOD | **52.5** | < LOD | < LOD | 513 | 5.25 | < LOD | < LOD | < LOD | < LOD | 30200 | 7.41 | < LOD | 562 | 186 | < LOD |
| **Folded OG** | < LOD | < LOD | < LOD | 1.95 | < LOD | < LOD | < LOD | < LOD | 3.24 | < LOD | < LOD | < LOD | < LOD | 29512 | 22.4 | < LOD | 692 | 91.2 | < LOD |
| **TriLink mRNA** | < LOD | < LOD | < LOD | < LOD | **17.0** | < LOD | < LOD | < LOD | 3.63 | < LOD | < LOD | < LOD | **5.01** | 28840 | < LOD | < LOD | 871 | 269 | < LOD |
| **Donor D4** | | | | | | | | | | | | | | | | | | | |
| **No EP** | < LOD | < LOD | < LOD | < LOD | < LOD | < LOD | 813 | 5.89 | < LOD | < LOD | < LOD | < LOD | **110** | < LOD | < LOD | 741 | < LOD | 112 | < LOD |
| **Mock** | < LOD | < LOD | < LOD | < LOD | < LOD | < LOD | 661 | < LOD | < LOD | < LOD | < LOD | < LOD | < LOD | < LOD | < LOD | 437 | < LOD | < LOD | < LOD |
| **Reg. mRNA** | 417 | < LOD | < LOD | < LOD | **10.7** | < LOD | 661 | < LOD | < LOD | < LOD | < LOD | < LOD | **158** | < LOD | 1.70 | 1585 | < LOD | 1.12 | < LOD |
| **Unfolded OG** | < LOD | < LOD | < LOD | < LOD | **28.8** | < LOD | 1698 | < LOD | < LOD | < LOD | < LOD | < LOD | < LOD | < LOD | 2.04 | 2188 | < LOD | < LOD | 105 |
| **Folded OG** | 437 | < LOD | < LOD | < LOD | **6.92** | 174 | 851 | 1.58 | < LOD | < LOD | < LOD | 3.09 | < LOD | < LOD | < LOD | 1698 | < LOD | < LOD | 12.3 |
| **Donor D5** | | | | | | | | | | | | | | | | | | | |
| **No EP** | < LOD | < LOD | < LOD | < LOD | < LOD | 41.7 | 7943 | < LOD | < LOD | < LOD | < LOD | < LOD | < LOD | 3548 | 4.17 | 1622 | < LOD | 89.1 | < LOD |
| **Mock** | < LOD | < LOD | < LOD | < LOD | < LOD | < LOD | 10715 | < LOD | < LOD | 14.1 | < LOD | < LOD | < LOD | < LOD | 10.5 | 3802 | < LOD | 112 | < LOD |
| **Reg. mRNA** | < LOD | < LOD | < LOD | < LOD | < LOD | < LOD | 7244 | < LOD | 2.82 | < LOD | < LOD | 2.45 | **9.55** | < LOD | 17.4 | 3162 | < LOD | 479 | < LOD |

| | | | | | | | | | | | | | | | | | | | |
|---|---|---|---|---|---|---|---|---|---|---|---|---|---|---|---|---|---|---|---|
| **Unfolded OG** | < LOD | 209 | < LOD | < LOD | < LOD | < LOD | 13804 | < LOD | 3.24 | < LOD | < LOD | < LOD | < LOD | 1349 | 28.8 | 2754 | < LOD | < LOD | < LOD |
| **Folded OG** | 269 | 135 | < LOD | < LOD | < LOD | < LOD | 13804 | < LOD | 1.38 | 105 | < LOD | < LOD | < LOD | 1778 | 33.9 | 3631 | 32.4 | < LOD | 79.4 |

**Supplementary Table S5. Complete multiplexed secretome of conditioned supernatants from TNFα/$PGE_2$-matured monocyte-derived dendritic cells (mDC) electroporated with mRNA-OG constructs or controls.** Concentrations (pg/mL) of 19 analytes quantified from conditioned supernatants collected 48 hours post-electroporation from four healthy donors (D2, D4, and D5) or 24 hours post-electroporation for donor 3 (non-standard timepoint; noted in-table). IL-15 was below the limit of detection in all 21 mDC samples across all donors and conditions and is included for completeness. IP-10/CXCL10 was at or below the assay floor across all RNA and mock conditions in mDC for all four donors at both available timepoints (D2, D3, D4, D5; 24h and 48h). Low-level IP-10 signals detected in No EP and Regular mRNA conditions in isolated donors (D4: No EP = 110 pg/mL, Regular mRNA = 158 pg/mL). IL-6 showed a directional hierarchy in mDC (Regular mRNA > Unfolded OG > Folded OG) in two donors across two independent timepoints (D2 at 48h; D3 at 24h), as described in the main text and Supplementary Figure 11. Data represent technical duplicates (duplicate CodePlex wells per condition per donor); mDC secretome comprises n = 4 biological donors (n = 3 at 48h). All values reported as described in Supplementary Table S4.

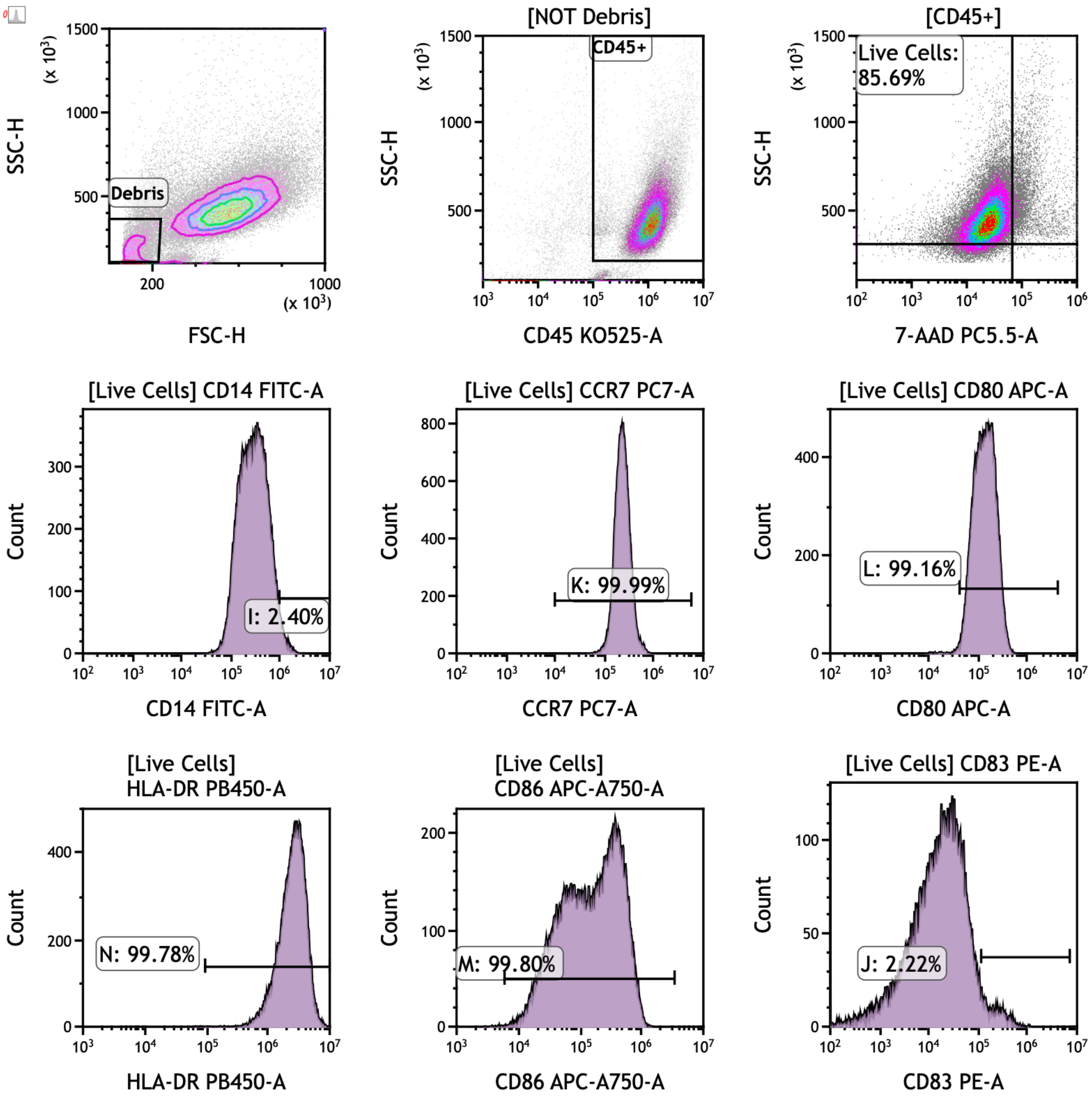


**Supplementary Figure S12. Flow cytometry gating strategy and phenotypic characterization of immature monocyte-derived dendritic cells (iDC) from Donor 2 at day 5 of culture.** Sequential gating hierarchy: scatter-based debris exclusion → CD45+ leukocyte selection → 7-AAD− live cell gate (85.69% viable). Surface marker expression on live cells: CD14 2.40% (confirming monocyte-to-DC differentiation), CD83 2.22% (confirming immature state), CCR7 99.99%, CD80 99.16%, CD86 99.80%, HLA-DR 99.78%.

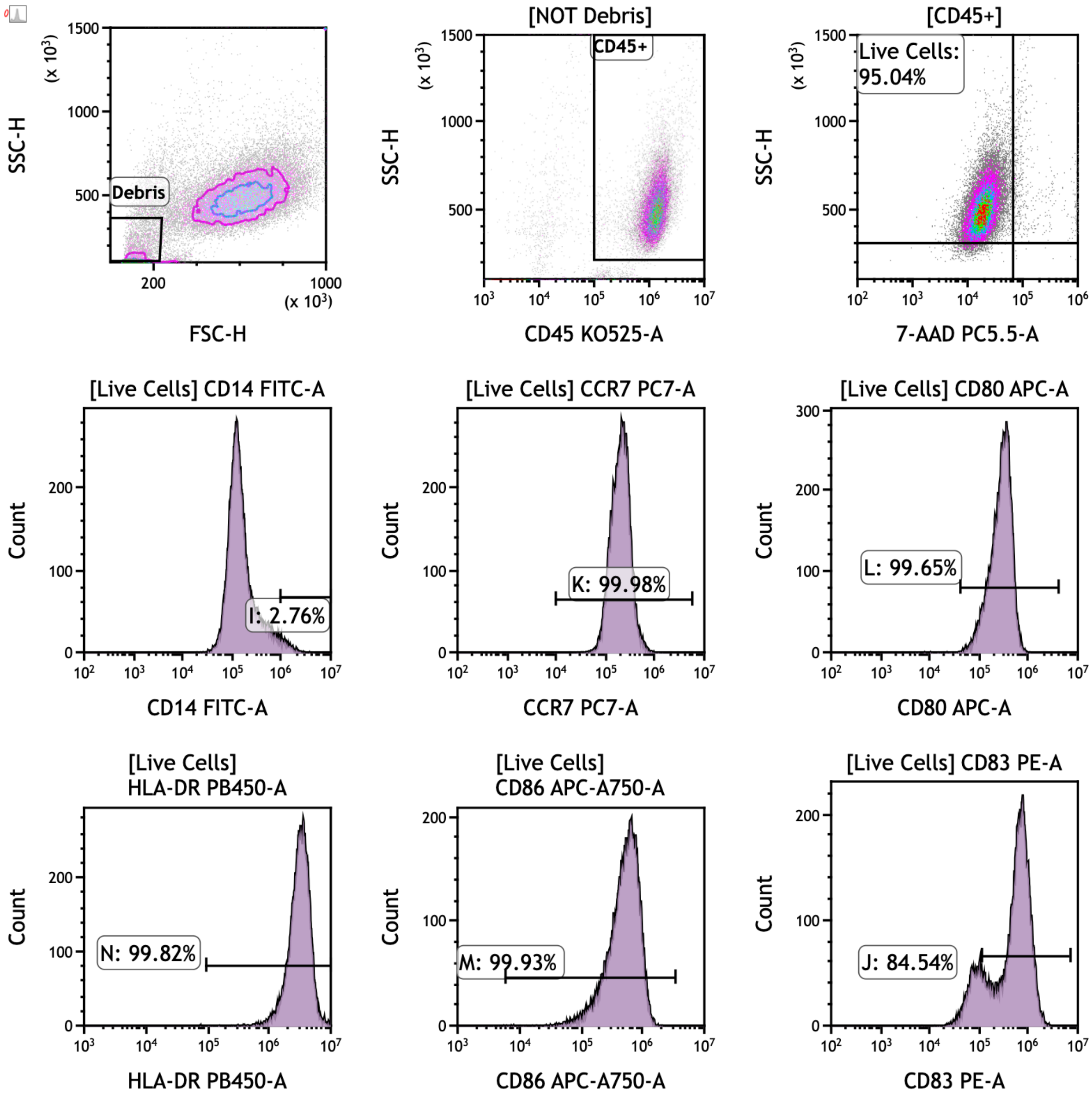


**Supplementary Figure S13. Flow cytometry gating strategy and phenotypic characterization of TNFα/$PGE_2$-matured monocyte-derived dendritic cells (mDC) from Donor 2 at day 7 of culture.** Sequential gating hierarchy: scatter-based debris exclusion → CD45$^+$ leukocyte selection → 7-AAD$^-$ live cell gate (95.04% viable). Surface marker expression on live cells: CD14 2.76% (confirming DC identity), CD83 84.54% (confirming successful maturation), CCR7 99.98%, CD80 99.65%, CD86 99.93%, HLA-DR 99.82%.

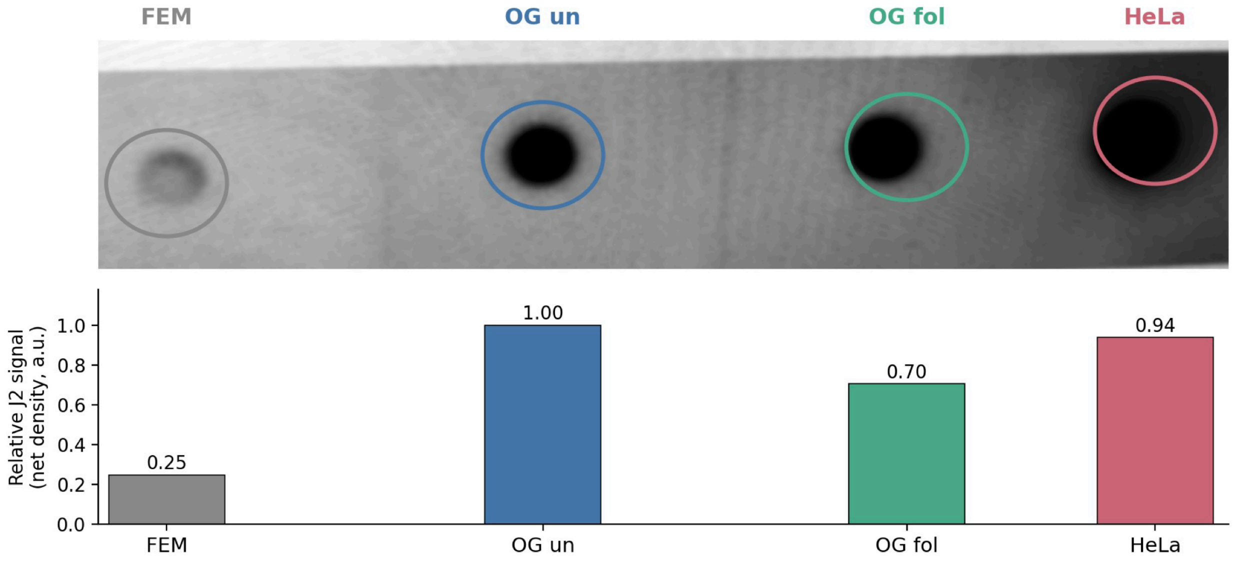


**Supplementary Figure S14. J2 (anti-dsRNA) immuno-dot blot.** J2 immuno-dot blot of FEM (linear RNA negative control), Unfolded mRNA-OG, Folded mRNA-OG, and HeLa RNA (positive control) qualitatively demonstrating recognition of J2 dsRNA binding motif in the folded nanostructure.

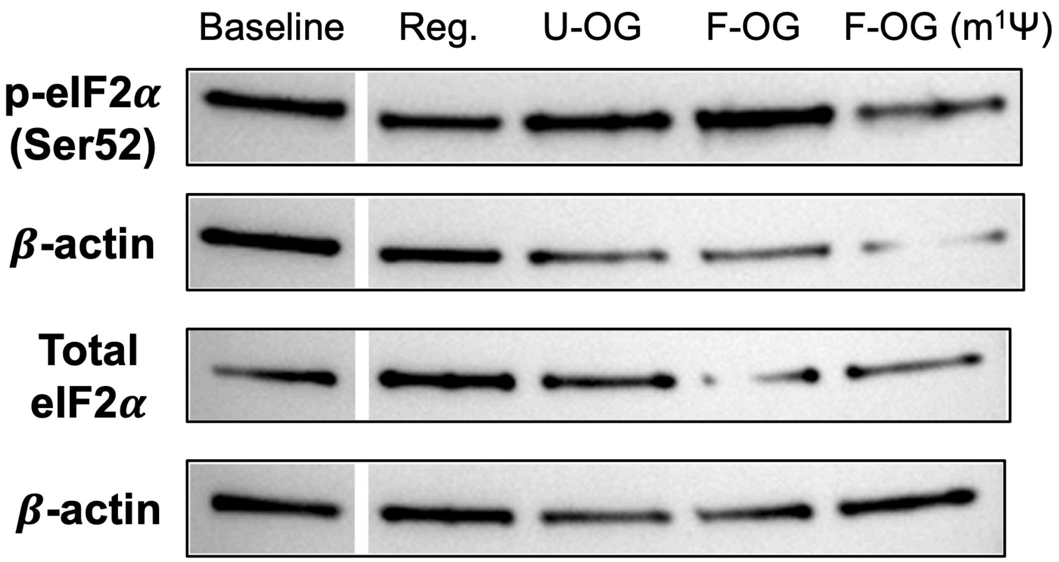


**Supplementary Figure S15. Western blot of phospho-eIF2α and total eIF2α with $\beta$-actin controls.** Western blot of phospho-eIF2α and total eIF2α with $\beta$-actin control in HEK293T cells 24 hours post-lipofection. Baseline is HEK293T without mRNA, Reg. is Regular mRNA, U-OG is Unfolded-OG, F-OG is Folded-OG, and F-OG ($m^1\Psi$) is Folded mRNA-OG synthesized with 100% substitution of uridine for $m^1\Psi$.

**Supplementary Table S6. EGFP mRNA-OG sequence.**

5' UTR – coding (EGFP) – structuring – 3' UTR – poly(A)x142

AGGAAATAAGAGAGAAAAGAAGAGTAAGAAGAAATATAAGAGCCACCATGGTGAGC
AAGGGCGAGGAGCTGTTCACCGGGGTGGTGCCCATCCTGGTCGAGCTGGACGGC
GACGTAAACGGCCACAAGTTCAGCGTGTCCGGCGAGGGCGAGGGCGATGCCACC
TACGGCAAGCTGACCCTGAAGTTCATCTGCACCACCGGCAAGCTGCCCGTGCCCT
GGCCCACCCTCGTGACCACCCTGACCTACGGCGTGCAGTGCTTCAGCCGCTACCC
CGACCACATGAAGCAGCACGACTTCTTCAAGTCCGCCATGCCCGAAGGCTACGTC
CAGGAGCGCACCATCTTCTTCAAGGACGACGGCAACTACAAGACCCGCGCCGAGG
TGAAGTTCGAGGGCGACACCCTGGTGAACCGCATCGAGCTGAAGGGCATCGACTT
CAAGGAGGACGGCAACATCCTGGGGCACAAGCTGGAGTACAACTACAACAGCCAC
AACGTCTATATCATGGCCGACAAGCAGAAGAACGGCATCAAGGTGAACTTCAAGAT
CCGCCACAACATCGAGGACGGCAGCGTGCAGCTCGCCGACCACTACCAGCAGAA
CACCCCCATCGGCGACGGCCCCGTGCTGCTGCCCGACAACCACTACCTGAGCAC
CCAGTCCGCCCTGAGCAAAGACCCCAACGAGAAGCGCGATCACATGGTCCTGCTG
GAGTTCGTGACCGCCGCCGGGATCACTCTCGGCATGGACGAGCTGTACAAGTAAT
AGTGAGAAGTTGCCATCGTAGTCGCACGACCTGGACACGCCGAACTTCCGCGGGT
CGGCTGACGTCGATGGGGTGCGTGCAAAAAAGACCTACGAAGCCAGAGTTCGTTC
CAGTGTGAAAGTGCACATCACGAGTTGTGCCAATGCACGTTGCATCGAGGGCTGA
AGCCGTCTTAATATAGACGGCACCTGAAGAGTGATTGATTCGTCTAGAAATAGACGA
ATCATGCTGATCTCAGGTGCTCACTTGATTAAGACGGCTGTTTATCTCGATGCTCGC
CCTCTTGGCACAATCGAACTTGTGCACTTCAGCACGGGAACGAACTCTGGCTTCGT
AGGTCAAAAAAGCACGCAAGCATGTAACGTCAGCCTAACGCTTGAAGTTCGCAGGT
GTGAGGTCGTGCATGTGGTCTGGCAACTCTGCTTGTTTACTTGTACAGCTCGTCCA
TGCCGGATCAGCACCCGGCGAAAAAAACGAACTGTGAGGTAACCATGTGGATACTC
TTCTCGTTGGCTTGCCGTCTCAGGGCTCACCAGGTGCTCAGGTAGTGGTTGTCGG
GCAGTCACACTGGGCCGTCGCTACATGCTGTGTTCTAAAAAAAGTGGTCGCACTAT
AGCACGCTGCCTCGTACAATGTTGTGAGAAGTCGTGAAGTTCACCTTGATGCCGTT
CTTTCTCACTACGGCCATGATCAAGTGATTGTGGCTTACCTCACTGTACTCAAAAAA
GTGCCCCAAGATTTGGCCGTCCTATGAACTTTCGATGCCCTTCAGCTCGATGCGGT
GGACTGGGGTGTCGCCCCTCGTGATCACCTCGGAAGCGTTATTGTAGTTGTATAGT
GCCTTGAAGAAAAAAGTGCGCTCTTGATCATAGCCTTCGGGCATGGCGGACTTGAG
CGGATCTTGCTGCTTCGACTACGAGGGGTAGCATAAACAGCACTGCACGAGAGTAT
CCAGGGTGGCAAATCTTGTGGGCCAAAAAACGGGCAGCTTGCCGGTGGTGCAGC
CTTGAAGCAGGGTCAGGGTCTTTGAGGTGGCAAACGTGCAGCCCTCGCCCACACC
TGTGAACTTGTGTACGAGTACGTCGCCTGATCAAGCGACCAGGATGGGGCGGCCG
CTTAATTAAGCTGCCTTCTGCGGGGCTTGCCTTCTGGCCATGCCCTTCTTCTCTCCC
TTGCACCTGTACCTCTTGGTCTTTGAATAAAGCCTGAGTAGGAAGAAAAAAAAAAAA
AAAAAAAAAAAAAAAAAAAAAAAAAAAAAAAAAAAAAAAAAAAAAAAAAAAAAAAAAAAAAAA

AAAAAAAAAAAAAAAAAAAAAAAAAAAAAAAAAAAAAAAAAAAAAAAAAAAAAAAAAA
AAAAAAAAAAAAAA

**SOURCE IMAGES (for Supplementary Figures):**

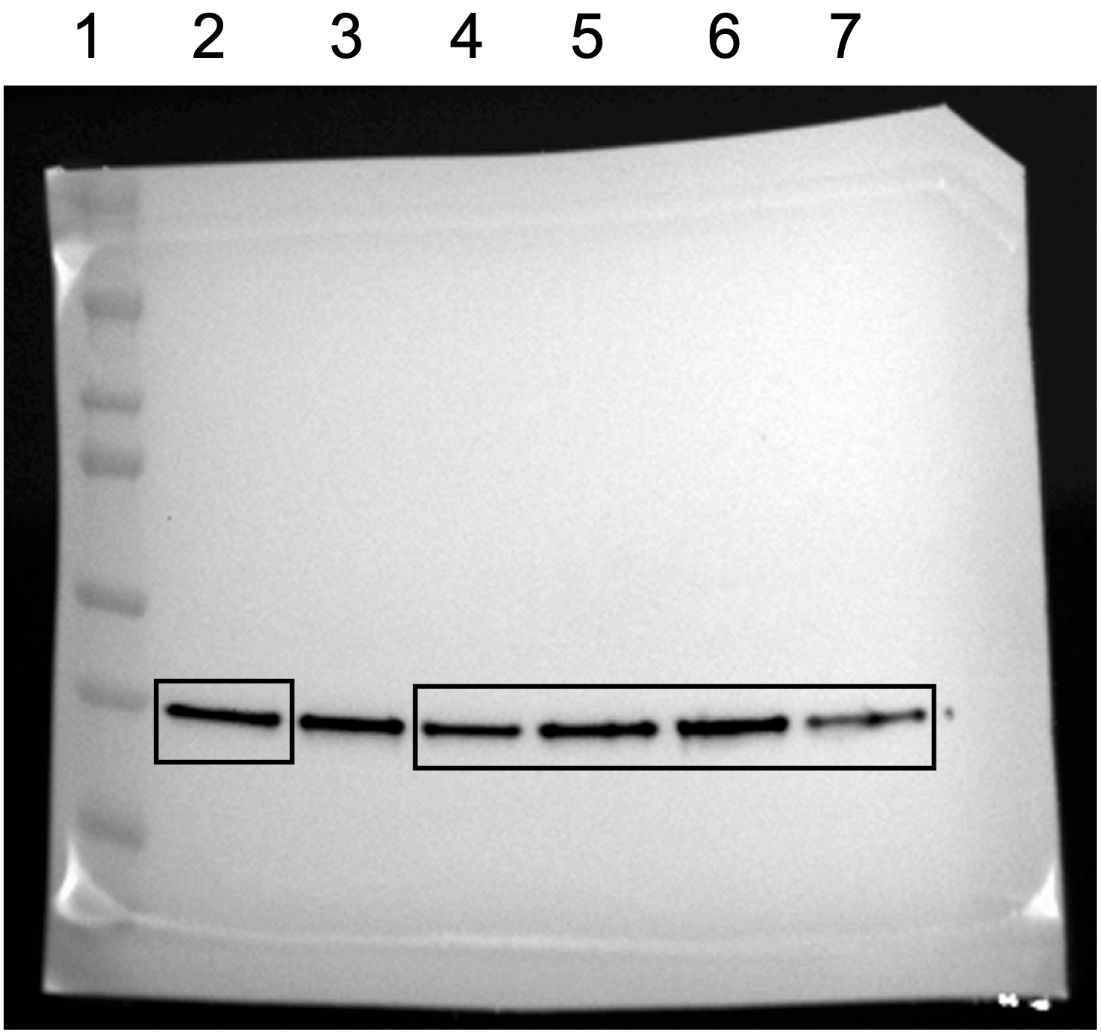


**Uncropped western blot for p-eIF2α in Supplementary Figure S15.** Western blot of phospho-eIF2α in HEK293T cells 24 hours post-lipofection. 1) Marker 2) HEK293T without mRNA 3) Thapsigargin positive control 4) Regular mRNA 5) Unfolded-OG 6) Folded-OG 7) Folded mRNA-OG synthesized with 100% substitution of uridine for $m^1\Psi$

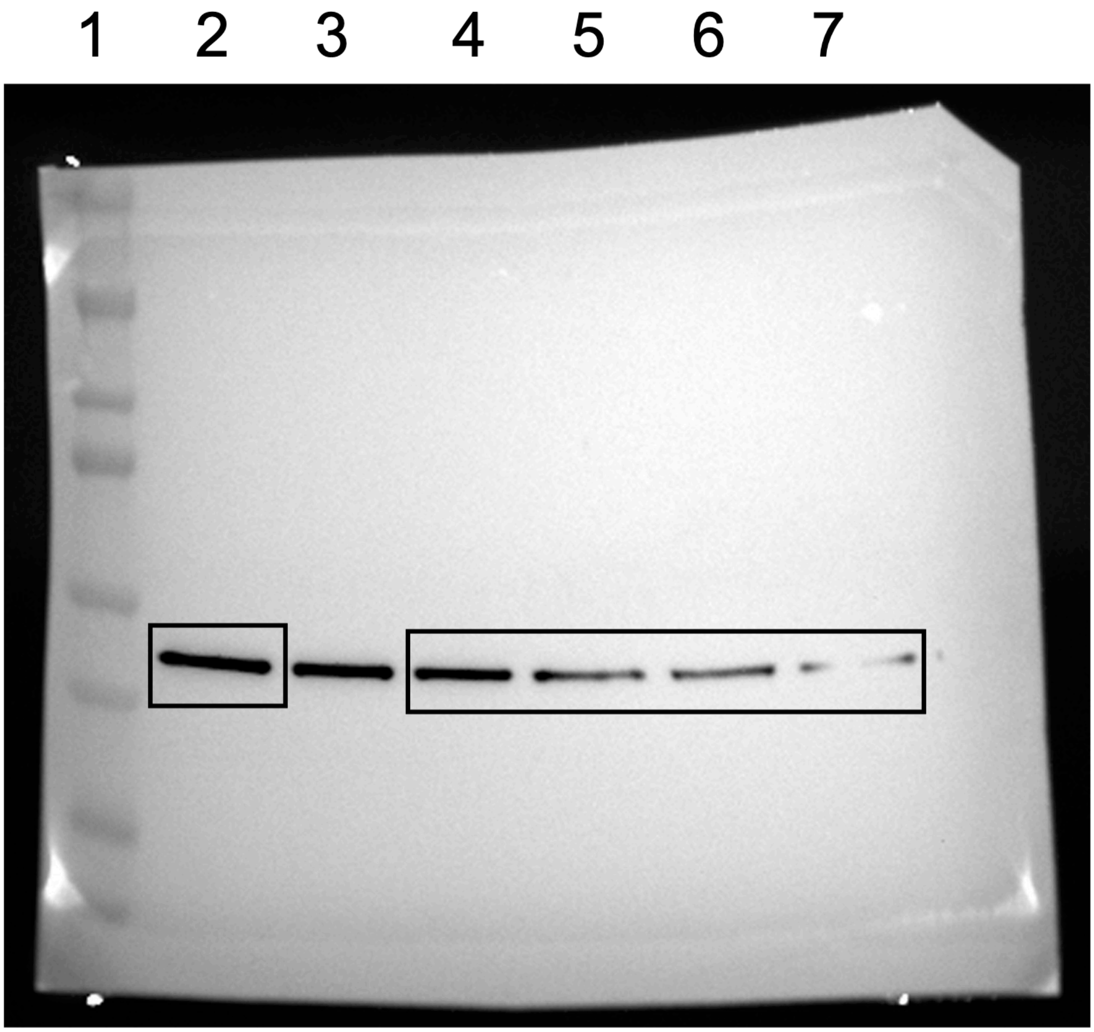


**Uncropped western blot for $\beta$-actin control for p-eIF2α in Supplementary Figure S15.** Western blot of $\beta$-actin for p-eIF2α in HEK293T cells 24 hours post-lipofection. 1) Marker 2) HEK293T without mRNA 3) Thapsigargin positive control 4) Regular mRNA 5) Unfolded-OG 6) Folded-OG 7) Folded mRNA-OG synthesized with 100% substitution of uridine for $m^1\Psi$

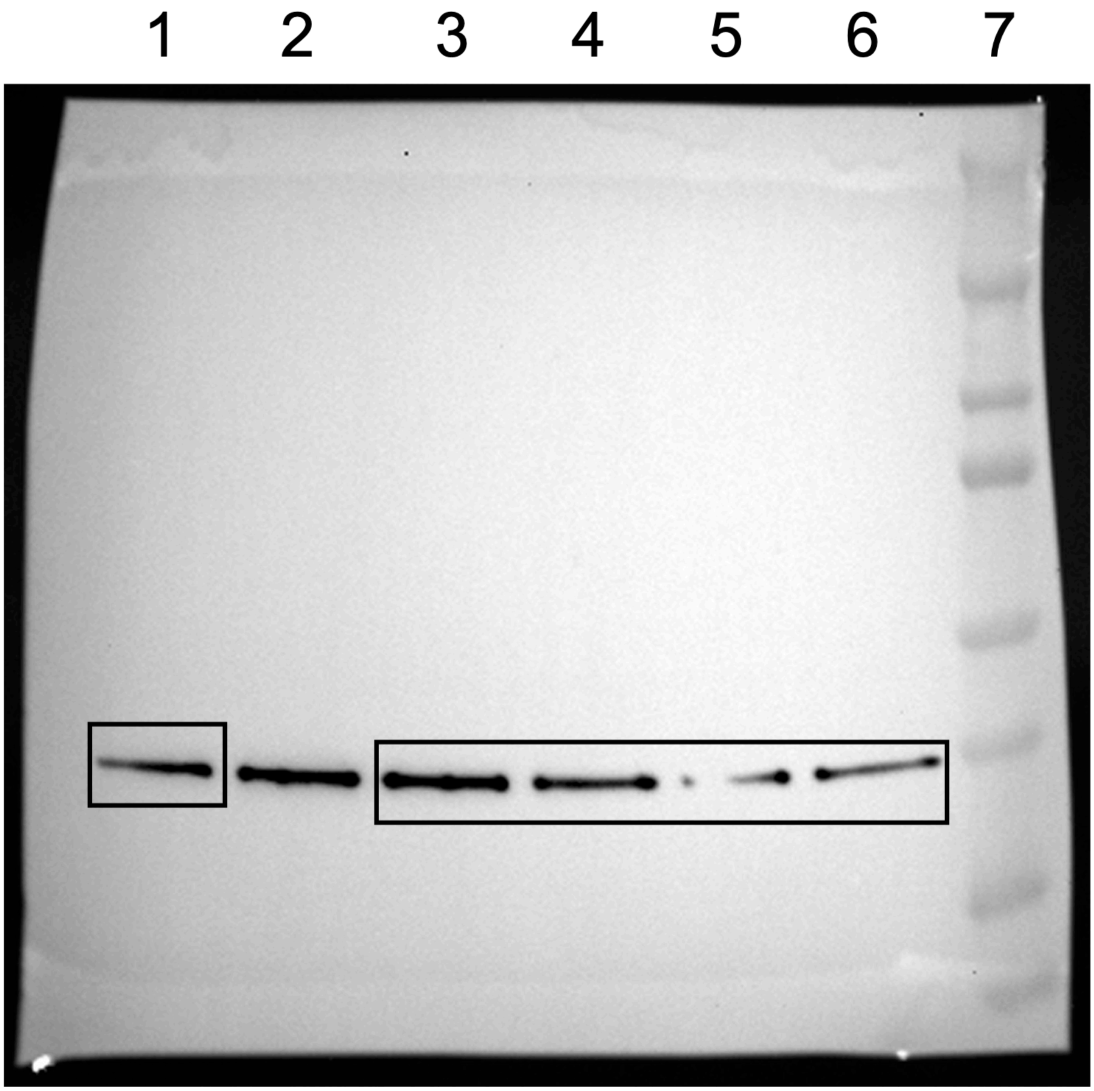


**Uncropped western blot for total eIF2α in Supplementary Figure S15.** Western blot of total eIF2α in HEK293T cells 24 hours post-lipofection. 1) HEK293T without mRNA 2) Thapsigargin positive control 3) Regular mRNA 4) Unfolded-OG 5) Folded-OG 6) Folded mRNA-OG synthesized with 100% substitution of uridine for $m^1\Psi$ 7) Marker

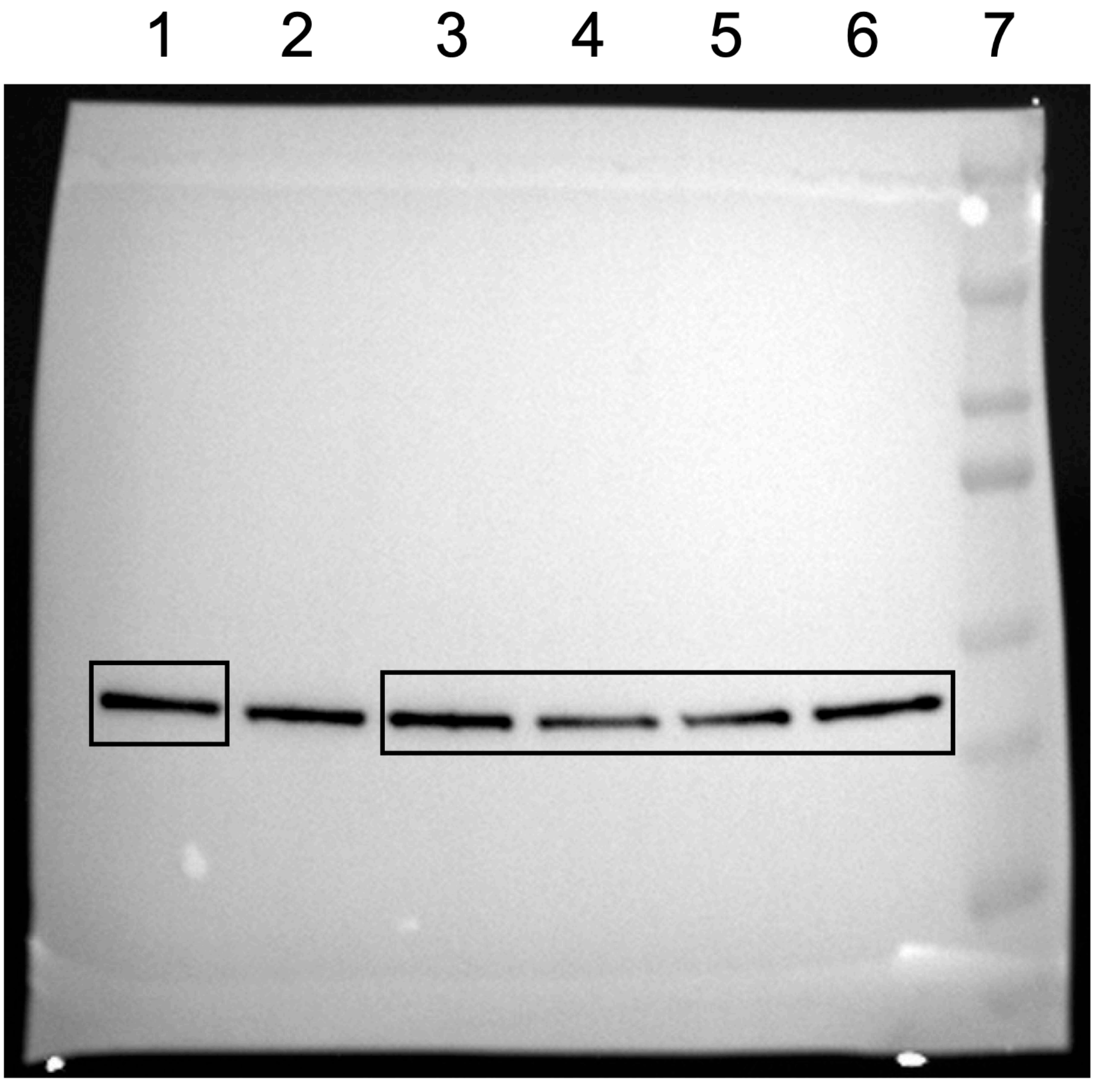


**Uncropped western blot for $\beta$-actin control for total eIF2α in Supplementary Figure S15.** Western blot of $\beta$-actin for total eIF2α in HEK293T cells 24 hours post-lipofection. 1) HEK293T without mRNA 2) Thapsigargin positive control 3) Regular mRNA 4) Unfolded-OG 5) Folded-OG 6) Folded mRNA-OG synthesized with 100% substitution of uridine for $m^1\Psi$ 7) Marker
